# CSF1R+ macrophage and osteoclast depletion impairs neural crest proliferation and craniofacial morphogenesis

**DOI:** 10.1101/2025.10.08.681206

**Authors:** Felix Ma, Rose Ru Jing Zhou, Matthew Rosin, Iris Zhou, Sabrina Ownsworth, Rouzbeh Ostadsharif Memar, Vincent B. Wong, Jessica M. Rosin

**Author notes:** Authors contributed equally (Iris Zhou, Sabrina Ownsworth).

## Abstract

Despite a wealth of knowledge on the mechanisms underlying craniofacial morphogenesis during gestation, the roles of fetal macrophages and osteoclasts during this process remain less well characterized. Here, we used the pharmacological inhibitor PLX5622 to disrupt colony stimulating factor-1 receptor (CSF1R) signaling, which is essential for macrophage and osteoclast proliferation, differentiation, and survival. Prenatal PLX5622 exposure resulted in ∼50% depletion of CSF1R+ macrophages, with complete loss of osteoclasts. While there were no notable changes in craniofacial nerve or muscle development, prenatal exposure to PLX5622 resulted in skull doming and cranial suture impairments, in addition to disruptions to development of the premaxilla, mandible, ear ossicles, palate, and cranial base. In response to PLX5622 exposure, cytokine and chemokine signaling was altered and neural crest proliferation was impaired. Our data also highlight sex– and strain-specific differences in PLX5622 phenotypes and together demonstrate that CSF1R+ macrophages and osteoclasts are essential for craniofacial morphogenesis.

## INTRODUCTION

Colony stimulating factor-1 receptor (CSF1R) is expressed in cells of the mononuclear phagocyte lineage—namely microglia, macrophages, and osteoclasts—and is required for their proliferation, differentiation, and survival^1–6^. CSF1R binds to two ligands, colony stimulating factor-1 (CSF1) and interleukin-34 (IL-34), which stimulate proliferative and differentiating signals via STAT, AKT, and ERK1/2 pathways^2,3,7^. CSF1R is essential for phagocytic and bone resorptive functions in macrophages and osteoclasts, respectively^6,8^. Deficient CSF1R signaling in rodents has been shown to cause an osteopetrotic phenotype, where a lack of CSF1R+ cells leads to disruption of bone marrow cavities, loss of bone marrow hematopoiesis, and increased bone density^9–12^. The *osteopetrotic* (*op*) phenotype has been characterized in *Csf1^op/op^* ligand mutant mice and *Csf1r* knockout (KO) mice^9,10,13,14^. These mice have reduced litter sizes and early postnatal lethality (up to 100%) depending on the strain^9,10,14^. Phenotypes observed in both *Csf1/Csf1r*-disrupted mouse models also include a domed skull, shortened snout, and toothlessness, while an osteosclerotic otic capsule, incus, and stapes have only been observed in *Csf1^op/op^* mice^10,15,16^. At the cellular level, these mice are deficient in blood monocytes and macrophages in most tissues; microglia are depleted in the receptor knockout but unaffected in the ligand mutant^9,14,17–19^. The *Csf1 toothless* (*tl*) rat and *Csf1r* KO rat closely replicate the mouse phenotypes, including osteopetrosis, doming of the skull, a shortened snout, toothlessness, and microglia and macrophage depletion across most tissues^11,12^. Interestingly, *Csf1r* KO rats also present with additional phenotypes such as hypomineralization of the calvaria, impairment of the cranial sutures, bulging eyes, and sclerosis of the cranial base^11,20–22^. Viability is markedly improved in rat models, with only 23% of males dying postnatally and all females surviving beyond weaning^11^. In humans, bi-allelic *CSF1R* mutations can cause bone phenotypes such as osteosclerosis of the cranial vault and cranial base, in addition to narrowing of the optic canal^23,24^.

To study the role of CSF1R+ cells in mediating the above-mentioned phenotypes, we previously reported a pharmacological inhibition model for disrupting these cells in a temporally-controlled manner, focusing on embryogenesis^25,26^. The CSF1R inhibitor PLX5622 was fed via chow to pregnant CD1 mice from embryonic day 3.5 (E3.5) onwards to expose mouse embryos to the inhibitor^25^. Indeed, like the severe depletion of microglia in the *Csf1r* KO mouse, treatment with PLX5622 depleted ∼99% of microglia in the hypothalamus by E15.5, demonstrating excellent oral bioavailability, penetrance, and efficacy of this model^25^. Outside of the brain, PLX5622 exposure was found to cause a noticeable reduction in CSF1R+ cells throughout craniofacial tissues at E18.5^26^. As in the mutant mouse models described, PLX5622 treatment also reduced litter sizes and increased postnatal lethality (∼23%)^25^. These CD1 mice also presented with domed skulls in addition to morphological disruptions to the incisors and molars; although, the continuous eruption of incisors did not appear to be inhibited^25^. Using Geometric Morphometric (GM) analyses, craniofacial and mandibular shape and size were shown to be significantly changed and decreased, respectively, by PLX5622 exposure^26^. Interestingly, a recent study showed that intragastric administration of PLX5622 to pregnant ICR mice between E14.5-17.5 reduced osteoclasts and resulted in an enlarged cartilaginous zone in the midpalatal suture^27^. However, previous studies did not fully characterize the depletion of CSF1R+ cells outside of the brain or across embryogenesis, nor did they thoroughly describe all craniofacial phenotypes or identify the underlying morphogenetic disruptions leading to the phenotypes observed using this pharmacological inhibition model.

In this study, we used our previously established pharmacological inhibition model to study the impact of prenatal PLX5622 exposure on craniofacial nerve, muscle, cartilage, and bone development across embryogenesis in two strains (CD1 & C57BL/6) and both sexes (males & females). Exposure to PLX5622 resulted in a continuous and stable depletion of CSF1R+ macrophages to ∼50% across gestation, in addition to the complete loss of osteoclasts. Although we did not observe overt changes in craniofacial nerve or muscle development across embryogenesis, disrupted bone phenotypes in craniofacial tissue of PLX5622-exposed animals could be observed as early as E15.5, with notable abnormalities in skull shape, cranial sutures, ear ossicles, premaxilla, mandible, palate, and cranial base observed by birth. We identified significant alterations in the secretion of 16 cytokines and chemokines from cultured craniofacial tissues in response to prenatal PLX5622 exposure and validated that several such secreted factors are indeed expressed by CSF1R+ macrophages. At the cellular level, exposure to PLX5622 across gestation decreased neural crest cell (NCC) proliferation both *in vitro* in a sphere assay and *in vivo* in embryonic craniofacial tissue cryosections, resulting in fewer NCCs in tissues impacted by PLX5622 exposure. Together, these data demonstrate that CSF1R+ macrophages and osteoclasts are required for craniofacial morphogenesis during the embryonic period and suggest that altered signaling to NCCs may underlie the phenotypes observed in PLX5622 offspring.

## RESULTS

### Embryonic Exposure to the CSF1R Inhibitor PLX5622 Depletes CSF1R+ Macrophages and Osteoclasts

As CSF1R function is essential for macrophage and osteoclast proliferation, differentiation, and survival^5,6^, we tested the impact of PLX5622 on reducing macrophage and osteoclast populations in craniofacial tissues across embryogenesis. Pregnant dams were fed control or PLX5622 diet starting at E3.5 to avoid disrupting embryo implantation—a process regulated by endometrial macrophages^28,29^, which was supported by our findings that administering PLX5622 gestationally did not significantly impact embryonic litter size (Table S1; CD1 p=0.0698, C57BL/6 p=0.5683), the number of embryo resorptions per litter (Table S1; CD1 p=0.6110, C57BL/6 p=0.4176), or embryo sex ratios (Table S1; CD1 p=0.5676, C57BL/6 p=0.4612) in CD1 or C57BL/6 mice. Craniofacial tissues were collected from *Csf1r^EGFP^* transgenic embryos, enabling CSF1R-expressing cells to be assessed by enhanced green fluorescent protein (EGFP) signal using flow cytometry (Figure 1A), which revealed a significant decrease in *Csf1r^EGFP+^* cells in response to PLX5622 exposure across all time-points assessed (Figure 1B; main effect of diet: E11.5 F_(1,18)_=24.42, p=0.0001; E13.5 F_(1,24)_=29.53, p<0.0001; E15.5 F_(1,17)_=27.25, p<0.0001; E17.5 F_(1,16)_=29.45, p<0.0001). Sustained depletion (40-63%) of *Csf1r^EGFP+^* cells was observed across gestation in PLX5622 male embryos, while a significant reduction in *Csf1r^EGFP+^* cells was only achieved at E17.5 for PLX5622 female embryos (Figure 1B; E11.5 male p=0.0009, female p=0.1321; E13.5 male p<0.0001, female p=0.0806; E15.5 male p<0.0001, female p=0.2610; E17.5 male p=0.0452, female p=0.0011). Interestingly, significantly more *Csf1r^EGFP+^* cells were found in control males as compared to females at E11.5 (Figure 1B; significant impact of sex F_(1,18)_=5.801, p=0.0270; control male vs. female p=0.0259).

**Figure 1.**
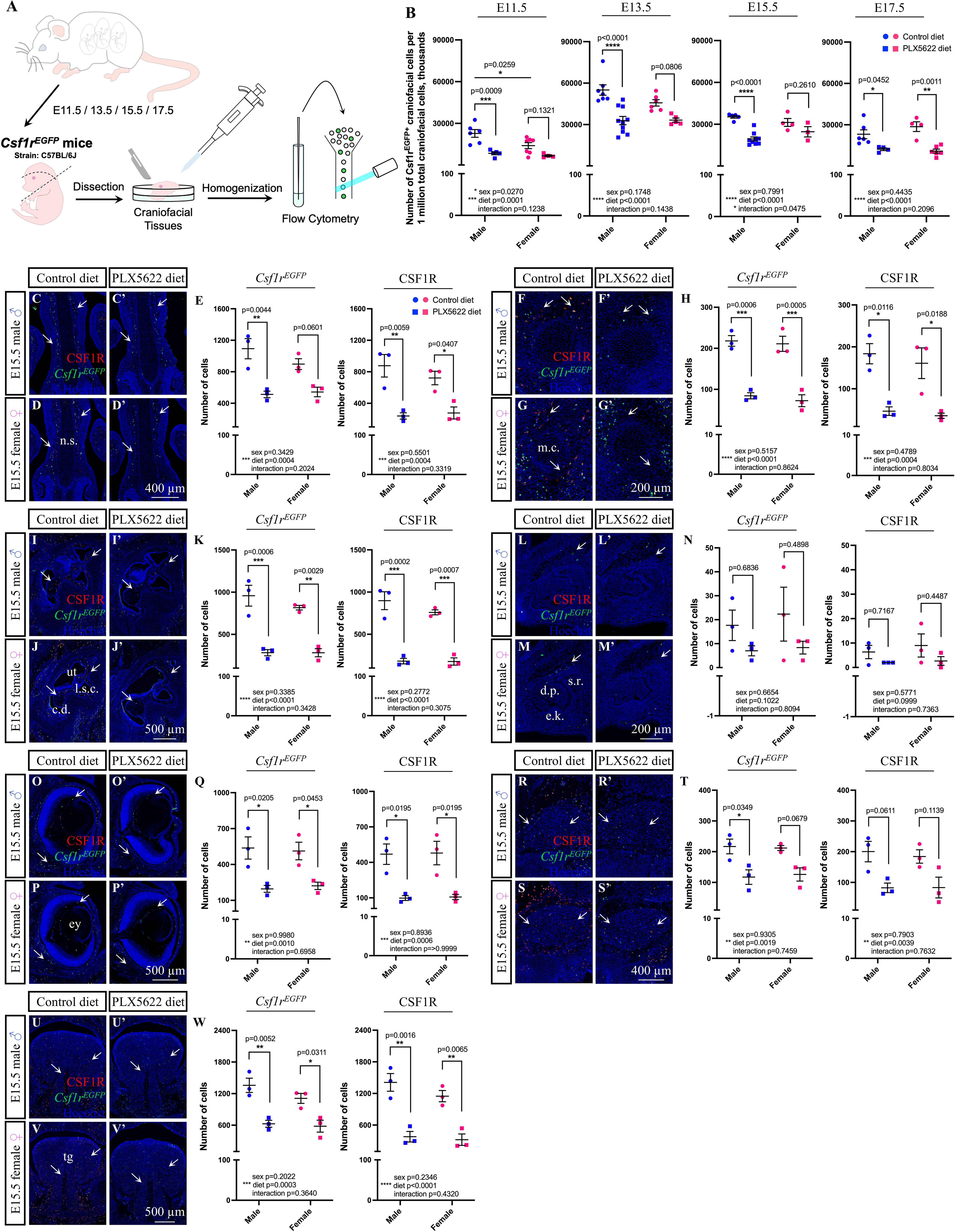
Gestational Exposure to PLX5622 Significantly Depletes CSF1R-Expressing Cells. (A) Schematic illustrating collection of *Csf1r^EGFP^*^+^ cells for flow cytometry. (B) Quantification of EGFP+ cells from *Csf1r^EGFP^* craniofacial tissues (N=3-10 embryos per sex/treatment/time-point from 2-4 dams). (C-W) Immunofluorescent images and quantification of CSF1R+ and *Csf1r^EGFP+^* cells in and around the E15.5 nasal septum (n.s.; C-E), Meckel’s cartilage (m.c.; F-H), ear (I-K), maxillary incisor (L-N), eye (ey; O-Q), trigeminal (R-T), and tongue (tg; U-W). Arrows mark CSF1R and *Csf1r^EGFP^* double-positive cells (N=3 embryos per sex/treatment from 2-3 dams). Abbreviations: c.d., cochlear duct; d.p., dental papilla; e.k., enamel knot; l.s.c., lateral semicircular canal; s.r., stellate reticulum; ut, utricle. Blue dots represent male and pink dots represent female. Counts represent mean ± SEM and were analyzed by a two-way ANOVA with Tukey’s post-hoc test.

To visualize and quantify tissue-specific depletion of CSF1R-expressing cells within the embryo, craniofacial tissue cryosections were collected from E15.5 male and female *Csf1r^EGFP^* transgenic embryos and stained with a CSF1R antibody, as only ∼92% of *Csf1r^EGFP+^* cells were found to also express CSF1R (data not shown). Quantification of CSF1R-expressing cells in and around several developing craniofacial structures of E15.5 PLX5622 embryos as compared to controls revealed a significant decrease of both *Csf1r^EGFP+^*(Figures 1C-1W; main effect of diet: nasal septum F_(1,8)_=32.85, p=0.0004; Meckel’s cartilage F_(1,8)_=95.02, p<0.0001; ear F_(1,8)_=74.37, p<0.0001; eye F_(1,8)_=25.33, p=0.0010; trigeminal F_(1,8)_=20.77, p=0.0019; tongue F_(1,8)_=35.67, p=0.0003) and CSF1R+ (Figures 1C-1W; main effect of diet: nasal septum F_(1,8)_=33.27, p=0.0004; Meckel’s cartilage F_(1,8)_=33.49, p=0.0004; ear F_(1,8)_=112.6, p<0.0001; eye F_(1,8)_=30.17, p=0.0006; trigeminal F_(1,8)_=16.12, p=0.0039; tongue F_(1,8)_=56.38, p<0.0001) cells in all tissues examined except for the maxillary incisor (Figure 1N). Specifically, CSF1R-expressing cells were significantly depleted in and around developing cartilaginous and/or bony structures of PLX5622 embryos, with only 53.7% of *Csf1r^EGFP+^* and 32.9% of CSF1R+ cells remaining around the nasal septum (Figures 1C–1E; *Csf1r^EGFP^* male p=0.0044, female p=0.0601; CSF1R male p=0.0059, female p=0.0407), 36.5% of *Csf1r^EGFP+^* and 23.8% of CSF1R+ cells remaining around Meckel’s cartilage (Figures 1F–1H; *Csf1r^EGFP^* male p=0.0006, female p=0.0005; CSF1R male p=0.0116, female p=0.0188), and 32.1% of *Csf1r^EGFP+^*and 21.8% of CSF1R+ cells remaining in the ear (Figures 1I–1K; *Csf1r^EGFP^* male p=0.0006, female p=0.0029; CSF1R male p=0.0002, female p=0.0007). CSF1R-expressing cells were also significantly reduced in craniofacial nervous tissue of PLX5622 embryos, with only 39.8% of *Csf1r^EGFP+^* and 21.6% of CSF1R+ cells remaining in the eye (Figures 1O–1Q; *Csf1r^EGFP^* male p=0.0205, female p=0.0453; CSF1R male p=0.0195, female p=0.0195), and 56.7% of *Csf1r^EGFP+^* cells remaining in the trigeminal; with only male embryos displaying a significant change in the trigeminal (Figures 1R–1T; *Csf1r^EGFP^* male p=0.0349, female p=0.0679). Moreover, CSF1R-expressing cells were also significantly reduced in muscle tissue, with only 49.2% of *Csf1r^EGFP+^* and 27.5% of CSF1R+ cells remaining in the tongue of E15.5 PLX5622 embryos as compared to controls (Figures 1U–1W; *Csf1r^EGFP^* male p=0.0052, female p=0.0311; CSF1R male p=0.0016, female p=0.0065).

While visualizing E15.5 *Csf1r^EGFP^* craniofacial tissue sections using high-resolution confocal microscopy, we observed multinucleated *Csf1r^EGFP^*^+^ cells within several craniofacial bones such as the premaxilla, maxilla, and mandible (Figures 2A–2F’, arrows). To quantify osteoclast-specific depletion following gestational exposure to PLX5622, we labelled osteoclasts with a Cathepsin K (CTSK) antibody. Intriguingly, exposure to PLX5622 completely depleted CTSK+/*Csf1r^EGFP+^* double-positive multinucleated osteoclasts in the E15.5 male and female mandible (Figures 2E–2I, arrows; main effect of diet F_(1,8)_=49.85, p=0.0001; male p=0.0009, female p=0.0052). Moreover, there was a complete loss of all single-positive *Csf1r^EGFP+^* multinucleated cells in the E15.5 PLX5622 mandible as compared to control for both sexes (Figures 2E–2I’; main effect of diet F_(1,8)_=49.85, p=0.0001; male p=0.0009, female p=0.0052). As embryonic osteoclasts appeared to be absent in the E15.5 mandible of PLX5622 embryos when compared to controls, we then looked to determine whether this led to loss of bone resorptive function across the embryonic head. Tartrate-resistant acid phosphatase (TRAP) staining of E15.5 craniofacial tissues revealed loss of osteoclastic activity surrounding developing craniofacial bones, including the premaxilla, mandible, maxilla, frontal, and basioccipital bones (Figures 2J–2S’, arrows). Moreover, quantification of the TRAP+ area in the premaxilla, mandible, and maxilla revealed a significant reduction in TRAP staining in both E15.5 male and female embryos in response to PLX5622 exposure (Figures 2T-2V; main effect of diet: premaxilla F_(1,8)_=101.0, p<0.0001; male p=0.0004, female p=0.0006; mandible F_(1,8)_=65.09, p<0.0001; male p=0.0016, female p=0.0027; maxilla F_(1,8)_=94.02, p<0.0001; male p=0.0004, female p=0.0010). Together, the data demonstrate that exposure to the CSF1R inhibitor PLX5622 during gestation significantly depletes CSF1R-expressing cells across embryogenesis and disrupts osteoclast development and activity.

**Figure 2.**
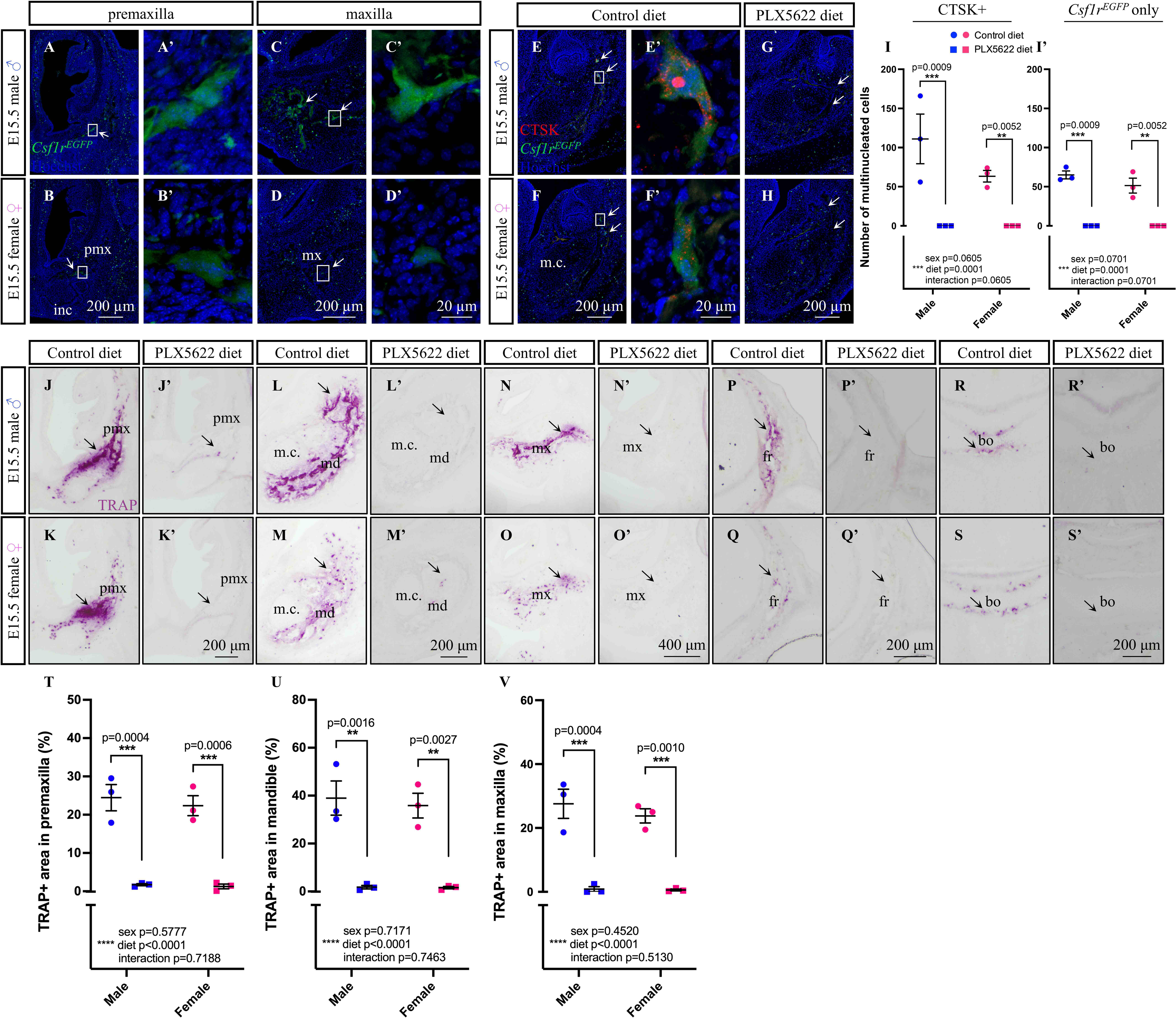
Prenatal Exposure to PLX5622 Disrupts Osteoclast Development and Function. (A-D’) *Csf1r^EGFP^*^+^ osteoclasts (arrows) in the E15.5 premaxilla (A-B’) and maxilla (C-D’). (E-I) Immunofluorescent images (E-H) and quantification (I-I’) of multinucleated CTSK/*Csf1r^EGFP^* double-positive osteoclasts (arrows; I) and *Csf1r^EGFP+^*single-positive osteoclasts (I’) in the E15.5 mandible. (J-S’) TRAP staining (arrows) in E15.5 premaxilla (pmx; J-K’), mandible (md; L-M’), maxilla (mx; N-O’), frontal (fr; P-Q’), and basioccipital (bo; R-S’) bones. (T-V) Quantification of total TRAP+ area in premaxilla (T), mandible (U), and maxilla (V). N=3 embryos per sex/treatment from 2-3 dams. Abbreviations: m.c, Meckel’s cartilage. Counts represent mean ± SEM and were analyzed by an ART ANOVA (I-I’) or a two-way ANOVA (T-V) with Tukey’s post-hoc test.

### CSF1R+ Cell Depletion Results in Tissue-Specific Changes in Apoptosis in the Developing Embryo

Considering that one of the main functions of macrophages is to phagocytose apoptotic cells and cellular debris^30,31^, and apoptosis is an essential process in embryonic craniofacial development^32–35^, craniofacial tissue cryosections collected from E15.5 *Csf1r^EGFP^* transgenic embryos were stained with an active cleaved Caspase 3 (CC3) antibody to visualize whether apoptotic cells accumulated following CSF1R+ cell depletion. We were also interested in whether we would observe CC3+/*Csf1r^EGFP+^* double-positive cells, as the reduction in CSF1R+ cells should be mediated in part by cell death. Quantification of CC3+ single-positive and CC3+/*Csf1r^EGFP+^* double-positive cells in and around several developing craniofacial structures of E15.5 embryos (Figures S1A-U’, arrows) revealed a significant increase in apoptosis in the ear (Figures S1I, F_(1,8)_=12.06, p=0.0084; S1I’, F_(1,8)_=7.043, p=0.0291), eye (Figures S1O, F_(1,8)_=9.904, p=0.0137; S1O’, F_(1,8)_=6.601, p=0.0392), trigeminal (Figure S1R, F_(1,8)_=16.18, p=0.0038), and tongue (Figure S1U, F_(1,8)_=332.0, p<0.0001) in response to PLX5622 exposure; while no changes in apoptosis were observed in and around the nasal septum (Figure S1C-C’), Meckel’s cartilage (Figure S1F-F’) or the maxillary incisor (Figure S1L-L’). A significant increase in CC3+ single-positive cells was only observed in the E15.5 PLX5622 female ear (Figures S1I; male p=0.8656, female p=0.0138); while no change in CC3+/*Csf1r^EGFP+^* double-positive cells was observed in either sex (Figure S1I’; male p=0.3976, female p=0.2363). Sex differences were also identified in the ear (Figure S1I; significant impact of sex F_(1,8)_=5.68, p=0.0443 and sex x diet F_(1,8)_=5.68, p=0.0443), whereby significantly more CC3+ cells were observed in PLX5622 females as compared to males (Figure S1I; p=0.0395). Surprisingly, we did not observe a statistically significant change in CC3+ single– (Figure S1O; male p=0.2945, female p=0.1279) or CC3+/*Csf1r^EGFP+^* double-positive (Figure S1O’; male p=0.2944, female p=0.4463) cells in the eye for either sex. In contrast, CC3+ cells showed a significant increase in the trigeminal of PLX5622 male embryos but not females (Figure S1R; male p=0.0216, female p=0.3065). Intriguingly, the tongue was the only craniofacial structure assessed that displayed a significant increase in CC3+ cells in both PLX5622 male and female embryos (Figure S1U; male p<0.0001, female p<0.0001), in addition to sex differences (Figure S1U; significant impact of sex F_(1,8)_=7.364, p=0.0265), where there were significantly more CC3+ cells observed in PLX5622 males as compared to females (Figure S1U; p=0.0406). Collectively, the data suggests that depleting CSF1R+ cells during embryogenesis leads to an accumulation of apoptotic cells predominantly in nervous and muscular structures, but not in or around developing cartilaginous/bony structures, except for the ear.

### Embryonic Exposure to the CSF1R Inhibitor PLX5622 Does Not Impact Craniofacial Nerve or Muscle Development

In this study, we sought to thoroughly characterize the craniofacial phenotypes resulting from embryonic PLX5622 exposure, especially as embryonic soft tissues were previously unexplored in *Csf1*/*Csf1r*-disrupted genetic models^9–12,14,15,21^ and our own pharmacological model using PLX5622^25,26^. Accordingly, we first examined craniofacial nerve development from E11.5 to E13.5 using a Neurofilament antibody (2H3) and whole-mount immunostaining (Figures S2A–S2L). Neurofilament staining in the trigeminal and facial nerves, including the zygomatic, buccal, and marginal branches of the trigeminal, were comparable between E11.5 control and PLX5622 male and female CD1 (Figures S2A–S2D) and C57BL/6 (data not shown) embryos. Similarly, neurofilament staining in the facial nerve, auriculotemporal nerve, and great auricular nerve were comparable between E12.5 control and PLX5622 male and female CD1 (Figures S2E–S2H) and C57BL/6 (data not shown) embryos. Moreover, craniofacial nerve development continued to be comparable at E13.5, whereby the great auricular nerve, auriculotemporal nerve, zygomatic branch of the trigeminal, superior buccolabial nerve, inferior buccolabial nerve, and marginal mandibular nerve appeared similar between control and PLX5622 male and female CD1 (Figures S2I–S2L) and C57BL/6 (data not shown) embryos.

Next, we examined craniofacial muscle development from E11.5 to E13.5 using a Myosin Heavy Chain antibody (MF 20) and whole-mount immunostaining (Figures S2M–S2X). Muscle staining in the masseter, auricularis, buccinator, and zygomaticomandibularis, appeared comparable across E11.5 to E13.5 in control and PLX5622 male and female CD1 embryos (Figures S2M–S2X). To assess the development of muscles deeper in the tissues, immunofluorescence staining was performed on E15.5 craniofacial tissue cryosections using the MF20 antibody (Figure S3). Muscle fiber diameter and density appeared comparable in the buccinator, zygomaticomandibularis, deep masseter, superficial masseter, temporalis, pterygoid, tongue, and extraocular muscles, including the superior oblique, inferior oblique, superior rectus, medial rectus, inferior rectus, and retractor bulbi between E15.5 control and PLX5622 male and female C57BL/6 embryos (Figures S3A–S3L’). These data suggests that depleting CSF1R+ cells during embryogenesis does not noticeably impact the gross morphology of craniofacial nerves or muscles in male or female CD1 or C57BL/6 embryos.

### Prenatal Exposure to the CSF1R Inhibitor PLX5622 Directs Tissue and Time-Dependent Changes in Craniofacial Morphogenesis During Embryogenesis

Previously, we reported doming of the cranial vault, mandibular disruptions, and dental phenotypes in P21 and P28 CD1 mice exposed to PLX5622 embryonically^25,26^. However, our prior analyses did not fully characterize all the cartilaginous and/or bony phenotypes present in PLX5622 offspring. Moreover, the *Csf1*/*Csf1r*-disrupted genetic models and our own pharmacological PLX5622 studies failed to examine craniofacial morphogenesis during the embryonic period^9–12,15,16,21,25,26^. As we observed a loss of osteoclasts and bone resorptive activity in E15.5 PLX5622 embryos in the current study (see Figure 2), we utilized E15.5 craniofacial tissue cryosections to visualize the impact of PLX5622 exposure on osteoblasts and bone formation during embryogenesis. To start, immunofluorescence staining with an Sp7 antibody was performed to label osteoblasts. Disrupted patterning in the premaxilla and mandible (Figures 3A–3D’, arrows) was observed in E15.5 male and female PLX5622 embryos, while other craniofacial bones such as the maxilla, frontal bone, and basioccipital bone displayed comparable Sp7 staining (Figures 3E–3J’).

**Figure 3.**
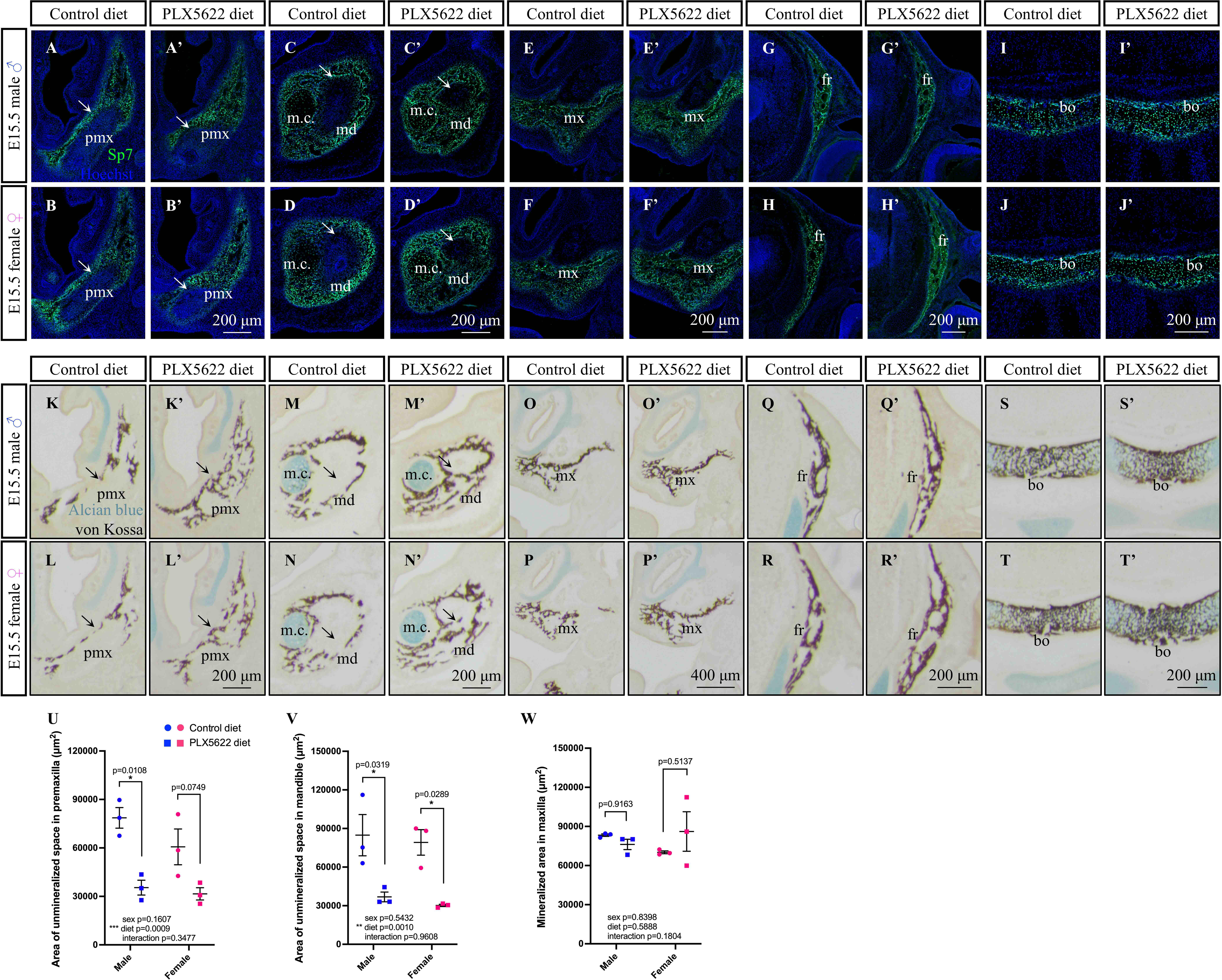
Exposure to PLX5622 During Embryogenesis Increases Bone Density in the Premaxilla and Mandible. (A-J’) E15.5 Sp7 osteoblast staining in the premaxilla (pmx; A-B’) mandible (md; C-D’), maxilla (mx; E-F’), frontal (fr; G-H’), and basioccipital (bo; I-J’) bones of CD1 embryos. (K-T’) E15.5 von Kossa and Alcian blue staining in the premaxilla (K-L’), mandible (M-N’), maxilla (O-P’), frontal (Q-R’), and basioccipital (S-T’) bones of CD1 embryos. (U-W) Quantification of unmineralized area within the premaxilla (U) and mandible (V), and mineralized area in the maxilla (W). N=3 embryos per sex/treatment from 2-3 dams. Abbreviations: m.c, Meckel’s cartilage. Measurements represent mean ± SEM and were analyzed by a two-way ANOVA with Tukey’s post-hoc test.

Next, we evaluated cartilage and bone development using Alcian blue and von Kossa staining, respectively. Craniofacial cartilage development appeared unimpacted, as Alcian blue staining was comparable between E15.5 control and PLX5622 male and female embryos (Figures 3K–3T’). In contrast, increased bone density in the premaxilla and mandible (Figures 3K–3N’, arrows) was observed in E15.5 male and female PLX5622 embryos, which complemented the altered osteoblast patterning observed in these bones. Moreover, consistent with Sp7 osteoblast staining, bone density in other craniofacial bones such as the maxilla, frontal bone, and basioccipital bone (Figures 3O–3T’) appeared comparable between E15.5 control and PLX5622 male and female embryos. Quantification of unmineralized areas in the premaxilla and mandible, which appeared to fill in with von Kossa staining in E15.5 PLX5622 embryos (Figures 3K-N’, arrows), was shown to significantly decrease in response to PLX5622 exposure (Figures 3U-3V; main effect of diet: premaxilla F_(1,8)_=26.23, p=0.0009; mandible F_(1,8)_=25.34, p=0.0010). Although the unmineralized area significantly decreased in the mandible of both sexes (Figure 3V; male p=0.0319, female p=0.0289), it was only found to be statistically significant in the premaxilla of E15.5 male embryos (Figure 3U; male p=0.0108, female p=0.0749). Importantly, quantification of mineralization in the maxilla, a region that displayed comparable von Kossa staining between E15.5 control and PLX5622 embryos (Figures 3O-3P’), showed no change in mineralization in response to PLX5622 exposure (Figure 3W).

To investigate if the changes in osteoblast/bone development in PLX5622 embryos begin prior to E15.5, micromass cultures of mesenchymal cells collected from the frontonasal mass (gives rise to premaxilla)^36–38^ and mandible of E12.5 control and PLX5622 male and female embryos were stained for alkaline phosphatase activity as a readout for bone mineralization. Relative to total micromass area, which was stained with hematoxylin, control and PLX5622 frontonasal mass-and mandible-derived micromass cultures did not show a significant difference in alkaline phosphatase staining (Figures S4A–S4L). Combined, these data demonstrate that prenatal exposure to PLX5622 drives region-specific disruptions to the development of the premaxilla and mandible between E12.5 and E15.5.

### Embryonic Exposure to the CSF1R Inhibitor PLX5622 Drives Sex and Strain-Dependent Disruptions to Craniofacial Bone Morphogenesis

Given our findings suggest a possible increase in osteoblast differentiation and/or osteogenic potential in the premaxilla and mandible of PLX5622 embryos between E12.5 and E15.5, we were interested in whether other cartilage and/or bony phenotypes would arise late in embryogenesis. Accordingly, to visualize any phenotypes that develop late in embryogenesis, we stained P1 CD1 and C57BL/6 male and female skulls with Alcian blue and Alizarin red to label cartilage and bone, respectively. Consistent with previous P21 and P28 findings^25^, doming of the cranial vault was observed in all P1 PLX5622 CD1 male and female pups, with overt changes to calvarial bone size and density, including impairments to sagittal and coronal suture development (Figures 4A–4D’). Moreover, the length of the skull, interparietal bone, and occipital bone were all found to significantly decrease in P1 PLX5622 CD1 male and female pups as compared to control (Figures 4E-G; main effect of diet: skull F_(1,20)_=35.64, p<0.0001; male p=0.0023, female p=0.0021; interparietal bone F_(1,20)_=49.10, p<0.0001; male p=0.0021, female p<0.0001; occipital bone F_(1,20)_=44.85, p<0.0001; male p=0.0013, female p=0.0004). Similar, but milder, doming of the cranial vault was observed in P1 PLX5622 C57BL/6 male and female pups (Figures S5A–S5B’, arrows). However, disruption to calvarial bone size and density, including impairments to sagittal and interfrontal sutures (Figures S5C–S5D’), were still prominent in P1 PLX5622 C57BL/6 male and female pups as compared to control. In contrast, skull and interparietal bone length were not significantly impacted by PLX5622 exposure (Figure S5E-S5F); although occipital bone length did significantly decrease in P1 PLX5622 C57BL/6 male pups (Figure S5G; main effect of diet F_(1,20)_=13.22, p=0.0016; male p=0.0219, female p=0.2397).

**Figure 4.**
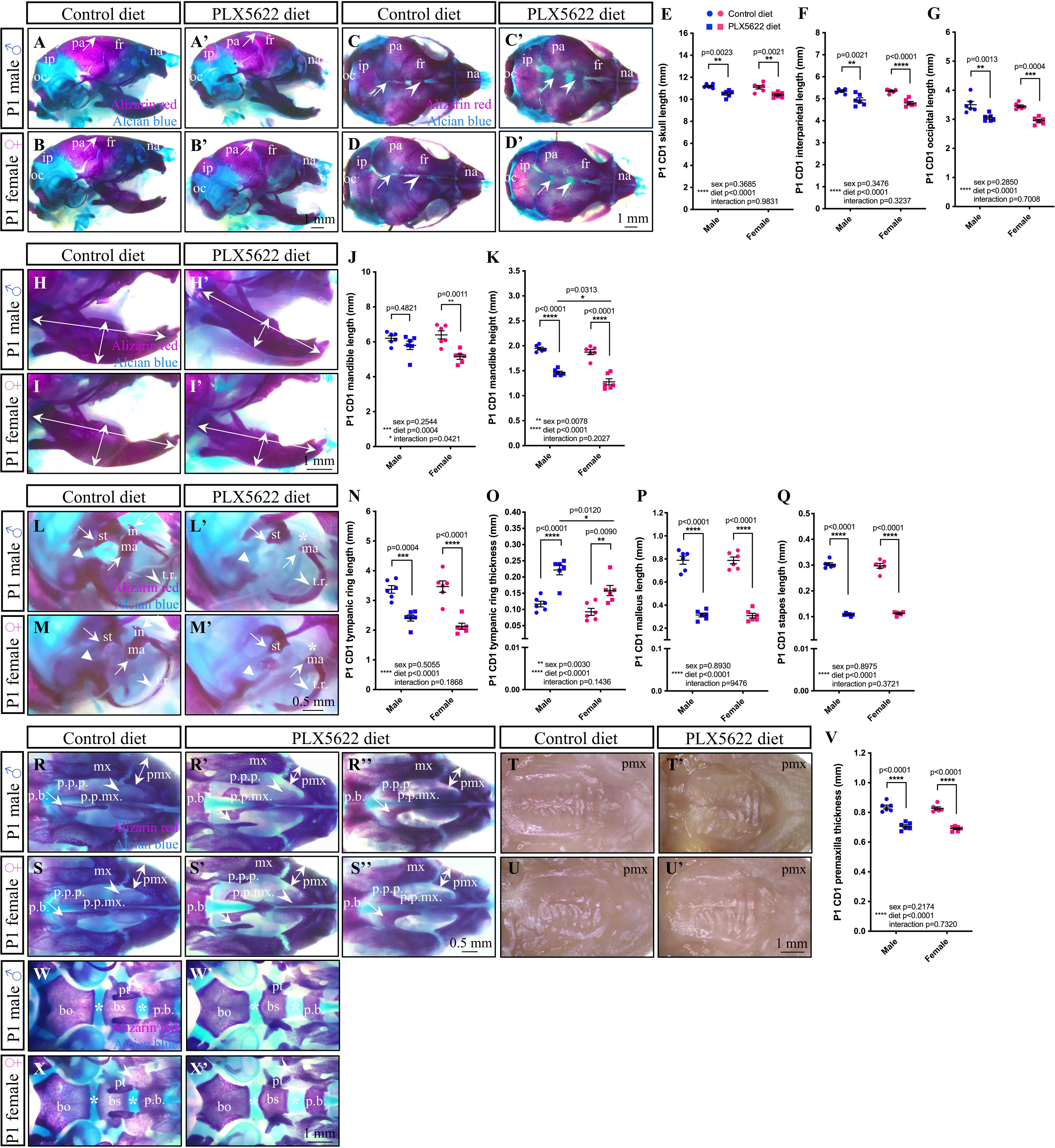
Prenatal Exposure to PLX5622 Disrupts Craniofacial Bone Morphogenesis. (A-D’) Lateral (A-B’) and dorsal (C-D’) views of P1 CD1 skull. (E-G) Quantification of skull (E), interparietal (ip; F), and occipital (oc; G) bone lengths. (H-K) Lateral view of P1 CD1 mandibles (H-I’) and quantification of mandible length (J) and height (K). (L-Q) Lateral view of the P1 CD1 ear (L-M’) and quantification of tympanic ring (t.r.) length (N) and thickness (O), and malleus (ma; P) and stapes (st; Q) lengths. Arrows mark incus (in; absent in PLX5622 pups, asterisks). (R-U’) Ventral views of the P1 CD1 palate. (V) Quantification of premaxilla (pmx) thickness. (W-X’) Ventral view of the P1 CD1 cranial base. N=5-8 pups per sex/treatment from 2-4 dams. Abbreviations: bo, basioccipital; bs, basisphenoid; fr, frontal bone; mx, maxilla; na, nasal bone; pa, parietal bone; p.b., palatine bone; p.p.mx., palatine process of the maxilla; p.p.p., palatine process of the palatine; pt, pterygoid process. Measurements represent mean ± SEM and were analyzed by a two-way ANOVA with Tukey’s post hoc test.

When focusing on the Pl CD1 mandible (Figures 4H–4I’), we began to observe notable sex differences in mandible length (Figure 4J; significant effect of diet F_(1,20)_=17.88, p=0.0004 and diet x sex interaction F_(1,20)_=4.717, p=0.0421) and height (Figure 4K; significant effect of diet F_(1,20)_=151.6, p<0.0001 and sex F_(1,20)_=8.754, p=0.0078) in response to embryonic PLX5622 exposure. Indeed, only P1 PLX5622 CD1 female pups had shorter mandibles (Figure 4J; male p=0.4821, female p=0.0011). While P1 PLX5622 CD1 mandible height significantly decreased in both sexes (Figure 4K; male p<0.0001, female p<0.0001), a greater disruption was observed in P1 PLX5622 CD1 female pups (Figure 4K; PLX5622 diet male vs. female p=0.0313). In contrast, P1 control and PLX5622 C57BL/6 male and female mandible length was comparable (Figures S5H–S5J), while mandible height was significantly decreased in both P1 PLX5622 C57BL/6 males and females alike (Figure S5K; main effect of diet F_(1,20)_=38.62, p<0.0001; male p<0.0001, female p=0.0233).

In assessing the P1 CD1 ear, we observed notable abnormalities to the ear ossicles, including an absence of the incus in PLX5622 male and female pups as compared to controls (Figures 4L–4M’). Bone density was disrupted in the otic capsule and tympanic ring (Figures 4L–4M’), where the tympanic ring was found to be significantly shorter in PLX5622 CD1 males and females alike (Figure 4N; main effect of diet F_(1,20)_=70.61, p<0.0001; male p=0.0004, female p<0.0001). Interestingly, we observed sex differences with respect to tympanic ring thickness (Figure 4O; significant effect of diet F_(1,20)_=43.62, p<0.0001 and sex F_(1,20)_=11.42, p=0.0030). While the P1 PLX5622 CD1 tympanic ring was significantly thicker in both sexes (Figure 4O; male p<0.0001, female p=0.0090), a larger disruption was observed in P1 PLX5622 CD1 male pups (Figure 4O; PLX5622 diet male vs. female p=0.0120). Prominent disruptions to the malleus and stapes were also observed when comparing P1 CD1 control and PLX5622 pups (Figures 4L–4M’), which included a significant decrease in both malleus (Figure 4P; main effect of diet F_(1,20)_=327.8, p<0.0001; male p<0.0001, female p<0.0001) and stapes (Figure 4Q; main effect of diet F_(1,20)_=1093, p<0.0001; male p<0.0001, female p<0.0001) length in both sexes in response to PLX5622 exposure. Interestingly, the P1 control C57BL/6 ear appeared less developed than what was observed at P1 in CD1 pups (compare Figures 4L–4M’ to Figures S5L–S5M’), as the incus was absent. Similar to the incus, the otic capsule appeared underdeveloped in the P1 control C57BL/6 ear as compared to what was observed in CD1 pups (compare Figures 4L–4M’ to Figures S5L–S5M’); yet otic capsule mineralization was still found to be disrupted in P1 PLX5622 C57BL/6 male and female pups when compared to control (Figures S5L–S5M’, triangles). Surprisingly, analysis of tympanic ring length (Figures S5L–S5M’, arrowheads) showed sex differences (Figure S5N; significant effect of diet F_(1,20)_=19.06, p=0.0003 and diet x sex interaction F_(1,20)_=4.939, p=0.0380), whereby P1 control C57BL/6 female tympanic ring length was found to be significantly shorter than male pups (Figure S5N; p=0.0490), and PLX5622 exposure was only found to impact tympanic ring length in P1 PLX5622 C57BL/6 male pups (Figure S5N; male p=0.0008, female p=0.4474). In contrast, tympanic ring thickness significantly increased in both P1 PLX5622 C57BL/6 males and females alike (Figure S5O; main effect of diet F_(1,20)_=97.98, p<0.0001; male p<0.0001, female p<0.0001).

Underdevelopment of the malleus and stapes was also observed when comparing P1 C57BL/6 control and PLX5622 pups (Figures S5L–S5M’), which appeared as a significant decrease in both malleus (Figure S5P; main effect of diet F_(1,20)_=27.87, p<0.0001) and stapes (Figure S5Q; main effect of diet F_(1,20)_=59.96, p<0.0001) length. While a significant decrease in malleus length was only observed in P1 C57BL/6 males (Figure S5P; male p=0.0008, female p=0.0505), stapes length significantly decreased in both sexes (Figure S5Q; male p<0.0001, female p=0.0029).

Intriguingly, P1 PLX5622 CD1 male and female pups also presented with disruptions to the palate, palatine sutures, and premaxilla when compared to controls (Figures 4R-4S’’); however, the palatine processes were only distinctly separated and sutures completely open in 50% of PLX5622 pups (Figures 4R’ and 4S’), compared to a milder phenotype seen in the remaining pups where the palatine processes were in contact but smaller (Figures 4R’’ and 4S’’). Considering that the palate phenotype was severe in some pups and reminiscent of cleft palate^39–42^, we examined unstained P1 PLX5622 CD1 palates to confirm that no overt clefts were present (Figures 4T–4U’). Importantly, quantification of premaxilla thickness in P1 CD1 pups demonstrated that embryonic exposure to PLX5622 significantly decreased the size of the premaxilla in both sexes (Figure 4V; main effect of diet F_(1,20)_=173.8, p<0.0001; male p<0.0001, female p<0.0001). Abnormalities in P1 PLX5622 CD1 male and female pup cranial base development, including wider spheno-occipital and intersphenoid synchondroses, and morphological alterations to the palatine-basisphenoid sutures, could also be observed (Figures 4W-4X’). Disruptions to the palate, palatine processes, and premaxilla appeared milder in Pl PLX5622 C57BL/6 male and female pups (Figures S5R–S5S’); however, quantification of premaxilla thickness in P1 C57BL/6 pups demonstrated that embryonic exposure to PLX5622 significantly decreased the size of the premaxilla in both sexes (Figure S5T; main effect of diet F_(1,20)_=371.9, p<0.0001; male p<0.0001, female p<0.0001). Similarly, subtle abnormalities were observed in the cranial base of Pl PLX5622 C57BL/6 male and female pups, including wider spheno-occipital and intersphenoid synchondroses, while the palatine bones and palatine sutures did not appear to be impacted (Figures S5U–S5V’). Taken together, these data suggest that CSF1R+ cells play important and widespread roles in craniofacial bone morphogenesis. Notably, we also observed sex differences, with male-specific phenotypes in the ear and female-specific phenotypes in the mandible, in addition to strain-dependent effects, with C57BL/6 pups presenting milder phenotypes overall.

### Skull Doming in PLX5622 Pups is Not Caused by Craniosynostosis

As doming of the cranial vault is often associated with midface hypoplasia and a shortened cranial base^43–48^, the cranial base of P1 control and PLX5622 CD1 male and female pups were also evaluated by micro-computed tomography (µCT; Figures 5A–5D). No disruptions to posterior cranial base length or anterior frontal complex length were observed between P1 control and PLX5622 CD1 male and female pups (Figures 5E-5F). Similarly, no significant differences were observed in the lengths of the basioccipital or basisphenoid bones when comparing P1 control and PLX5622 CD1 male and female pups (Figures 5G-5H); however, the presphenoid bone was significantly shorter in P1 PLX5622 CD1 male and female pups (Figure 5I; significant effect of diet F_(1,8)_=65.07, p<0.0001; male p=0.0003, female p=0.0203). Although there was a significant impact of PLX5622 on the size of the spheno-occipital synchondrosis (Figure 5J; F_(1,8)_=5.740, p=0.0435), assessment of P1 control and PLX5622 CD1 male and female pups did not show significance (Figure 5J; male p=0.3106, female p=0.4719). In contrast, P1 PLX5622 CD1 male and female pups presented with significantly wider inter-sphenoid synchondroses (Figure 5K; main effect of diet F_(1,8)_=31.50, p=0.0005; male p=0.0066, female p=0.0487).

**Figure 5.**
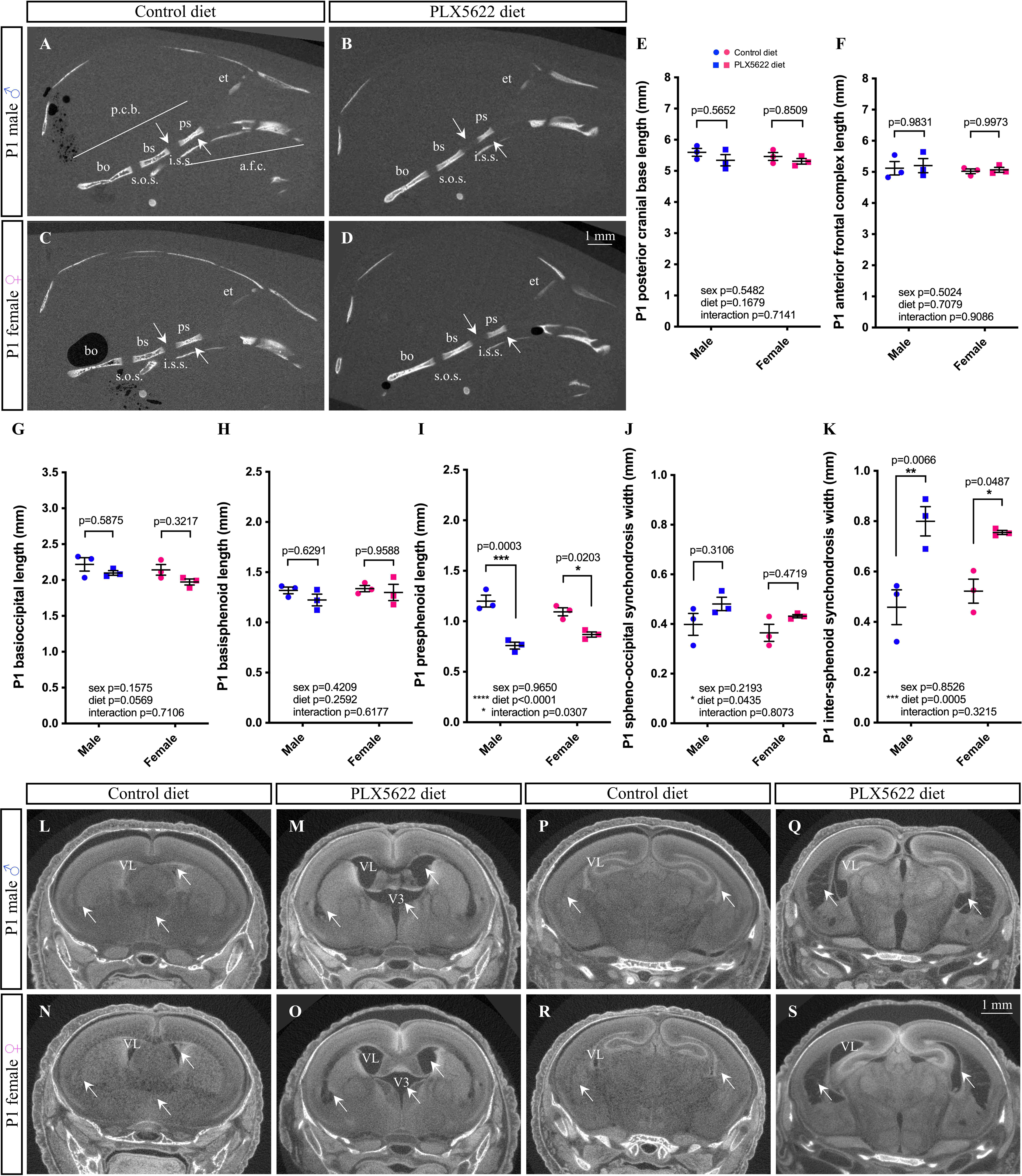
Prenatal Exposure to PLX5622 Disrupts Cranial Base and Brain Development. (A-D) Lateral view of μCT scans of P1 CD1 skulls show presphenoid bone (ps) and inter-sphenoid synchondrosis (i.s.s.). (E-K) Quantification of posterior cranial base (p.c.b.) length (E), anterior frontal complex (a.f.c.) length (F), basioccipital (bo) length (G), basisphenoid (bs) length (H), presphenoid length (I), spheno-occipital synchondrosis (s.o.s.) width (J), and inter-sphenoid synchondrosis (i.s.s.) width (K). (L-S) Coronal view of μCT scans of P1 CD1 skulls across the rostral (L-O) to caudal (P-S) axis. Arrows mark lateral (VL) and third ventricles (V3). N=3-5 pups per sex/treatment from 2-3 dams. Abbreviations: et, ethmoid. Measurements represent mean ± SEM and were analyzed by a two-way ANOVA with Tukey’s post-hoc test.

As hydrocephalus could also be contributing to skull doming^49–54^, contrast enhanced µCT was used to examine the gross morphology of P1 control and PLX5622 CD1 male and female pup brains (Figures 5L–5S). Ventriculomegaly of the lateral and third ventricles, in addition to bilateral cavitary lesions at the cortico-striato-amygdalar boundary, could be observed both rostrally (Figures 5L–5O, arrows) and caudally (Figures 5P–5S, arrows) when comparing P1 PLX5622 CD1 male and female pups to controls. Overall, while our data suggests that PLX5622 pups do not display craniosynostosis, exposure to PLX5622 during embryogenesis does drive disruptions to the cranial base and structural changes to the gross morphology of the brain.

### Gestational Exposure to PLX5622 Alters Cytokine and Chemokine Secretion from CSF1R+ Cells

Given that macrophages are known to secrete cytokines and chemokines, which are capable of stimulating tissue growth postnatally^21,55^ and during wound healing^56,57^, we next investigated whether PLX5622 exposure disrupts CSF1R+ cell-derived secreted factors to contribute to the reported craniofacial phenotypes. Luminex analysis of cytokines and chemokines secreted from cultured E13.5 control and PLX5622 C57BL/6 craniofacial tissues (Figure 6A) showed a significant downregulation of CCL2, CCL3, CCL4, CCL5, CCL12, CCL22, CXCL1, CXCL2, CXCL10, G-CSF, IFNβ-1, IL-6, and TNFα (Figures 6B–6G, 6I–O; main effect of diet: CCL2 F_(1,32)_=113.7, p<0.0001; CCL3 F_(1,32)_=68.72, p<0.0001; CCL4 F_(1,32)_=90.93, p<0.0001; CCL5 F_(1,32)_=18.84, p=0.0001; CCL12 F_(1,32)_=120.1, p<0.0001; CCL22 F_(1,32)_=46.38, p<0.0001; CXCL1 F_(1,32)_=20.25, p<0.0001; CXCL2 F_(1,32)_=117.2, p<0.0001; CXCL10 F_(1,32)_=59.47, p<0.0001; G-CSF F_(1,32)_=70.92, p<0.0001; IFNβ-1 F_(1,32)_=15.89, p=0.0004; IL-6 F_(1,32)_=71.26, p<0.0001; TNFα F_(1,32)_=137.4, p<0.0001) and a significant upregulation of CX3CL1 (Figure 6H; main effect of diet F_(1,32)_=30.35, p<0.0001) upon PLX5622 exposure. Interestingly, CCL2 (Figure 6B; male p<0.0001, female p<0.0001), CCL3 (Figure 6C; male p<0.0001, female p<0.0001), CCL4 (Figure 6D; male p<0.0001, female p<0.0001), CCL12 (Figure 6F; male p<0.0001, female p<0.0001), CCL22 (Figure 6G; male p=0.0001, female p=0.0003), CXCL2 (Figure 6J; male p<0.0001, female p<0.0001), CXCL10 (Figure 6K; male p<0.0001, female p<0.0001), G-CSF (Figure 6L; male p=0.0001, female p<0.0001), IL-6 (Figure 6N; male p=0.0003, female p<0.0001), and TNFα (Figure 6O; male p<0.0001, female p<0.0001) were downregulated in both E13.5 PLX5622 male and female craniofacial tissue cultures, while CCL5 (Figure 6E; male p=0.2950, female p=0.0007), CXCL1 (Figure 6I; male p=0.0838, female p=0.0025), and IFNβ-1 (Figure 6M; male p=0.2697, female p=0.0034) were only found to be significantly downregulated in E13.5 PLX5622 female craniofacial tissue cultures. In contrast, CX3CL1 was significantly upregulated in both sexes (Figure 6H; male p=0.0002, female p=0.0255). Although 34 other cytokines and chemokines were assessed in our analyses, these secreted factors were either lowly expressed, not significantly changed, or below the level of detection (Table S2).

**Figure 6.**
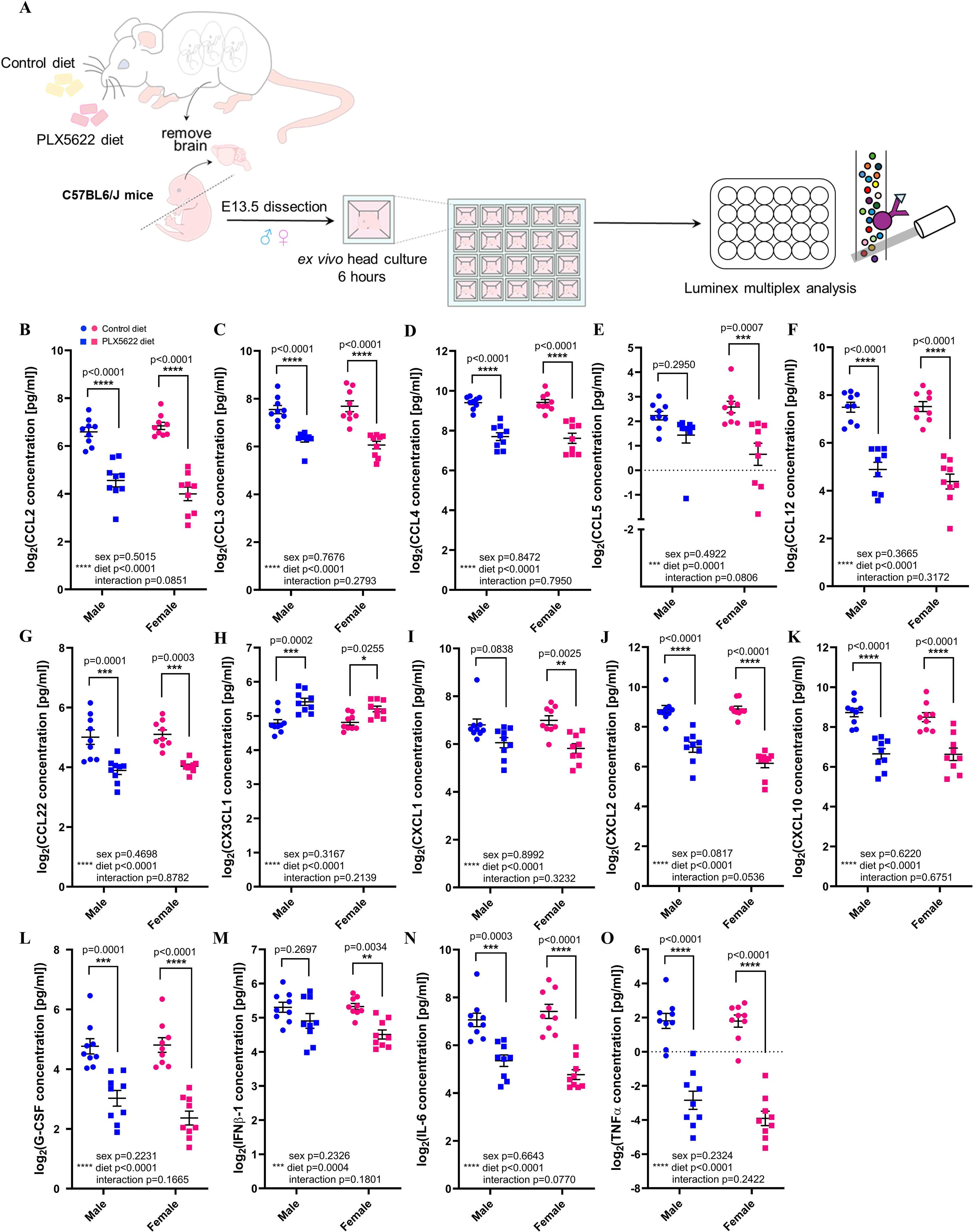
Exposure to PLX5622 During Embryogenesis Disrupts Cytokine and Chemokine Secretion in Craniofacial Tissue Cultures. (A) Schematic illustrating craniofacial tissue culture and Luminex multiplex analysis. (B-O) Cytokine/chemokine analysis of E13.5 craniofacial tissue culture media. N=9 per sex/treatment from 3 dams. Quantifications represent mean ± SEM and were analyzed by a two-way ANOVA with Tukey’s post hoc test.

To confirm that these cytokines and chemokines could indeed be derived from CSF1R+ cells in craniofacial tissues, we analyzed publicly available single-cell RNA sequencing datasets from E12.5 and E13.5 FVB/NJ and C57BL/6J craniofacial mesenchyme^58,59^. *Csf1r*-expressing cells extracted from these datasets formed three distinct clusters, with clusters 1 and 2 upregulating macrophage markers such as *Ptprc* and *Lyve1*, while cluster 3 expressed the osteoclast-specific marker *Ctsk* (Figure S6A). Cluster 1 was found to express *Ccl2, Ccl3, Ccl4,* and *Ccl12*, while cluster 2 only upregulated *Ccl12* (Figure S6A). Other cytokines and chemokines, such as *Cxcl2*, *Cxcl10*, *Tnf*, etc., were not found to be upregulated in any of the clusters (Figure S6A). Importantly, assessment of mRNA expression in the E15.5 mandible, maxilla, and ear (Figures S6B–S6D) using fluorescent *in situ* hybridization revealed the presence of *Csf1r*/*Ccl4* double-positive cells in each craniofacial tissue examined (Figures S6E–S6G). Moreover, immunofluorescence staining of E15.5 craniofacial tissue cryosections also confirmed the presence of CCL3 and CCL4 in *Csf1r^EGFP+^* cells in each of the craniofacial tissues assessed (Figures S6H–S6S).

Given that macrophages can themselves respond to secreted signals^60–78^, we were interested in whether treatment with the cytokines and chemokines found to be altered would impact macrophage physiology, such as phagocytosis (Figure S7A). Although incubation of E13.5 craniofacial tissue with a mixture of CCL2, CCL3, CCL4, CCL12, CXCL2, CXCL10, and IL-6 did not increase phagocytosis of pHrodo BioParticles by *Csf1r^EGFP+^* cells *ex vivo* (Figure S7B), we observed interactions between *Csf1r^EGFP+^* macrophages and *Wnt1^Cre^*-driven tdTomato+ NCCs across numerous embryonic craniofacial structures such as the premaxilla and mandible (Figures S7C–J), which has been observed in human embryonic skin^79^. Combined, these data suggest that embryonic exposure to PLX5622 significantly disrupts cytokine and chemokine levels in craniofacial tissues, with these changes likely being primarily driven by CSF1R+ cells and suggest that CSF1R+ cells may signal to and physically interact with NCCs to impact NCC developmental programs.

### CSF1R+ Cell Depletion Disrupts Neural Crest Proliferation

To test whether the depletion of CSF1R+ cells and altered cytokine/chemokine signaling impact surrounding cells to influence craniofacial morphogenesis, we focused on NCCs as many of the craniofacial structures disrupted by PLX5622 exposure are derived from this lineage. To start, we employed a neural crest-derived sphere assay to examine the impact of PLX5622 exposure on NCCs (Figure 7A). Briefly, neural crest-derived cells were isolated from E12.5 *Wnt1^Cre^*;*Rosa26^tdTomato^*control and PLX5622 male and female embryos and cultured in 24-well plates for 10 days to allow for sphere formation^80^. Primary spheres were then dissociated, and cells were cultured for an additional 10 days to form secondary spheres. Primary and secondary neural crest-derived spheres retained *Wnt1^Cre^*-driven tdTomato+ signal and also expressed NCC markers such as PDGFRα^81^, SOX2^82,83^, and SOX10^84,85^, regardless of whether spheres were derived from E12.5 control or PLX5622 embryos (Figures 7B–7M). Intriguingly, gestational exposure to PLX5622 resulted in a significant decrease in the number of primary neural crest-derived spheres in both sexes (Figure 7N; significant effect of diet F_(1,25)_=51.00, p<0.0001; male p=0.0098, female p<0.0001), suggesting a reduced proliferative capacity of NCCs.

**Figure 7.**
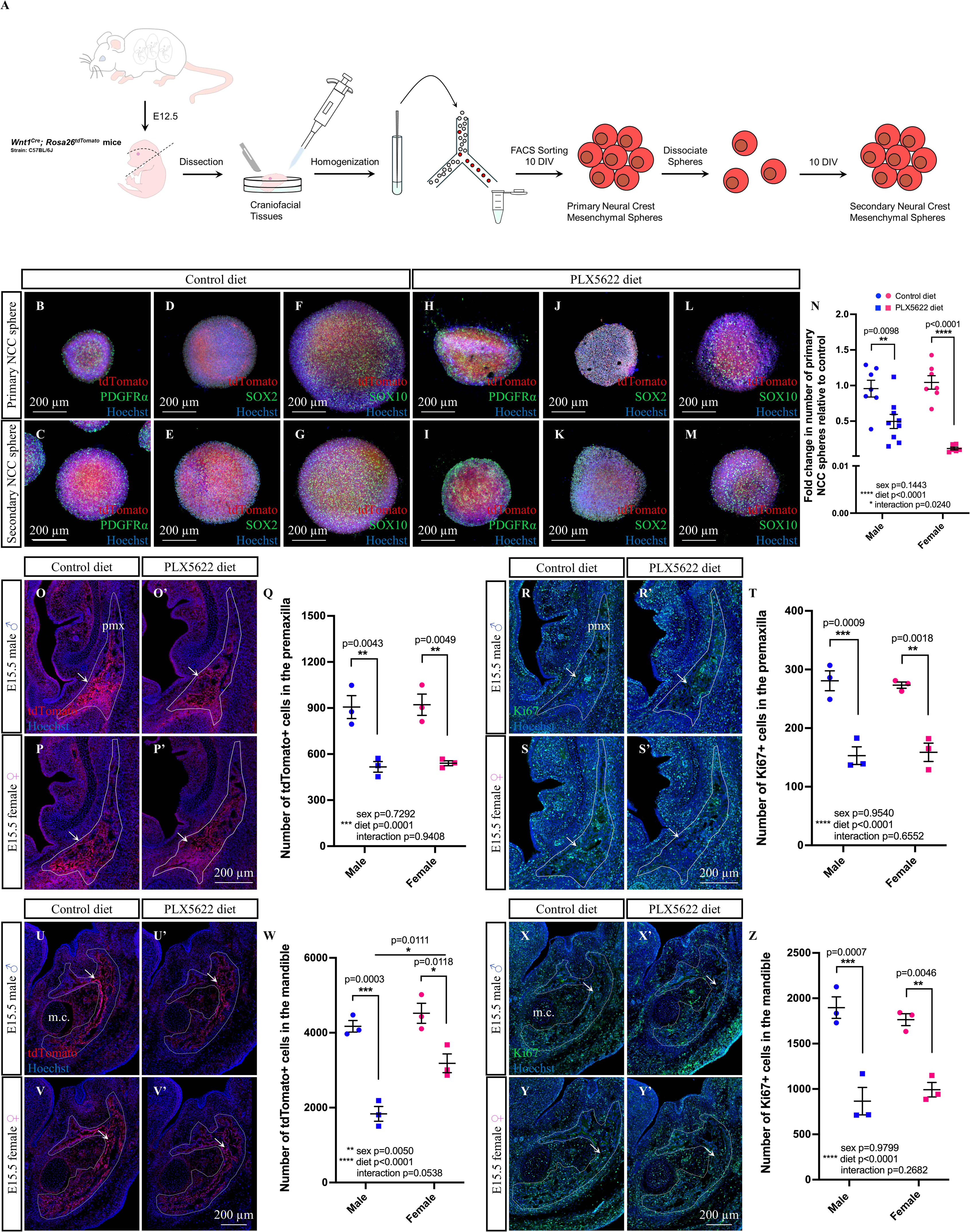
Gestational Exposure to PLX5622 Decreases Neural Crest Proliferation. (A) Schematic illustrating sphere assay. (B-M) E12.5 *Wnt1^Cre^; Rosa26^tdTomato^*neural crest-derived spheres express PDGFRα (B, C, H, I), SOX2 (D, E, J, K), and SOX10 (F, G, L, M). (N) Primary sphere quantification (N=6-9 embryos per sex/treatment from 2-3 dams). (O-Z) Immunofluorescent images of E15.5 *Wnt1^Cre^*-driven tdTomato+ signal (O-P’, U-V’) and Ki67 staining (R-S’, X-Y’) in the premaxilla (pmx; O-T) and mandible (U-Z). Quantification of tdTomato+ (Q, W) and Ki67+ (T, Z) cells. White lines mark the tissue regions used for quantifications (N=3 embryos per sex/treatment from 3 dams). Abbreviations: DIV, days *in vitro*; m.c., Meckel’s cartilage. Counts represent mean ± SEM and were analyzed by a two-way ANOVA with Tukey’s post hoc test.

To validate that embryonic NCC proliferation was indeed decreased in response to gestational PLX5622 exposure, we performed immunofluorescence staining on E15.5 *Wnt1^Cre^*;*Rosa26^tdTomato^* craniofacial tissue cryosections to quantify tdTomato+ cell numbers and proliferation using a Ki67 antibody. *Wnt1^Cre^*-driven tdTomato+ signal was found to be highly expressed within mineralizing regions of craniofacial bones—particularly, in regions where we had previously identified abnormal embryonic osteoblast staining/mineralization in response to PLX5622 exposure. Accordingly, we focused our analyses on the E15.5 premaxilla and mandible. Quantification of *Wnt1^Cre^*-driven tdTomato+ and Ki67+ cells (Figures 7O–T, arrows) in the E15.5 male and female premaxilla showed a significant decrease in both tdTomato+ neural crest-derived cells (Figure 7Q; main effect of diet F_(1,8)_=50.24, p=0.0001; male p=0.0043, female p=0.0049) and Ki67+ proliferating cells (Figure 7T; main effect of diet F_(1,8)_=74.75, p<0.0001; male p=0.0009, female p=0.0018) in PLX5622 embryos as compared to control. Similarly, quantification of *Wnt1^Cre^*-driven tdTomato+ and Ki67+ cells (Figures 7U–Z, arrows) in the E15.5 male and female mandible showed a significant decrease in tdTomato+ cells (Figure 7W; significant effect of diet F_(1,8)_=68.58, p<0.0001; male p=0.0003, female p=0.0118) and Ki67+ cells (Figure 7Z; main effect of diet F_(1,8)_=68.25, p<0.0001; male p=0.0007, female p=0.0046) in PLX5622 embryos as compared to control. Interestingly, E15.5 male embryos showed a more severe decrease in tdTomato+ cells in the mandible in response to PLX5622 exposure (Figure 7W; significant effect of sex F_(1,8)_=14.64, p=0.0050; PLX5622 male vs. female p=0.0111). Taken together, these data suggest that the proliferative capacity of cranial NCCs decreases in response to PLX5622 exposure.

## DISCUSSION

This study is the first to provide an in-depth characterization of CSF1R+ macrophage and osteoclast depletion in craniofacial tissues across embryogenesis in response to PLX5622 exposure^25–27,86–90^—a pharmacological tool that allows for temporal control and transient depletion of CSF1R-expressing cells. Exposure to PLX5622 significantly depleted CSF1R+ cells across embryogenesis, and either halts or significantly delays osteoclastogenesis, leading to a loss of bone resorptive activity throughout the developing craniofacial bones in the embryo. To our surprise, depleting CSF1R+ cells using PLX5622 does not impact the gross morphology of craniofacial nerves or muscles in male or female CD1 or C57BL/6 embryos. In contrast, craniofacial defects in the premaxilla and mandible appear as early as E15.5—the earliest stage at which a craniofacial phenotype has been reported in a CSF1R-disrupted model, while abnormalities in skull shape, cranial sutures, ear ossicles, palate, and cranial base development appear to arise later in embryogenesis, between E15.5 and birth. Although our data suggest that craniosynostosis does not account for the skull doming in P1 PLX5622 pups, disruptions to the cranial base and/or structural changes to the gross morphology of the brain could impact skull development and contribute to skull doming. Notably, we observed sex differences in P1 PLX5622 pups, whereby males presented with more prominent ear phenotypes and females displayed greater disruptions to the mandible. We also observed strain-dependent effects, with overall stronger phenotypes being observed in CD1 pups than in C57BL/6 pups. Interestingly, embryonic exposure to PLX5622 also significantly altered cytokine and chemokine levels in craniofacial tissues, with these changes likely being primarily driven by CSF1R+ cells—namely macrophages. These changes in signaling within craniofacial tissues have the potential to alter CSF1R+ cell dynamics and interactions with neighbouring cells, which could explain the observed decrease in proliferative capacity of NCCs in response to PLX5622 exposure. Altogether, a reduction in NCC proliferation in craniofacial tissues could contribute to alterations in differentiation and osteogenic potential, resulting in the phenotypic disruptions in the premaxilla and mandible, and likely the many other bony abnormalities seen in PLX5622 offspring.

Here, we used both a *Csf1r^EGFP^* transgenic fluorescent reporter and immunostaining against CSF1R to characterize CSF1R+ cell depletion in response to PLX5622 exposure. Although we found EGFP signal in cells negative for CSF1R immunostaining, this could be due to variable timing between *Csf1r^EGFP^* transgene expression and CSF1R protein or alternatively, EGFP signal could be retained^91–93^. In response to PLX5622 exposure from E3.5 onwards, CSF1R+ cell depletion was observed in all tissues examined and remained relatively stable (∼50%) across embryogenesis, albeit with notable sex differences in the timing of depletion. Apoptotic cells accumulated predominantly in nervous and muscular structures and not in or around developing cartilaginous/bony tissues, except for the ear. Indeed, programmed cell death of mesenchyme between the lens and retina at E10 in C57BL/6J mice is essential for normal eye development^94^; while, caspase-3 deficient C57BL/6 mice present with inner ear disruptions in the cochlea and vestibule and severe auditory dysfunction, demonstrating that caspase-3-dependent apoptosis is critical for normal inner ear development and function^95^. It is interesting to note that eye defects were previously reported in some PLX5622-exposed mice, where one eye could not be opened until later in adulthood^25^. Therefore, increased apoptosis resulting from PLX5622 exposure could perturb normal apoptotic programs and disrupt craniofacial morphogenesis, resulting in nerve and/or muscle phenotypes that could appear after E15.5—especially since bone disruptions exist across nearly the entire PLX5622 head and changes in the shape of mineralized tissue can directly impact craniofacial nerve and muscle development^96–102^. For example, the shape of the temporalis muscle can change upon deformation of the mandibular condyle, and the size of neurovascular bundles can be reduced upon removal of teeth from the mandible^96,97^. Accordingly, craniofacial nerve and muscle development in PLX5622-exposed offspring should be studied postnatally to confirm whether phenotypes eventually develop.

Interestingly, despite CSF1R+ cell deficiency, *Csf1r* KO rat phenotypes appear to develop postnatally, with normal placental and embryonic development reported^103^. Indeed, *Csf1/Csf1r*-disrupted genetic mouse models show craniofacial phenotypes consistently at P10, but as early as P6^9,10,15,104–106^. Here, our pharmacological CSF1R+ cell depletion model displays embryonic bone disruptions as early as E15.5 and multiple craniofacial bone abnormalities at P1—the vast majority of which are consistent with *Csf1/Csf1r*-disrupted genetic models (Table S3). As *Csf1/Csf1r*-disrupted genetic models and the PLX5622 pharmacological depletion model similarly display domed skulls^9–11,15,25,26,104–106^ (Table S3), and skull doming can be caused by midface hypoplasia and inhibited cranial base expansion^43–48^, we thoroughly assessed the cranial base. Although cranial base length was unaffected by PLX5622 exposure, suggesting that craniosynostosis and/or cranial base hypoplasia is likely not responsible for the skull doming seen in PLX5622 pups, the smaller cranial base bones and widened synchondroses observed in PLX5622 mice could contribute to the skull phenotype. We also noted enlarged lateral and third ventricles in PLX5622 pups, suggesting that mild hydrocephalus could contribute to skull doming, which has been seen in other animal models^49–54^. Moreover, we observed brain lesions at the cortico-striato-amygdalar boundary in PLX5622 pups—an area susceptible to physiological morphogenetic stress that is essential for fetal brain morphogenesis^107^. Given that microglia play critical roles in maintaining structural integrity at this boundary^107^, the loss of microglia upon PLX5622 exposure^25^ could disrupt normal maintenance and repair mechanisms, resulting in mechanical pressures on the cranial vault and skull doming; however, skull doming has not been reported in *Csf1r*^Δ*FIRE/*Δ*FIRE*^ mice that show similar brain lesions and loss of structural integrity^107,108^ (Table S3). Therefore, it is also possible that insufficient mineralization compromises the integrity of the calvaria such that it bulges outward in response to normal mechanical forces of the developing brain. Interestingly, the morphological disruptions observed in the brain and cranial base of PLX5622 mice are reminiscent of the brain abnormalities and cranial base sclerosis observed in <u>b</u>rain <u>a</u>bnormalities, <u>n</u>euro<u>d</u>egeneration, and <u>d</u>ys<u>os</u>teosclerosis (BANDDOS) patients—a human disorder caused by bi-allelic mutation in *CSF1R*^23,24,109–115^. The skull doming, cranial suture impairments, and cranial base disruptions seen in PLX5622 mice could be a useful model for studying the cranial vault malformations observed in BANDDOS^23,24,109,111^; especially considering the titratable nature of PLX5622, as CSF1R inhibition could more accurately reflect hypomorphic *CSF1R* mutations which account for 8/11 identified mutations causing BANDDOS^116^.

Although the phenotypes seen in *Csf1r/CSF1R*-disrupted rodents and humans are of an osteosclerotic nature^9,15,21,23,24,104,109,111,117,118^, likely due to a lack of osteoclasts, we observed decreased bone formation in the calvaria of PLX5622 pups and disruptions in the development of the premaxilla, mandible, ear bones, palate, and cranial base (Table S3). These phenotypes are unlikely to result solely from a loss of bone resorption by osteoclasts. Instead, prenatal exposure to PLX5622 could disrupt signaling from CSF1R+ macrophages that are required for bone formation and growth^21,119–124^. Intriguingly, these osteogenic pathways may also respond to PLX5622 differently between sexes given the sex-specific mandible and ear phenotypes observed and sex-dependent changes in several cytokines/chemokines. CSF1R+ osteoclasts are also more abundant and active in long bones of female mice compared to males, resulting in decreased bone mass^125–127^, and ovariectomy-induced estrogen deficiency increases CSF1 production^128–130^, which together further suggest a potential for sex hormone-dependent responses to CSF1R+ cell depletion following PLX5622 exposure. It is also interesting to note that we observed basal sex differences that were independent of PLX5622 exposure, which are in line with sex differences observed in craniofacial bones that can be observed at birth in humans^131^, and highlight the importance of studying both sexes in craniofacial research.

Interestingly, the impact of PLX5622 exposure on osteogenesis appears to be independent of whether the bone is formed by intramembranous (calvaria) or endochondral ossification (auditory ossicles)^132–140^. This suggests that signaling alterations may act on a common progenitor, which is supported by our findings as PLX5622 exposure alters cytokine/chemokine levels and NCCs give rise to the majority of the craniofacial bones found to be disrupted in PLX5622 mice^141–146^. Indeed, NCCs do respond to several of the cytokines/chemokines disrupted by PLX5622, including CCL2 which induces NCC migration into the mouse heart^147^, and CXCL1 and IL-6 which mediate NCC differentiation into osteoblasts during intramembranous ossification in a mouse calvarial defect model^148^. CCL2 and CXCL1 are also chemoattractants for osteoclast precursors and osteoclastogenesis^149–154^; while altering the secretion of any number of the other cytokines/chemokines found to change in our study could impact osteoclast function and bone remodeling^75,155–172^. Uniquely, IL-6 has been shown to be a critical regulator of bone remodeling by negatively regulating osteoblasts and modulating osteoclast numbers and resorption^173–177^. The downregulation of these signals in PLX5622-exposed mice could impact recruitment and differentiation of osteoblast and osteoclast precursors and disrupt bone remodeling processes necessary for normal craniofacial bone morphogenesis. Specifically, the decreases in bone size could be caused by reduced NCC proliferation and subsequent NCC deficiency, while the increased bone density observed from E15.5 onwards may be driven by decreased bone resorption and remodeling^178–182^. Alternatively, increased osteogenesis could reflect premature NCC differentiation into osteoblasts—possibly in response to altered CXCL1 and IL-6 signaling^148^, resulting in NCC progenitor deficiencies^183–185^. Although it is unclear how the disruptions to NCCs and osteoclast precursors/osteoclastogenesis culminate in the phenotypic disruptions seen in PLX5622 offspring, our findings suggest that CSF1R+ macrophages are a likely source for the many cytokines/chemokines found to be altered in response to PLX5622 exposure and may mediate NCC and/or osteoclast deficiencies.

Herein, we demonstrated that CSF1R+ cells are required for development of the skull, cranial sutures, premaxilla, mandible, ear ossicles, palate, and cranial base. Additionally, we showed that embryonic exposure to PLX5622 drives sex, strain, and time-dependent disruptions in craniofacial development. Together, these data highlight an underappreciated role for CSF1R+ macrophages and osteoclasts in craniofacial morphogenesis during the embryonic period and suggest that altered interactions with and/or signaling to NCCs may underlie the phenotypes observed in PLX5622 offspring.

### Limitations of the study

A potential limitation of our study is our use of the CSF1R inhibitor PLX5622 to deplete CSF1R-expressing cells. It has been reported that PLX5622 could have off-target effects on non-CSF1R-expressing brain endothelial cells, potentially due to inhibition of CSF1R-related tyrosine kinases FLT3, KIT, AURKC, and/or KDR^186^. KIT and KDR may be expressed by embryonic NCCs^187–193^, suggesting that PLX5622 could have off-target impacts on NCCs; therefore, future studies should examine changes to fetal NCCs in *Csf1r* KO mice. However, it is important to note that PLX5622 is more than 50-fold more active against CSF1R than KIT and KDR in cell-free enzyme assays^90^. The effects on endothelial cells are also specific to the central nervous system^186^. Indeed, brain alterations could impact development of the skull or surrounding craniofacial tissue; although, the postnatal brain phenotypes described herein match *Csf1r*^Δ*FIRE/*Δ*FIRE*^ and other microglia-depleted mice^107^, and the skull doming is consistent with phenotypes in *Csf1*/*Csf1r*-disrupted genetic models, suggesting that these phenotypes are not driven by disruption to brain endothelial cells^9–11,15,104–106^.

## RESOURCE AVAILABILITY

### Lead contact

Requests for further information and resources and reagents should be directed to and will be fulfilled by the lead contact, Dr. Jessica M. Rosin.

### Materials availability

This study did not generate new unique reagents.

### Data and code availability

- Images of control and PLX5622-exposed whole-mount E11.5, E12.5, and E13.5 nerve (2H3) and muscle (MF20) staining and P1 skeletal staining described in this study are available here: Felix Ma, Jessica M. Rosin. Phenotypic impacts of embryonic exposure to the CSF1R inhibitor PLX5622 on craniofacial development. FaceBase Consortium https://doi.org/10.25550/A9-12PJ
- This paper analyzes existing, publicly available data, accessible at the NCBI GEO database: GSM5324643, GSM6146223, and GSM7508877.
- All data reported in this paper will be shared by the lead contact upon request.
- This paper does not report original code.
- Seurat codes used for analysis of publicly available datasets deposited on GEO are available upon request.
- Any additional information required to reanalyze the data reported in this paper is available from the lead contact upon request.

## ACKNOWLEDGEMENTS

The authors thank Andrew Johnson and Justin Wong from the UBC Flow Core Facility for help with FACS, the UBC Centre for High-Throughput Phenogenomics—a facility supported by the Canada Foundation for Innovation, British Columbia Knowledge Development Foundation, and the UBC Faculty of Dentistry—for help with µCT scans, and the UBC Centre for Disease Modeling for animal care. We would also like to acknowledge Shruti S. Tophkhane, Joaquin I. Henriquez, Katherine Fu, Joy M. Richman and the entire Richman Lab for help with micromass culture and TRAP staining, and Siddharth R. Vora for help with the µCT scans. F.M. was supported by an NSERC PGS-D (579343-2023), a UBC Four Year Doctoral Fellowship, and a UBC Faculty of Dentistry Joseph Tonzetich Fellowship. R.R.J.Z. was supported by a UBC Faculty of Dentistry Summer Research Award (2022 & 2023) and a Charles Shuler Research Fellow award (2023). I.Z. and S.O. were supported by a UBC Faculty of Dentistry Summer Research Award (2025). V.B.W. was supported by an NSERC USRA. This work was supported by an NSERC Discovery Grant to J.M.R. (RGPIN-2022-03718). J.M.R. is a Michael Smith Health Research (MSHR) BC Scholar and a Tier 2 Canada Research Chair (CRC) in Immune Regulation of Developmental Programs.

## AUTHOR CONTRIBUTIONS

Conceptualization, F.M. and J.M.R.; data curation, F.M., M.R., and R.O.; formal analysis, F.M., R.O., and J.M.R.; funding acquisition, J.M.R.; investigation, F.M., R.R.J.Z., M.R., I.Z., S.O., R.O., V.B.W., and J.M.R.; methodology, F.M., M.R., R.O., and J.M.R.; project administration, J.M.R.; resources, F.M., M.R., and J.M.R.; software, F.M. and R.O.; supervision, J.M.R.; validation, F.M., M.R., R.O., and J.M.R.; visualization, F.M., M.R., R.O., and J.M.R.; writing—original draft, F.M. and J.M.R.; writing—review & editing, F.M., M.R., and J.M.R.

## SUPPLEMENTAL INFORMATION

Document S1. Figures S1–S7, Tables S1–S4, and supplemental references

## MATERIALS AND METHOD

### EXPERIMENTAL MODEL DETAILS

Animal work was carried out in accordance with guidelines and regulations of the Canadian Council of Animal Care and received prior approval from the University of British Columbia’s Animal Care Committee (protocols A21-0170 and A21-0171). CD1 (CR: 022; RRID: IMSR_CRL:022, Charles River) and C57BL/6 (JAX: 005304; RRID: IMSR_JAX:005304, The Jackson Laboratory) embryos and postnatal day 1 (P1) pups were utilized across the study, as described. *Csf1r^EGFP^*mice (JAX: 005304; RRID: IMSR_JAX:005304, The Jackson Laboratory) were crossed to C57BL/6 mice to generate heterozygous embryos for flow cytometry and immunofluorescence experiments. *Wnt1^Cre^* mice (JAX: 022501; RRID: IMSR_JAX:022501, The Jackson Laboratory) were crossed to *Rosa26^tdTomato^* mice (JAX: 007914; RRID: IMSR_JAX:007914, The Jackson Laboratory) to generate embryos for neural crest-derived sphere culture and for immunofluorescence experiments. Timed pregnancies were used to collect embryonic samples. For embryonic staging, female mice were plug-checked in the morning and those with a vaginal plug were assigned embryonic day 0.5 (E0.5). Female mice continued to receive standard chow (LabDiet PicoLab Rodent Diet 20) and water ad libitum during pregnancy, unless otherwise specified. For embryo collection, pregnant female mice were anaesthetized with isoflurane and immediately euthanized by cervical dislocation followed by decapitation. For postnatal staging (i.e., skeletal preparations, µCT scanning), the day of birth was assigned P0. All mouse embryos and pups were genotyped for sex and results were reported for both sexes.

### METHOD DETAILS

#### Mouse handling

Depletion of CSF1R+ cells was achieved by administering the Plexxikon CSF1R inhibitor PLX5622 (1200 PPM added to chow AIN-76A, Research Diets) to pregnant female mice starting at E3.5. Control dams received control diet (AIN-76A, Research Diets). Pregnant female mice were exposed to either control diet or PLX5622 diet from E3.5 until sample collection.

Following birth, female mice were placed on our standard chow (LabDiet PicoLab Rodent Diet 20) from P0 to P1.

#### Genotyping

Tails were collected from all mouse embryos/pups and incubated in extraction buffer (100 mM NaCl, 50 mM Tris, 100 mM EDTA, 1% SDS) with proteinase K (8 units, ∼0.4 mg/mL; New England Biolabs Cat#P8107S; UniProtKB: P06873) overnight at 56°C. DNA was extracted from digested tails by precipitation with saturated NaCl solution (∼6M), followed by centrifugation at 20,000 g for 10 min. DNA was precipitated from the supernatant with isopropanol, and pellets were collected by centrifugation at 20,000 g for 5 min. Pellets were washed with 70% ethanol and dried at 37°C. Dried DNA was dissolved in 200 µL of Tris-EDTA buffer (10 mM Tris, 1 mM EDTA, pH 8) for 1 h at 68°C, and 1.6 µL of DNA was subsequently added to OneTaq® Quick-Load® 2X Master Mix with Standard Buffer (New England Biolabs Cat#M0486S) along with 1 µM each of primers for SX (*Sly* & *Xlr*)^194^, *GFP*^195^, *Wnt1^Cre^*^196^, or *Rosa26^tdTomato^*^197^ (see Table S4 for primer sequences). PCR reactions were run on a Applied Biosystems™ MiniAmp™ Thermal Cycler (Applied Biosystems) as follows: 95°C for 3 min, 35 cycles of denaturation at 95°C for 30 sec, annealing at 60°C (57°C for SX primers) 30 sec, and extension at 72°C for 30 sec, followed by final extension at 72°C for 5 min. Amplified fragments were run on a 1.5% agarose gel in TAE buffer (0.4 M Tris, 0.2 M acetic acid, 10 mM EDTA, pH 8) and visualized using SmartGlow Pre-stain for nucleic acid gels (Accuris Instruments Cat#E4500-PS) on a Fisherbrand™ Real Time Electrophoresis System (Fisher Scientific). DNA fragments were sized according to GeneRuler Ready-to-Use 100bp DNA Ladder (Thermo Fisher Scientific Cat#SM0243).

#### Flow cytometry

Flow cytometry methods were adapted from Rosin et al^25^. E11.5, 13.5, 15.5, and 17.5 mouse embryo heads were collected from control and PLX5622-exposed *Csf1r^EGFP^* mice in PBS on ice. Craniofacial tissues were then micro-dissected away from the whole head while in PBS on ice. The micro-dissected craniofacial tissues were cut/dissociated into smaller pieces using a blade while in culture media on ice containing (v/v): 65.3% DMEM (GIBCO 11965-092, Thermo Fisher Scientific), 32.7% F-12 (GIBCO 11765-054, Thermo Fisher Scientific), and 2% B-27 supplement (GIBCO 17504-044, Thermo Fisher Scientific). The resulting dissociated tissue was filtered through a 35 μm strainer (Falcon 352235), placed in a chilled 1.5 mL tube and centrifuged at 300 g for 5 min at room temperature (RT). The resulting cell pellets were re-suspended in 500 μL cold HBSS with 5% FBS and filtered through a 35 μm strainer before flow cytometry. The resulting cell suspensions were analyzed by the University of British Columbia’s Flow Core Facility using a Beckman Coulter Life Sciences CytoFLEX LX machine and analyzed using Beckman Coulter CytExpert software (Beckman Coulter Life Sciences^198^; RRID:SCR_017217).

#### Tissue section immunohistochemistry

E15.5 *Csf1r^EGFP^*, *Wnt1^Cre^*;*Rosa26^tdTomato^*, and CD1 mouse embryo heads were collected in ice-cold PBS and fixed in 4% paraformaldehyde (PFA) overnight at 4°C. The tissues were then washed in PBS and equilibrated in 20% sucrose/PBS overnight at 4°C. Heads were embedded in Clear Frozen Section Compound (VWR, 95057-838) and cryosectioned (14 μm sections) on a Leica CM1950 cryostat (Nussloch, Germany). Cryosections were rehydrated in PBS, washed 4 × 10 min with PBT (PBS with 0.1% Triton X-100), permeabilized for 30 min with 1% Triton X-100 in PBS, blocked using 5% normal donkey serum (NDS, Sigma) in PBT for 1 h at RT, and exposed to sheep anti-CSF1R (1:200; R&D Systems Cat#AF3818; RRID: AB_884158), rabbit anti-Cathepsin K (1:200; Abcam Cat#ab19027; RRID: AB_2261274), rabbit anti-Active Caspase 3 (1:500; BD Pharmingen Cat#559565; RRID: AB_397274), rabbit anti-Sp7 (1:500; Abcam Cat#ab209484; RRID: AB_2892207), mouse 2H3 (1:200; Developmental Studies Hybridoma Bank Cat#2H3; RRID: AB_531793), mouse MF20 (1:100; Developmental Studies Hybridoma Bank Cat#MF 20; RRID: AB_2147781), goat anti-CCL3 (1:200; R&D Systems Cat#AF-450-NA; RRID: AB_354492), goat anti-CCL4 (1:200; R&D Systems Cat#AF-451-NA; RRID: AB_2071055), and/or mouse anti-Ki67 (1:200; BD Pharmingen Cat#556003; RRID: AB_396287) at 4°C overnight. Slides were then washed 4 × 10 min with PBT and exposed to secondary antibody (1:200, Alexa 488 or Alexa 594 donkey anti-sheep IgG, donkey anti-rabbit IgG, donkey anti-mouse IgG, or donkey anti-goat IgG, Invitrogen) for 2 h at RT. Nuclei were stained with 1:1000 Hoechst 33342 (Invitrogen Cat#H3570; CAS: 23491-52-3) in PBS for 5 min at RT and washed 3 × 5 min with PBT. Sections were mounted using Aqua Poly/Mount (Polysciences Inc.). Fluorescent images were captured on a ZEISS Axioplan fluorescent microscope with ZEISS Axiocam HRm camera or a ZEISS LSM900 confocal microscope and further processed using LSM+. Brightness and/or contrast of the entire image was adjusted using ZEISS ZEN 3.9 (RRID:SCR_013672; ZEISS Microscopy^199^) and/or Adobe Photoshop CC (RRID:SCR_014199; Adobe Inc.^200^) if deemed appropriate.

#### Tartrate-resistant acid phosphatase staining

E15.5 *Csf1r^EGFP^* mouse embryo head cryosections (described above) were rehydrated in distilled water and incubated for 30 min at 37°C in pre-warmed TRAP staining solution consisting of 0.1 mg/mL naphthol AS-MX phosphate (Sigma-Aldrich Cat#N4875; CAS: 1596-56-1), 0.6 mg/mL Fast Red Violet LB Salt (Sigma-Aldrich Cat#F3381; CAS: 32348-81-5), 11.4 mg/mL L-(+) Tartaric Acid (Sigma-Aldrich Cat#T109; CAS: 87-69-4), and 9.2 mg/mL sodium acetate anhydrous (Sigma) in 0.28% glacial acetic acid, with pH adjusted to 4.7–5.0 with 5M sodium hydroxide (Sigma). Slides were then washed with distilled water, and sections were mounted using Aqua Poly/Mount (Polysciences Inc.). Light microscope images were captured on a ZEISS Axiocam 208 color camera mounted on a ZEISS Stemi 508 microscope. Brightness and/or contrast of the entire image was adjusted using Adobe Photoshop CC if deemed appropriate.

#### Whole-mount immunohistochemistry

Whole-mount staining methods were adapted from Rosin et al^201^. Antibodies recognizing neurofilament (2H3) and muscle myosin (MF20) were used to visualize embryonic nerves and muscles. E11.5, 12.5, and 13.5 whole CD1 and C57BL/6 mouse embryos were fixed overnight in Dent’s fixative (4:1, Methanol:DMSO) at 4°C. Embryos were bleached overnight in 5:1 Dent’s fixative:30% H_2_O_2_ at RT, then rehydrated with successive 30-minute washes in 50%/15%/0% Methanol in PBS–0.5% Tween 20 (PBST). Embryos were blocked two times for 1 h in PBST/1% DMSO/2% skim milk powder (PBSTMD) at RT then incubated overnight at 4°C with primary antibody diluted 1:130–1:150 in PBSTMD to achieve a final concentration of 2 μg/mL. Embryos were then washed four times for 1 h with PBST, blocked with PBSTMD, and incubated overnight at 4°C with peroxidase-conjugated goat anti-mouse IgG (Sigma A-9169) diluted 1:300 in PBSTMD, then washed four times for 1 h with PBST, stained with 0.5 mg/mL DAB (3,3′-Diaminobenzidine tetrahydrochloride; Sigma-Aldrich Cat#D5905; CAS: 7411-49-6) and 0.3% H_2_O_2_ in PBST, dehydrated to 100% methanol, and cleared in BABB (1:2, benzyl alcohol:benzyl benzoate). Light microscope images were captured on a ZEISS Axiocam 208 color camera mounted on a ZEISS Stemi 508 microscope. Brightness and/or contrast of the entire image was adjusted using Adobe Photoshop CC if deemed appropriate.

#### Von Kossa staining

Von Kossa staining was adapted from Tosun et al^202^. E15.5 CD1 mouse embryo head cryosections (described above) were rehydrated in distilled water and incubated in 0.45 µm-filtered 2% silver-nitrate solution (Sigma-Aldrich Cat#209139; CAS: 7761-88-8) under direct light exposure (14 W, 1500 lumen bulb) for 1 hour, washed with 1% acetic acid, and subsequently stained for 20 minutes in 0.02% Alcian blue in 70% ethanol and 30% acetic acid. Sections were counterstained in 0.1% Nuclear Fast Red Solution for 10 minutes and rinsed in distilled water before mounting with Aqua Poly/Mount (Polysciences Inc.). Slides were imaged on a ZEISS Axiocam 208 color camera mounted on a ZEISS Stemi 508 microscope.

#### Micromass culture

Micromass culture methods were adapted from Tophkhane et al, Ralphs, and Ueharu et al^203–205^. E12.5 CD1 mouse embryo heads were collected and the frontonasal mass and mandible were dissected in cold Hank’s balanced saline solution (HBSS, without calcium and magnesium; Thermo Fisher Scientific, #14185052) with 10% FBS and 1% Penicillin/Streptomycin. Dissected craniofacial tissues were individually incubated in 2% trypsin (Gibco) on ice for 1 h. HBSS with 10% FBS and 1% Penicillin/Streptomycin was added to inhibit the enzymatic activity of trypsin, pipetting up and down to dissociate the ectoderm and mesenchyme. The cell solution was then centrifuged at 1000 g for 5 min. The supernatant was removed, and the cells were resuspended in HBSS with 10% FBS and 1% Penicillin/Streptomycin. The resulting suspension was filtered through a 35 μm strainer. The mesenchymal cells were counted using a Countess II automated cell counter (Thermo Fisher Scientific) and were resuspended to 2×10^7^ cells/mL in chondrogenic medium containing 40% DMEM (GIBCO 11995-065, Thermo Fisher Scientific) and 60% F12 (GIBCO 11765-054, Thermo Fisher Scientific) supplemented with 10% FBS, GlutaMAX (Gibco), 50 µg/mL ascorbic acid (Thermo Fisher Scientific Cat#850-3080IM; CAS: 50-81-7), 10 mM β-glycerol phosphate (Sigma-Aldrich Cat#G9422; CAS: 154804-51-0) and 1% Penicillin/Streptomycin. Cells were dropped into Nunc cell culture-treated 24-well plates (Thermo Fisher Scientific, Cat#142475) in 10 µL drops at 2×10^5^ cells/well in the center of the well and incubated at 37°C and 5% CO_2_ for 90 min to allow cells to attach and then flooded with 1 mL/well of chondrogenic medium. The culture medium was changed every other day (days 3, 5, and 7) until day 8. On day 8, cultures were fixed in 4% PFA for 30 min at RT and stored at 4°C in 100 mM Tris (pH 8.3) until staining. Fixed cultures were incubated at RT in 0.6 mg/mL Fast red violet LB salt and 0.1 mg/mL naphthol AS-MX phosphate in 100 mM Tris (pH 8.3) for 60 min to stain for alkaline phosphatase activity. The cultures were then stained with 1% Alcian blue in 3% acetic acid and 1% HCl to detect the area occupied by cartilage. All cultures were counterstained with 50% Shandon’s Instant Hematoxylin (Thermo Fisher Scientific Cat#6765015) and stored in 100% glycerol. Stained cultures were visualized on a ZEISS Stemi 508 light microscope, and images were captured on a ZEISS Axiocam 208 color camera.

#### Bone and cartilage skeletal staining

P1 CD1 and C57BL/6 mouse pups were collected and sacrificed by decapitation. Eyes were removed from skinned heads, and heads were then fixed in ethanol with 1% glacial acetic acid for at least 24 h and placed in 0.45 μm-filtered Alcian blue solution (1 mg/mL Alcian blue 8GX in 80% ethanol and 20% glacial acetic acid; Sigma-Aldrich Cat#05500; CAS: 33864-99-2) overnight to stain cartilage. After washing with ethanol for two 1 h washes, mouse heads were transferred to 1.5% KOH solution for 3.5 h, then stained with Alizarin Red S solution (0.15 mg/mL Alizarin red S in 0.5% KOH; Sigma-AldrichCat#A5533; CAS: 130-22-3) for 3 h to stain bone. Skulls were cleared by incubation in 0.5% KOH with 20% glycerol two times for 2–3 days and stored in a 40% glycerol mixture. Light microscope images were captured on a ZEISS Axiocam 208 color camera mounted on a ZEISS Stemi 508 microscope. Brightness and/or contrast of the entire image was adjusted using Adobe Photoshop CC if deemed appropriate.

#### Micro-computed tomography (CT)

P1 CD1 mouse pups were collected and sacrificed by decapitation. Heads were frozen at –20°C in 15 mL Falcon tubes. For contrast enhanced micro-CT scanning, P1 heads were fixed via overnight incubation with 4% PFA at 4°C. The samples were then dehydrated through serial incubations with increasing methanol concentrations (30%, 50% and 70%) in PBS. Subsequently, samples were incubated in a 1% phosphotungstic acid (PTA)/90% methanol solution for 2 weeks, ending with rehydration through serial incubations with decreasing methanol concentrations (70%, 50% and 30%). Incubation with each methanol concentration at both the dehydration and rehydration steps was done for 2 days. Micro-CT scans were performed with the Scanco Medical μCT100 scanner at 55 kVp and 200 μA in the Centre for High-Throughput Phenogenomics at the University of British Columbia. PTA-stained specimens were scanned at 7 µm resolution while embedded in 1% agarose. Additionally, unstained heads were micro-CT scanned after thawing at RT for 1 hour, at 15 µm resolution while embedded in 1% agarose. All outputs were exported as DICOM files.

#### Craniofacial tissue culture for cytokine and chemokine analysis

Methods for tissue culture for cytokine/chemokine analysis were adapted from Rosin et al^206^. E13.5 C57BL/6 mouse embryo heads were collected in PBS at RT. Craniofacial tissues were then micro-dissected away from the whole head and placed in a cell culture dish with 400 μL of 37°C culture media containing (v/v): 65.3% DMEM, 32.7% F-12, and 2% B27 supplement. Craniofacial tissues were cultured in a humid incubator with ambient oxygen and 5% CO_2_ at 37 °C for 6 h. Culture media was collected and spun down at 3000 g for 10 min at RT, then supernatant was collected into a new tube, flash frozen, and stored at –75°C until sent for multiplexed quantification of 45 mouse cytokines, chemokines, and growth factors using Luminex xMAP technology with MILLIPLEX® Mouse Cytokine/Chemokine Magnetic Bead Panel (Millipore Cat#MCYTOMAG-70K) and MILLIPLEX® Mouse Cytokine/Chemokine Magnetic Bead Panel II (Millipore Cat#MECY2MAG-73K), according to the manufacturer’s protocol. The analytes were evaluated on the Luminex™ 200 system (Luminex, Austin, TX, USA) by Eve Technologies Corp. (Calgary, AB, Canada).

#### Single-cell RNA sequencing data analysis

Publicly available single-cell RNA sequencing datasets of wild-type E12.5 and E13.5 C57BL/6J and FVB/NJ mouse craniofacial mesenchyme were downloaded from the NCBI GEO database^58,59^. Accession numbers GSM5324643 (E13.5 C57BL/6J), GSM6146223 (E12.5 C57BL/6J), and GSM7508877 (E12.5 FVB/NJ) were imported into Seurat v5 (RRID:SCR_016341; Hao et al.^207^, Hao et al.^208^, Stuart et al.^209^, Butler et al.^210^, Satija et al.^211^) in RStudio (RRID:SCR_000432; Posit team^212^), which runs on the R Project for Statistical Computing software environment (RRID:SCR_001905; R Core Team^213^), for analysis. Low quality cells (>20,000 counts, >6,000 genes detected, or >20% mitochondrial genes for GSM6146223; >35,000 counts, >7,000 genes detected, or >15% mitochondrial genes for GSM5324643; <1,500 or >7,500 genes detected for GSM7508877) were filtered from downstream analysis as they may have arisen from damaged cells, multiplets, or other processing artifacts. Counts were normalized and datasets were filtered to leave only cells with non-zero *Csf1r* expression. *Csf1r*-expressing cell clusters were identified using graph-based clustering with the *FindClusters* function, based on variable features and similar cells identified by *FindNeighbors* function. The *clustree* package was used to assist in determining 0.2 as the appropriate resolution for clustering, and proper clustering was then verified by manual inspection of marker genes. Uniform Manifold Approximation and Projections (UMAP) were visualized using *DimPlot* and gene signatures were visualized using *Vlnplot*. A cluster of hemoglobin-enriched cells did not express any markers of monocytes, microglia, macrophages, or osteoclasts except *Csf1r*; this cluster was excluded in the final analysis.

#### Fluorescence *in situ* hybridization

E15.5 CD1 mouse embryo head cryosections (described above) were rehydrated in PBS and stained according to manufacturer’s protocol using the RNAscope™ Multiplex Fluorescent Reagent Kit v2 (Advanced Cell Diagnostics Cat#323100). Briefly, sections were post-fixed for 15 min at 4°C in 4% PFA. Slides were then incubated in hydrogen peroxide for 10 min, washed with water, and incubated in boiling target retrieval buffer for 5 min. Sections were washed with water and dehydrated in 100% ethanol for 3 min and treated with protease 3 for 30 min at 40°C in a ACD HybEZ™ II Hybridization System (Advanced Cell Diagnostics). Slides were washed with water and probes for *Csf1r* (Advanced Cell Diagnostics, Cat#428191-C3) and *Ccl4* (Advanced Cell Diagnostics, Cat#421071) were hybridized to tissue for 2h at 40°C, followed by incubation in amplification reagents and TSA Vivid 520 (Advanced Cell Diagnostics Cat#323271) and TSA Vivid 570 (Advanced Cell Diagnostics Cat#323272) fluorophores, with washes in wash buffer between incubations. Nuclei were stained with DAPI for 30 seconds and slides were mounted using Aqua Poly/Mount (Polysciences Inc.). High-resolution images were captured on a ZEISS LSM900 confocal microscope and were further processed using LSM+. Brightness and/or contrast of the entire image was adjusted using ZEISS ZEN 3.9 and/or Adobe Photoshop CC if deemed appropriate.

#### Neural crest cell sphere culture

NCC sphere culture methods were adapted from Rosin et al. and Hagiwara et al^206,214^. E12.5 mouse heads were collected (*Wnt1^Cre^* mice crossed to *Rosa26^tdTomato^*mice to label neural crest-derived cells with tdTomato) in 37°C PBS containing 1% Penicillin/Streptomycin. Craniofacial tissues were then micro-dissected away from the whole head while in PBS containing 1% Penicillin/Streptomycin. Micro-dissected craniofacial tissues were cut/dissociated into smaller pieces using a sterile blade. The resulting dissociated tissue was filtered through a 35 μm strainer, placed in a 1.5 mL tube, and centrifuged at 300 g for 10 min at RT. The resulting cell pellets were re-suspended in 500 μL of 37 °C warmed culture media containing (v/v): 98% DMEM/F-12 (no phenol red; GIBCO 21041-025, Thermo Fisher Scientific) and 2% B-27 supplement, with 20 ng/mL each of human recombinant epidermal growth factor (STEMCELL Technologies Cat#78006.1) and basic fibroblast growth factor (STEMCELL Technologies Cat#78003.1) added. The resulting cell suspensions were filtered through a 35 μm strainer before fluorescence-activated cell sorting (FACS) by the University of British Columbia’s Flow Core Facility using a Beckman Coulter MoFlo Astrios EQ cell sorter. tdTomato+ neural crest-derived cells were collected in 500 μL of culture media. Cells were plated at 12,500 cells/mL per well in 24-well plates and were incubated for 10 days at 37°C with 5% CO_2_, with 50% media replenishment at day 5. At day 10, primary spheres were counted and imaged, dissociated in sterile media, counted, and replated at 1,000 cells/mL and cultured again for 10 days as above for secondary sphere formation.

#### Neural crest sphere immunohistochemistry

Cultured neural crest-derived spheres (described above) were fixed in 4% PFA for 30 min at RT and stored in PBS at 4°C until staining. Spheres were washed with PBT (PBS with 0.1% Triton X-100), blocked using 5% NDS (Sigma) for 1 h at RT, and exposed to goat anti-PDGFRα (1:150; R&D Systems Cat#AF1062; RRID: AB_2236897), rabbit anti-SOX2 (1:500; Millipore Cat#AB5603; RRID: AB_2286686) or mouse anti-SOX10 (1:500; R&D Systems Cat#MAB2864; RRID: AB_2195180) at 4°C overnight. Spheres were then washed with PBT and exposed to secondary antibody (1:200, Alexa 488 donkey anti-goat IgG, donkey anti-rabbit IgG or donkey anti-mouse IgG, Invitrogen) for 2 h at RT. Nuclei were stained with Hoechst 33342 (Invitrogen) for 5 min at RT. Spheres were aspirated with a wide bore pipette tip and deposited onto a glass microscope slide, and were subsequently mounted using Aqua Poly/Mount (Polysciences Inc.). High-resolution images were captured on a ZEISS LSM900 with confocal microscope and were further processed using LSM+. Brightness and/or contrast of the entire image was adjusted using ZEISS ZEN 3.9 and/or Adobe Photoshop CC if deemed appropriate.

#### Phagocytosis assay

E13.5 mouse embryo heads were collected from *Csf1r^EGFP^*mice in 37°C PBS. *Csf1r^EGFP^* embryos were screened for EGFP expression on a ZEISS Axiovert 5 microscope. *Csf1r^EGFP+^* craniofacial tissues were then micro-dissected away from the whole head while in 37°C PBS. The micro-dissected craniofacial tissues were bisected along the midline using a blade, and each half was cut once more into rostral and caudal quarters. Each half-head (one rostral quarter + one caudal quarter of the same side) was placed in 300 µL of 37°C culture media containing (v/v): 60.7% DMEM (GIBCO 11965-092, Thermo Fisher Scientific), 30.3% F-12 (GIBCO 11765-054, Thermo Fisher Scientific), 2% B-27 supplement (GIBCO 17504-044, Thermo Fisher Scientific), 7% PBS, 10 ng/mL CCL2 (R&D Systems, Cat#479-JE), 20 ng/mL CCL3 (R&D Systems, Cat#450-MA), 40 ng/mL CCL4 (R&D Systems, Cat#451-MB), 20 ng/mL CCL12 (R&D Systems, Cat#428-P5), 30 ng/mL CXCL2 (R&D Systems, Cat#452-M2), 30 ng/mL CXCL10 (R&D Systems, Cat#466-CR), and 20 ng/mL IL-6 (R&D Systems, Cat#406-ML). Each contralateral half-head was placed in 300 µL of identical culture media without cytokines. All craniofacial tissues were incubated at 37°C with 5% CO_2_ for 3 hours. pHrodo™ Deep Red E. coli BioParticles™ Conjugate (Invitrogen P35360, Thermo Fisher Scientific) were warmed and homogenized by addition of 171.4 µL of 37°C culture media to 400 µL of BioParticles. 50 µL of 37°C BioParticles-media was added to each 300 µL craniofacial culture. Craniofacial cultures were further incubated at 37°C with 5% CO_2_ for 1 hour. After 1 hour incubation, craniofacial tissues were removed from culture media with fine forceps and placed in 3 cm Petri dishes containing 37°C HBSS with 5% FBS. The craniofacial tissues were cut/dissociated into smaller pieces using a blade, filtered through a 35 μm strainer (Falcon 352235), placed in a 1.5 mL tube, and centrifuged at 300 g for 5 min at RT. The resulting cell pellets were re-suspended in 500 μL 37°C HBSS with 5% FBS and filtered through a 35 μm strainer before flow cytometry. The resulting cell suspensions were analyzed by the University of British Columbia’s Flow Core Facility using a Beckman Coulter Life Sciences CytoFLEX LX machine and analyzed using Beckman Coulter CytExpert software.

## QUANTIFICATION AND STATISTICAL ANALYSIS

E15.5 craniofacial tissue sections were obtained by cryosectioning PFA-fixed frozen embryo heads on a Leica CM1950 cryostat (Nussloch, Germany). Tissue was sectioned coronally starting from the nose, and 14 µm serial tissue sections were captured across 10 slide sets, with 21 embryonic head sections placed on each slide, and each set comprised of 2 slides. A single set (2 slides) of serial sectioned slides was used for each staining experiment described above. Every 14 μm section containing the craniofacial structure of interest was used for cell counts of cells with positive expression of *Csf1r^EGFP^*, CSF1R, cathepsin K, cleaved caspase 3, *Wnt1^Cre^;Rosa26^tdTomato^*, and Ki67 (N=3 per sex/treatment from 2-3 dams). For CSF1R, *Csf1r^EGFP^*, cathepsin K, and cleaved caspase 3 counts, positive cells were counted within the ear, eye, trigeminal, tongue, and mandible. A 5-cell radius was used for counting cells surrounding the nasal septum, Meckel’s cartilage, and maxillary incisor. For *Wnt1^Cre^;Rosa26^tdTomato^* counts, the mineralizing region of the premaxilla and mandible were outlined and cells within this area were counted. For Ki67 counts, mineralizing areas of the premaxilla and mandible were outlined comparably to the *Wnt1^Cre^;Rosa26^tdTomato^* outlines, and Ki67-positive cells within this area were counted. For micromass alkaline phosphatase staining analysis, images were adjusted by Fiji v2.16 (RRID:SCR_002285; Schindelin et al.^215^) with ImageJ v1.54 (RRID:SCR_003070; Schneider et al.^216^) with the *Color Threshold* function using *red, green, and blue (RGB)* color space^215–217^. The mask created by the threshold color was set to *black and white (B&W)*. The image was converted to binary by setting *Image Type* to *8-bit*. The stained area was first delineated by the *Freehand selections* tool, and the *Edit Clear Outside* function was used to eliminate pixels outside of the selected area. *Analyze Particles* was used to measure the total stained area. The process was repeated for hematoxylin-stained images to obtain the total micromass culture area. The same process was also used to analyze the TRAP+ area in the premaxilla, maxilla, and mandible, and for analyzing the von Kossa-stained mineralized area in the maxilla for craniofacial tissue sections. For analysis of the premaxilla and mandible of von Kossa-stained sections, images were imported into Fiji v2.16 (RRID:SCR_002285; Schindelin et al.^215^) with ImageJ v1.54 (RRID:SCR_003070; Schneider et al.^216^) and the unstained area within each mineralized structure was measured with *Freehand selections* tool and *Measure*. For neural crest-derived sphere analysis, each well was quantified by visualization on a ZEISS Axiovert 25 at 10x magnification, and a minimum cell cluster diameter of 40 µm was used as a cut-off to qualify as a counted sphere. Quantitative results (N=3-6 mouse embryos from 2-4 dams, unless otherwise mentioned in the Results and/or Figure Legends) for all counts and measurements are represented by mean ± SEM and were either analyzed by a two-tailed unpaired t-test, two-way ANOVA with Tukey’s post hoc analysis in GraphPad Prism 9 (RRID:SCR_002798; GraphPad Software^218^) or an aligned ranks transformation ANOVA with Tukey’s post hoc analysis using ARTool v0.11.2 (RRID:SCR_027412; Kay et al.^219,220^) in RStudio (RRID:SCR_000432; Posit team^212^), which runs on the R Project for Statistical Computing software environment (RRID:SCR_001905; R Core Team^213^). Statistical significance was defined as p<0.05.

## REFERENCES

1. Arai, F., Miyamoto, T., Ohneda, O., Inada, T., Sudo, T., Brasel, K., Miyata, T., Anderson, D.M., and Suda, T. (1999). Commitment and Differentiation of Osteoclast Precursor Cells by the Sequential Expression of C-Fms and Receptor Activator of Nuclear Factor κb (Rank) Receptors. Journal of Experimental Medicine 190, 1741–1754. 10.1084/jem.190.12.1741.

2. Boulakirba, S., Pfeifer, A., Mhaidly, R., Obba, S., Goulard, M., Schmitt, T., Chaintreuil, P., Calleja, A., Furstoss, N., Orange, F., et al. (2018). IL-34 and CSF-1 display an equivalent macrophage differentiation ability but a different polarization potential. Sci Rep 8, 256. 10.1038/s41598-017-18433-4.

3. Marks, D.C., Csar, X.F., Wilson, N.J., Novak, U., Ward, A.C., Kanagasundarum, V., Hoffmann, B.W., and Hamilton, J.A. (1999). Expression of a Y559F Mutant CSF-1 Receptor in M1 Myeloid Cells: A Role for Src Kinases in CSF-1 Receptor-Mediated Differentiation. Molecular Cell Biology Research Communications 1, 144–152. 10.1006/mcbr.1999.0123.

4. Giulian, D., and Ingeman, J.E. (1988). Colony-stimulating factors as promoters of ameboid microglia. J Neurosci 8, 4707–4717. 10.1523/JNEUROSCI.08-12-04707.1988.

5. Tushinski, R.J., Oliver, I.T., Guilbert, L.J., Tynan, P.W., Warner, J.R., and Stanley, E.R. (1982). Survival of mononuclear phagocytes depends on a lineage-specific growth factor that the differentiated cells selectively destroy. Cell 28, 71–81. 10.1016/0092-8674(82)90376-2.

6. Feng, X., Takeshita, S., Namba, N., Wei, S., Teitelbaum, S.L., and Ross, F.P. (2002). Tyrosines 559 and 807 in the cytoplasmic tail of the macrophage colony-stimulating factor receptor play distinct roles in osteoclast differentiation and function. Endocrinology 143, 4868–4874. 10.1210/en.2002-220467.

7. Chihara, T., Suzu, S., Hassan, R., Chutiwitoonchai, N., Hiyoshi, M., Motoyoshi, K., Kimura, F., and Okada, S. (2010). IL-34 and M-CSF share the receptor Fms but are not identical in biological activity and signal activation. Cell Death Differ 17, 1917–1927. 10.1038/cdd.2010.60.

8. Delaney, C., Farrell, M., Doherty, C.P., Brennan, K., O’Keeffe, E., Greene, C., Byrne, K., Kelly, E., Birmingham, N., Hickey, P., et al. (2021). Attenuated CSF-1R signalling drives cerebrovascular pathology. EMBO Mol Med 13, e12889. 10.15252/emmm.202012889.

9. Dai, X.M., Ryan, G.R., Hapel, A.J., Dominguez, M.G., Russell, R.G., Kapp, S., Sylvestre, V., and Stanley, E.R. (2002). Targeted disruption of the mouse colony-stimulating factor 1 receptor gene results in osteopetrosis, mononuclear phagocyte deficiency, increased primitive progenitor cell frequencies, and reproductive defects. Blood 99, 111–120. 10.1182/blood.v99.1.111.

10. Marks, S.C., Jr., and Lane, P.W. (1976). Osteopetrosis, a new recessive skeletal mutation on chromosome 12 of the mouse. J Hered 67, 11–18. 10.1093/oxfordjournals.jhered.a108657.

11. Pridans, C., Raper, A., Davis, G.M., Alves, J., Sauter, K.A., Lefevre, L., Regan, T., Meek, S., Sutherland, L., Thomson, A.J., et al. (2018). Pleiotropic Impacts of Macrophage and Microglial Deficiency on Development in Rats with Targeted Mutation of the Csf1r Locus. J Immunol 201, 2683–2699. 10.4049/jimmunol.1701783.

12. Van Wesenbeeck, L., Odgren, P.R., MacKay, C.A., D’Angelo, M., Safadi, F.F., Popoff, S.N., Van Hul, W., and Marks, S.C., Jr. (2002). The osteopetrotic mutation toothless (tl) is a loss-of-function frameshift mutation in the rat Csf1 gene: Evidence of a crucial role for CSF-1 in osteoclastogenesis and endochondral ossification. Proc Natl Acad Sci U S A 99, 14303–14308. 10.1073/pnas.202332999.

13. Pollard, J.W., Hunt, J.S., Wiktor-Jedrzejczak, W., and Stanley, E.R. (1991). A pregnancy defect in the osteopetrotic (opop) mouse demonstrates the requirement for CSF-1 in female fertility. Developmental Biology 148, 273–283. 10.1016/0012-1606(91)90336-2.

14. Erblich, B., Zhu, L., Etgen, A.M., Dobrenis, K., and Pollard, J.W. (2011). Absence of colony stimulation factor-1 receptor results in loss of microglia, disrupted brain development and olfactory deficits. PLoS One 6, e26317. 10.1371/journal.pone.0026317.

15. Okano, T., and Kishimoto, I. (2019). Csf1 Signaling Regulates Maintenance of Resident Macrophages and Bone Formation in the Mouse Cochlea. Frontiers in Neurology 10. 10.3389/fneur.2019.01244.

16. Sundquist, K., Cecchini, M., and Marks Jr, S. (1995). Colony-stimulating factor-1 injections improve but do not cure skeletal sclerosis in osteopetrotic (op) mice. Bone 16, 39–46.

17. Begg, S.K., Radley, J.M., Pollard, J.W., Chisholm, O.T., Stanley, E.R., and Bertoncello, I. (1993). Delayed hematopoietic development in osteopetrotic (op/op) mice. Journal of Experimental Medicine 177, 237–242. 10.1084/jem.177.1.237.

18. Cecchini, M.G., Dominguez, M.G., Mocci, S., Wetterwald, A., Felix, R., Fleisch, H., Chisholm, O., Hofstetter, W., Pollard, J.W., and Stanley, E.R. (1994). Role of colony stimulating factor-1 in the establishment and regulation of tissue macrophages during postnatal development of the mouse. Development 120, 1357–1372. 10.1242/dev.120.6.1357.

19. Wiktor-Jedrzejczak, W.W., Ahmed, A., Szczylik, C., and Skelly, R.R. (1982). Hematological characterization of congenital osteopetrosis in op/op mouse. Possible mechanism for abnormal macrophage differentiation. J Exp Med 156, 1516–1527. 10.1084/jem.156.5.1516.

20. Hume, D.A., Caruso, M., Ferrari-Cestari, M., Summers, K.M., Pridans, C., and Irvine, K.M. (2020). Phenotypic impacts of CSF1R deficiencies in humans and model organisms. Journal of Leukocyte Biology 107, 205–219. 10.1002/JLB.MR0519-143R.

21. Keshvari, S., Caruso, M., Teakle, N., Batoon, L., Sehgal, A., Patkar, O.L., Ferrari-Cestari, M., Snell, C.E., Chen, C., Stevenson, A., et al. (2021). CSF1R-dependent macrophages control postnatal somatic growth and organ maturation. PLoS Genet 17, e1009605. 10.1371/journal.pgen.1009605.

22. Patkar, O.L., Caruso, M., Teakle, N., Keshvari, S., Bush, S.J., Pridans, C., Belmer, A., Summers, K.M., Irvine, K.M., and Hume, D.A. (2021). Analysis of homozygous and heterozygous Csf1r knockout in the rat as a model for understanding microglial function in brain development and the impacts of human CSF1R mutations. Neurobiol Dis 151, 105268. 10.1016/j.nbd.2021.105268.

23. Guo, L., Bertola, D.R., Takanohashi, A., Saito, A., Segawa, Y., Yokota, T., Ishibashi, S., Nishida, Y., Yamamoto, G.L., Franco, J., et al. (2019). Bi-allelic CSF1R Mutations Cause Skeletal Dysplasia of Dysosteosclerosis-Pyle Disease Spectrum and Degenerative Encephalopathy with Brain Malformation. Am J Hum Genet 104, 925–935. 10.1016/j.ajhg.2019.03.004.

24. Monies, D., Maddirevula, S., Kurdi, W., Alanazy, M.H., Alkhalidi, H., Al-Owain, M., Sulaiman, R.A., Faqeih, E., Goljan, E., Ibrahim, N., et al. (2017). Autozygosity reveals recessive mutations and novel mechanisms in dominant genes: implications in variant interpretation. Genet Med 19, 1144–1150. 10.1038/gim.2017.22.

25. Rosin, J.M., Vora, S.R., and Kurrasch, D.M. (2018). Depletion of embryonic microglia using the CSF1R inhibitor PLX5622 has adverse sex-specific effects on mice, including accelerated weight gain, hyperactivity and anxiolytic-like behaviour. Brain Behav Immun 73, 682–697. 10.1016/j.bbi.2018.07.023.

26. Nagra, A., Katsube, M., Gao, W., Rosin, J.M., and Vora, S.R. (2023). Embryonic inhibition of colony-stimulating factor 1 receptor impacts craniofacial morphogenesis. Orthod Craniofac Res 26 *Suppl 1*, 20–28. 10.1111/ocr.12671.

27. Yongzhen, L., Yan, G., Jing, L., Chenyan, R., Chuanqing, M., Yun, S., and Weihui, C. (2024). Embryonic inhibition of colony-stimulating factor 1 receptor induces enlarged cartilaginous zone of the midpalatal suture in postnatal mice. Orthod Craniofac Res 27, 276–286. 10.1111/ocr.12724.

28. Care, A.S., Diener, K.R., Jasper, M.J., Brown, H.M., Ingman, W.V., and Robertson, S.A. (2013). Macrophages regulate corpus luteum development during embryo implantation in mice. J Clin Invest 123, 3472–3487. 10.1172/JCI60561.

29. Wang, J., Xie, D., Liu, M., Gong, Y., Shi, X., Wei, J.Y., and Quan, S. (2016). Uterine macrophages affect embryo implantation via regulating vascular endothelial growth factor A in mice. Nan fang yi ke da xue xue bao = Journal of Southern Medical University 36, 909–914.

30. Metschnikoff, E. (1891). Lecture on Phagocytosis and Immunity. Br Med J 1, 213–217. 10.1136/bmj.1.1570.213.

31. van Furth, R., Cohn, Z.A., Hirsch, J.G., Humphrey, J.H., Spector, W.G., and Langevoort, H.L. (1972). The mononuclear phagocyte system: a new classification of macrophages, monocytes, and their precursor cells. Bull World Health Organ 46, 845–852.

32. Bourez, R.L.J.H., Mathijssen, I.M.J., Vaandrager, J.M., and Vermeij-Keers, C. (1997). Apoptotic Cell Death During Normal Embryogenesis of the Coronal Suture: Early Detection of Apoptosis in Mice Using Annexin V. Journal of Craniofacial Surgery 8, 441–445.

33. Dupe, V., Ghyselinck, N.B., Thomazy, V., Nagy, L., Davies, P.J., Chambon, P., and Mark, M. (1999). Essential roles of retinoic acid signaling in interdigital apoptosis and control of BMP-7 expression in mouse autopods. Dev Biol 208, 30–43. 10.1006/dbio.1998.9176.

34. Tang, L.S., Santillano, D.R., Wlodarczyk, B.J., Miranda, R.C., and Finnell, R.H. (2005). Role of Folbp1 in the regional regulation of apoptosis and cell proliferation in the developing neural tube and craniofacies. Am J Med Genet C Semin Med Genet 135C, 48-58. 10.1002/ajmg.c.30053.

35. Zakeri, Z., Quaglino, D., and Ahuja, H.S. (1994). Apoptotic cell death in the mouse limb and its suppression in the hammertoe mutant. Dev Biol 165, 294–297. 10.1006/dbio.1994.1255.

36. Iyyanar, P.P.R., Qin, C., Adhikari, N., Liu, H., Hu, Y.C., Jiang, R., and Lan, Y. (2023). Developmental origin of the mammalian premaxilla. Dev Biol 503, 1–9. 10.1016/j.ydbio.2023.07.005.

37. Richman, J.M., and Tickle, C. (1989). Epithelia are interchangeable between facial primordia of chick embryos and morphogenesis is controlled by the mesenchyme. Dev Biol 136, 201–210. 10.1016/0012-1606(89)90142-5.

38. Wedden, S.E. (1987). Epithelial-mesenchymal interactions in the development of chick facial primordia and the target of retinoid action. Development 99, 341–351. 10.1242/dev.99.3.341.

39. Funato, N., Nakamura, M., and Yanagisawa, H. (2015). Molecular basis of cleft palates in mice. World J Biol Chem 6, 121–138. 10.4331/wjbc.v6.i3.121.

40. Iwata, J., Hacia, J.G., Suzuki, A., Sanchez-Lara, P.A., Urata, M., and Chai, Y. (2012). Modulation of noncanonical TGF-beta signaling prevents cleft palate in Tgfbr2 mutant mice. J Clin Invest 122, 873–885. 10.1172/JCI61498.

41. Sanford, L.P., Ormsby, I., Gittenberger-de Groot, A.C., Sariola, H., Friedman, R., Boivin, G.P., Cardell, E.L., and Doetschman, T. (1997). TGFbeta2 knockout mice have multiple developmental defects that are non-overlapping with other TGFbeta knockout phenotypes. Development 124, 2659–2670. 10.1242/dev.124.13.2659.

42. Zhao, Y., Guo, Y.J., Tomac, A.C., Taylor, N.R., Grinberg, A., Lee, E.J., Huang, S., and Westphal, H. (1999). Isolated cleft palate in mice with a targeted mutation of the LIM homeobox gene lhx8. Proc Natl Acad Sci U S A 96, 15002–15006. 10.1073/pnas.96.26.15002.

43. Vora, S.R. (2017). Mouse models for the study of cranial base growth and anomalies. Orthod Craniofac Res 20 *Suppl 1*, 18–25. 10.1111/ocr.12180.

44. Eswarakumar, V.P., Horowitz, M.C., Locklin, R., Morriss-Kay, G.M., and Lonai, P. (2004). A gain-of-function mutation of Fgfr2c demonstrates the roles of this receptor variant in osteogenesis. Proc Natl Acad Sci U S A 101, 12555–12560. 10.1073/pnas.0405031101.

45. Kawasaki, M., Izu, Y., Hayata, T., Ideno, H., Nifuji, A., Sheffield, V.C., Ezura, Y., and Noda, M. (2017). Bardet-Biedl syndrome 3 regulates the development of cranial base midline structures. Bone 101, 179–190. 10.1016/j.bone.2016.02.017.

46. Laurita, J., Koyama, E., Chin, B., Taylor, J.A., Lakin, G.E., Hankenson, K.D., Bartlett, S.P., and Nah, H.D. (2011). The Muenke syndrome mutation (FgfR3P244R) causes cranial base shortening associated with growth plate dysfunction and premature perichondrial ossification in murine basicranial synchondroses. Dev Dyn 240, 2584–2596. 10.1002/dvdy.22752.

47. Nagata, M., Nuckolls, G.H., Wang, X., Shum, L., Seki, Y., Kawase, T., Takahashi, K., Nonaka, K., Takahashi, I., Noman, A.A., et al. (2011). The primary site of the acrocephalic feature in Apert Syndrome is a dwarf cranial base with accelerated chondrocytic differentiation due to aberrant activation of the FGFR2 signaling. Bone 48, 847–856. 10.1016/j.bone.2010.11.014.

48. Panda, S.P., Guntur, A.R., Polusani, S.R., Fajardo, R.J., Gakunga, P.T., Roman, L.J., and Masters, B.S. (2013). Conditional deletion of cytochrome p450 reductase in osteoprogenitor cells affects long bone and skull development in mice recapitulating antley-bixler syndrome: role of a redox enzyme in development. PLoS One 8, e75638. 10.1371/journal.pone.0075638.

49. Schmidt, M., and Ondreka, N. Hydrocephalus in Animals.

50. Cohen, A.R., Leifer, D.W., Zechel, M., Flaningan, D.P., Lewin, J.S., and Lust, W.D. (1999). Characterization of a model of hydrocephalus in transgenic mice. J Neurosurg 91, 978–988. 10.3171/jns.1999.91.6.0978.

51. Kohn, D.F., Chinookoswong, N., and Chou, S.M. (1981). A new model of congenital hydrocephalus in the rat. Acta Neuropathol 54, 211–218. 10.1007/BF00687744.

52. Stottmann, R.W., Moran, J.L., Turbe-Doan, A., Driver, E., Kelley, M., and Beier, D.R. (2011). Focusing forward genetics: a tripartite ENU screen for neurodevelopmental mutations in the mouse. Genetics 188, 615–624. 10.1534/genetics.111.126862.

53. Jimenez, A.J., Tome, M., Paez, P., Wagner, C., Rodriguez, S., Fernandez-Llebrez, P., Rodriguez, E.M., and Perez-Figares, J.M. (2001). A programmed ependymal denudation precedes congenital hydrocephalus in the hyh mutant mouse. J Neuropathol Exp Neurol 60, 1105–1119. 10.1093/jnen/60.11.1105.

54. Yang, J., Simonneau, C., Kilker, R., Oakley, L., Byrne, M.D., Nichtova, Z., Stefanescu, I., Pardeep-Kumar, F., Tripathi, S., Londin, E., et al. (2019). Murine MPDZ-linked hydrocephalus is caused by hyperpermeability of the choroid plexus. EMBO Mol Med 11. 10.15252/emmm.201809540.

55. Gow, D.J., Sester, D.P., and Hume, D.A. (2010). CSF-1, IGF-1, and the control of postnatal growth and development. J Leukoc Biol 88, 475–481. 10.1189/jlb.0310158.

56. Ploeger, D.T., Hosper, N.A., Schipper, M., Koerts, J.A., de Rond, S., and Bank, R.A. (2013). Cell plasticity in wound healing: paracrine factors of M1/ M2 polarized macrophages influence the phenotypical state of dermal fibroblasts. Cell Commun Signal 11, 29. 10.1186/1478-811X-11-29.

57. Weber, K.S., Nelson, P.J., Grone, H.J., and Weber, C. (1999). Expression of CCR2 by endothelial cells: implications for MCP-1 mediated wound injury repair and In vivo inflammatory activation of endothelium. Arterioscler Thromb Vasc Biol 19, 2085–2093. 10.1161/01.atv.19.9.2085.

58. Angelozzi, M., Pellegrino da Silva, R., Gonzalez, M.V., and Lefebvre, V. (2022). Single-cell atlas of craniogenesis uncovers SOXC-dependent, highly proliferative, and myofibroblast-like osteodermal progenitors. Cell Rep 40, 111045. 10.1016/j.celrep.2022.111045.

59. Rajderkar, S.S., Paraiso, K., Amaral, M.L., Kosicki, M., Cook, L.E., Darbellay, F., Spurrell, C.H., Osterwalder, M., Zhu, Y., Wu, H., et al. (2024). Dynamic enhancer landscapes in human craniofacial development. Nat Commun 15, 2030. 10.1038/s41467-024-46396-4.

60. Biguetti, C.C., Vieira, A.E., Cavalla, F., Fonseca, A.C., Colavite, P.M., Silva, R.M., Trombone, A.P.F., and Garlet, G.P. (2018). CCR2 Contributes to F4/80+ Cells Migration Along Intramembranous Bone Healing in Maxilla, but Its Deficiency Does Not Critically Affect the Healing Outcome. Front Immunol 9, 1804. 10.3389/fimmu.2018.01804.

61. Namangkalakul, W., Nagai, S., Jin, C., Nakahama, K.I., Yoshimoto, Y., Ueha, S., Akiyoshi, K., Matsushima, K., Nakashima, T., Takechi, M., and Iseki, S. (2023). Augmented effect of fibroblast growth factor 18 in bone morphogenetic protein 2-induced calvarial bone healing by activation of CCL2/CCR2 axis on M2 macrophage polarization. J Tissue Eng 14, 20417314231187960. 10.1177/20417314231187960.

62. Xu, H., Zhang, S., Sathe, A.A., Jin, Z., Guan, J., Sun, W., Xing, C., Zhang, H., and Yan, B. (2022). CCR2(+) Macrophages Promote Orthodontic Tooth Movement and Alveolar Bone Remodeling. Front Immunol 13, 835986. 10.3389/fimmu.2022.835986.

63. Leichtle, A., Hernandez, M., Ebmeyer, J., Yamasaki, K., Lai, Y., Radek, K., Choung, Y.H., Euteneuer, S., Pak, K., Gallo, R., et al. (2010). CC chemokine ligand 3 overcomes the bacteriocidal and phagocytic defect of macrophages and hastens recovery from experimental otitis media in TNF−/− mice. J Immunol 184, 3087–3097. 10.4049/jimmunol.0901167.

64. Repeke, C.E., Ferreira, S.B., Jr., Claudino, M., Silveira, E.M., de Assis, G.F., Avila-Campos, M.J., Silva, J.S., and Garlet, G.P. (2010). Evidences of the cooperative role of the chemokines CCL3, CCL4 and CCL5 and its receptors CCR1+ and CCR5+ in RANKL+ cell migration throughout experimental periodontitis in mice. Bone 46, 1122–1130. 10.1016/j.bone.2009.12.030.

65. Takahashi, H., Tashiro, T., Miyazaki, M., Kobayashi, M., Pollard, R.B., and Suzuki, F. (2002). An essential role of macrophage inflammatory protein 1α/CCL3 on the expression of host’s innate immunities against infectious complications. Journal of Leukocyte Biology 72, 1190–1197. 10.1189/jlb.72.6.1190.

66. Wang, J., Tian, Y., Phillips, K.L., Chiverton, N., Haddock, G., Bunning, R.A., Cross, A.K., Shapiro, I.M., Le Maitre, C.L., and Risbud, M.V. (2013). Tumor necrosis factor alpha-and interleukin-1beta-dependent induction of CCL3 expression by nucleus pulposus cells promotes macrophage migration through CCR1. Arthritis Rheum 65, 832–842. 10.1002/art.37819.

67. Cheung, R., Malik, M., Ravyn, V., Tomkowicz, B., Ptasznik, A., and Collman, R.G. (2009). An arrestin-dependent multi-kinase signaling complex mediates MIP-1beta/CCL4 signaling and chemotaxis of primary human macrophages. J Leukoc Biol 86, 833–845. 10.1189/jlb.0908551.

68. von Stebut, E., Metz, M., Milon, G., Knop, J., and Maurer, M. (2003). Early macrophage influx to sites of cutaneous granuloma formation is dependent on MIP-1alpha /beta released from neutrophils recruited by mast cell-derived TNFalpha. Blood 101, 210–215. 10.1182/blood-2002-03-0921.

69. Huang, J., Yang, G., Xiong, X., Wang, M., Yuan, J., Zhang, Q., Gong, C., Qiu, Z., Meng, Z., Xu, R., et al. (2020). Age-related CCL12 Aggravates Intracerebral Hemorrhage-induced Brain Injury via Recruitment of Macrophages and T Lymphocytes. Aging Dis 11, 1103–1115. 10.14336/AD.2019.1229.

70. Ma, X., Mai, L., He, Y., Jia, S., Yang, R., Fan, W., and Huang, F. (2025). Chemokine CCL12 in trigeminal ganglion contributes to CFA-induced mechanical allodynia in mice. Neuroscience 580, 115–123. 10.1016/j.neuroscience.2025.06.026.

71. Bao, Z., Zeng, W., Zhang, D., Wang, L., Deng, X., Lai, J., Li, J., Gong, J., and Xiang, G. (2022). SNAIL Induces EMT and Lung Metastasis of Tumours Secreting CXCL2 to Promote the Invasion of M2-Type Immunosuppressed Macrophages in Colorectal Cancer. Int J Biol Sci 18, 2867–2881. 10.7150/ijbs.66854.

72. Herbold, W., Maus, R., Hahn, I., Ding, N., Srivastava, M., Christman, J.W., Mack, M., Reutershan, J., Briles, D.E., Paton, J.C., et al. (2010). Importance of CXC chemokine receptor 2 in alveolar neutrophil and exudate macrophage recruitment in response to pneumococcal lung infection. Infect Immun 78, 2620–2630. 10.1128/IAI.01169-09.

73. Nie, F., Zhang, J., Tian, H., Zhao, J., Gong, P., Wang, H., Wang, S., Yang, P., and Yang, C. (2024). The role of CXCL2-mediated crosstalk between tumor cells and macrophages in Fusobacterium nucleatum-promoted oral squamous cell carcinoma progression. Cell Death Dis 15, 277. 10.1038/s41419-024-06640-7.

74. Caetano, A.J., Redhead, Y., Karim, F., Dhami, P., Kannambath, S., Nuamah, R., Volponi, A.A., Nibali, L., Booth, V., D’Agostino, E.M., and Sharpe, P.T. (2023). Spatially resolved transcriptomics reveals pro-inflammatory fibroblast involved in lymphocyte recruitment through CXCL8 and CXCL10. Elife 12. 10.7554/eLife.81525.

75. Lee, J.H., Kim, B., Jin, W.J., Kim, H.H., Ha, H., and Lee, Z.H. (2017). Pathogenic roles of CXCL10 signaling through CXCR3 and TLR4 in macrophages and T cells: relevance for arthritis. Arthritis Res Ther 19, 163. 10.1186/s13075-017-1353-6.

76. Tsai, C.F., Chen, J.H., and Yeh, W.L. (2019). Pulmonary fibroblasts-secreted CXCL10 polarizes alveolar macrophages under pro-inflammatory stimuli. Toxicol Appl Pharmacol 380, 114698. 10.1016/j.taap.2019.114698.

77. Calvo, C.F., Yoshimura, T., Gelman, M., and Mallat, M. (1996). Production of monocyte chemotactic protein-1 by rat brain macrophages. Eur J Neurosci 8, 1725–1734. 10.1111/j.1460-9568.1996.tb01316.x.

78. Kaplanski, G., Marin, V., Montero-Julian, F., Mantovani, A., and Farnarier, C. (2003). IL-6: a regulator of the transition from neutrophil to monocyte recruitment during inflammation. Trends in Immunology 24, 25–29. 10.1016/S1471-4906(02)00013-3.

79. Wang, Z., Wu, Z., Wang, H., Feng, R., Wang, G., Li, M., Wang, S.Y., Chen, X., Su, Y., Wang, J., et al. (2023). An immune cell atlas reveals the dynamics of human macrophage specification during prenatal development. Cell 186, 4454–4471 e4419. 10.1016/j.cell.2023.08.019.

80. Lewis, A.E., Vasudevan, H.N., O’Neill, A.K., Soriano, P., and Bush, J.O. (2013). The widely used Wnt1-Cre transgene causes developmental phenotypes by ectopic activation of Wnt signaling. Dev Biol 379, 229–234. 10.1016/j.ydbio.2013.04.026.

81. Schatteman, G.C., Morrison-Graham, K., van Koppen, A., Weston, J.A., and Bowen-Pope, D.F. (1992). Regulation and role of PDGF receptor alpha-subunit expression during embryogenesis. Development 115, 123–131. 10.1242/dev.115.1.123.

82. Cai, J., Wu, Y., Mirua, T., Pierce, J.L., Lucero, M.T., Albertine, K.H., Spangrude, G.J., and Rao, M.S. (2002). Properties of a fetal multipotent neural stem cell (NEP cell). Dev Biol 251, 221–240. 10.1006/dbio.2002.0828.

83. Roellig, D., Tan-Cabugao, J., Esaian, S., and Bronner, M.E. (2017). Dynamic transcriptional signature and cell fate analysis reveals plasticity of individual neural plate border cells. Elife 6. 10.7554/eLife.21620.

84. Kim, J., Lo, L., Dormand, E., and Anderson, D.J. (2003). SOX10 maintains multipotency and inhibits neuronal differentiation of neural crest stem cells. Neuron 38, 17–31. 10.1016/s0896-6273(03)00163-6.

85. Southard-Smith, E.M., Kos, L., and Pavan, W.J. (1998). Sox10 mutation disrupts neural crest development in Dom Hirschsprung mouse model. Nat Genet 18, 60–64. 10.1038/ng0198-60.

86. Bosch, A.J.T., Keller, L., Steiger, L., Rohm, T.V., Wiedemann, S.J., Low, A.J.Y., Stawiski, M., Rachid, L., Roux, J., Konrad, D., et al. (2023). CSF1R inhibition with PLX5622 affects multiple immune cell compartments and induces tissue-specific metabolic effects in lean mice. Diabetologia 66, 2292–2306. 10.1007/s00125-023-06007-1.

87. Elmore, M.R.P., Hohsfield, L.A., Kramar, E.A., Soreq, L., Lee, R.J., Pham, S.T., Najafi, A.R., Spangenberg, E.E., Wood, M.A., West, B.L., and Green, K.N. (2018). Replacement of microglia in the aged brain reverses cognitive, synaptic, and neuronal deficits in mice. Aging Cell 17, e12832. 10.1111/acel.12832.

88. Hennen, E.M., Uppuganti, S., de la Visitacion, N., Chen, W., Krishnan, J., Vecchi Iii, L.A., Patrick, D.M., Siedlinski, M., Lemoli, M., Delgado, R., et al. (2025). Hypertension promotes bone loss and fragility by favoring bone resorption in mouse models. J Clin Invest. 10.1172/JCI184325.

89. Lei, F., Cui, N., Zhou, C., Chodosh, J., Vavvas, D.G., and Paschalis, E.I. (2020). CSF1R inhibition by a small-molecule inhibitor is not microglia specific; affecting hematopoiesis and the function of macrophages. Proc Natl Acad Sci U S A 117, 23336–23338. 10.1073/pnas.1922788117.

90. Spangenberg, E., Severson, P.L., Hohsfield, L.A., Crapser, J., Zhang, J., Burton, E.A., Zhang, Y., Spevak, W., Lin, J., Phan, N.Y., et al. (2019). Sustained microglial depletion with CSF1R inhibitor impairs parenchymal plaque development in an Alzheimer’s disease model. Nat Commun 10, 3758. 10.1038/s41467-019-11674-z.

91. de Luis, M., Xu, S., and Zinn, K. (2025). Fluorescent labeling of proteins in vitro and in vivo using encoded peptide tags. J Biol Chem 301, 110229. 10.1016/j.jbc.2025.110229.

92. Snapp, E. (2005). Design and use of fluorescent fusion proteins in cell biology. Curr Protoc Cell Biol Chapter 21, 21 24 21–21 24 13. 10.1002/0471143030.cb2104s27.

93. Stadler, C., Rexhepaj, E., Singan, V.R., Murphy, R.F., Pepperkok, R., Uhlen, M., Simpson, J.C., and Lundberg, E. (2013). Immunofluorescence and fluorescent-protein tagging show high correlation for protein localization in mammalian cells. Nat Methods 10, 315–323. 10.1038/nmeth.2377.

94. Silver, J., and Hughes, A.F. (1974). The relationship between morphogenetic cell death and the development of congenital anophthalmia. J Comp Neurol 157, 281–301. 10.1002/cne.901570303.

95. Makishima, T., Hochman, L., Armstrong, P., Rosenberger, E., Ridley, R., Woo, M., Perachio, A., and Wood, S. (2011). Inner ear dysfunction in caspase-3 deficient mice. BMC Neurosci 12, 102. 10.1186/1471-2202-12-102.

96. Wadu, S.G., Penhall, B., and Townsend, G.C. (1997). Morphological variability of the human inferior alveolar nerve. Clinical Anatomy 10, 82–87. 10.1002/(sici)1098-2353(1997)10:2<82::Aid-ca2>3.0.Co;2-v.

97. Yamamoto, M., Takada, H., Ishizuka, S., Kitamura, K., Jeong, J., Sato, M., Hinata, N., and Abe, S. (2020). Morphological association between the muscles and bones in the craniofacial region. PLoS One 15, e0227301. 10.1371/journal.pone.0227301.

98. Shen, H., Grimston, S., Civitelli, R., and Thomopoulos, S. (2015). Deletion of connexin43 in osteoblasts/osteocytes leads to impaired muscle formation in mice. J Bone Miner Res 30, 596–605. 10.1002/jbmr.2389.

99. Kitai, N., Fujii, Y., Murakami, S., Furukawa, S., Kreiborg, S., and Takada, K. (2002). Human masticatory muscle volume and zygomatico-mandibular form in adults with mandibular prognathism. J Dent Res 81, 752–756. 10.1177/0810752.

100. Yamamoto, M., Ho Cho, K., Murakami, G., Abe, S., and Rodriguez-Vazquez, J.F. (2018). Early Fetal Development of the Otic and Pterygopalatine Ganglia with Special Reference to the Topographical Relationship with the Developing Sphenoid Bone. Anat Rec (Hoboken) 301, 1442–1453. 10.1002/ar.23833.

101. Toth, M., Sirirattanapan, J., and Mann, W. (2013). Patterns of anomalies of structures of the middle ear and the facial nerve as revealed in newborn temporal bones. Otol Neurotol 34, 1121–1126. 10.1097/MAO.0b013e318283987f.

102. Iyyanar, P.P.R., Wu, Z., Lan, Y., Hu, Y.C., and Jiang, R. (2022). Alx1 Deficient Mice Recapitulate Craniofacial Phenotype and Reveal Developmental Basis of ALX1-Related Frontonasal Dysplasia. Front Cell Dev Biol 10, 777887. 10.3389/fcell.2022.777887.

103. Hume, D.A., Teakle, N., Keshvari, S., and Irvine, K.M. (2023). Macrophage deficiency in CSF1R-knockout rat embryos does not compromise placental or embryo development. J Leukoc Biol 114, 421–433. 10.1093/jleuko/qiad052.

104. Dai, X.-M., Zong, X.-H., Akhter, M.P., and Stanley, E.R. (2004). Osteoclast Deficiency Results in Disorganized Matrix, Reduced Mineralization, and Abnormal Osteoblast Behavior in Developing Bone. Journal of Bone and Mineral Research 19, 1441–1451. 10.1359/JBMR.040514.

105. Harris, S.E., MacDougall, M., Horn, D., Woodruff, K., Zimmer, S.N., Rebel, V.I., Fajardo, R., Feng, J.Q., Gluhak-Heinrich, J., Harris, M.A., and Abboud Werner, S. (2012). Meox2Cre-mediated disruption of CSF-1 leads to osteopetrosis and osteocyte defects. Bone 50, 42–53. 10.1016/j.bone.2011.09.038.

106. Ryan, G.R., Dai, X.M., Dominguez, M.G., Tong, W., Chuan, F., Chisholm, O., Russell, R.G., Pollard, J.W., and Stanley, E.R. (2001). Rescue of the colony-stimulating factor 1 (CSF-1)-nullizygous mouse (Csf1(op)/Csf1(op)) phenotype with a CSF-1 transgene and identification of sites of local CSF-1 synthesis. Blood 98, 74–84. 10.1182/blood.v98.1.74.

107. Lawrence, A.R., Canzi, A., Bridlance, C., Olivie, N., Lansonneur, C., Catale, C., Pizzamiglio, L., Kloeckner, B., Silvin, A., Munro, D.A.D., et al. (2024). Microglia maintain structural integrity during fetal brain morphogenesis. Cell 187, 962–980 e919. 10.1016/j.cell.2024.01.012.

108. Rojo, R., Raper, A., Ozdemir, D.D., Lefevre, L., Grabert, K., Wollscheid-Lengeling, E., Bradford, B., Caruso, M., Gazova, I., Sanchez, A., et al. (2019). Deletion of a Csf1r enhancer selectively impacts CSF1R expression and development of tissue macrophage populations. Nat Commun 10, 3215. 10.1038/s41467-019-11053-8.

109. Beerepoot, S., Verbeke, J., Plantinga, M., Nierkens, S., Pouwels, P.J.W., Wolf, N.I., Simons, C., and van der Knaap, M.S. (2024). Leukoencephalopathy with calcifications, developmental brain abnormalities and skeletal dysplasia due to homozygosity for a hypomorphic CSF1R variant: A report of three siblings. Am J Med Genet A 194, e63800. 10.1002/ajmg.a.63800.

110. Jiang, J., Li, W., Wang, X., Du, Z., Chen, J., Liu, Y., Li, W., Lu, Z., Wang, Y., and Xu, J. (2022). Two Novel Intronic Mutations in the CSF1R Gene in Two Families With CSF1R-Microglial Encephalopathy. Front Cell Dev Biol 10, 902067. 10.3389/fcell.2022.902067.

111. Kindis, E., Simsek-Kiper, P.O., Kosukcu, C., Taskiran, E.Z., Gocmen, R., Utine, E., Haliloglu, G., Boduroglu, K., and Alikasifoglu, M. (2021). Further expanding the mutational spectrum of brain abnormalities, neurodegeneration, and dysosteosclerosis: A rare disorder with neurologic regression and skeletal features. Am J Med Genet A 185, 1888–1896. 10.1002/ajmg.a.62179.

112. Oosterhof, N., Chang, I.J., Karimiani, E.G., Kuil, L.E., Jensen, D.M., Daza, R., Young, E., Astle, L., van der Linde, H.C., Shivaram, G.M., et al. (2019). Homozygous Mutations in CSF1R Cause a Pediatric-Onset Leukoencephalopathy and Can Result in Congenital Absence of Microglia. The American Journal of Human Genetics 104, 936–947. 10.1016/j.ajhg.2019.03.010.

113. Oosterhof, N., Kuil, L.E., van der Linde, H.C., Burm, S.M., Berdowski, W., van Ijcken, W.F.J., van Swieten, J.C., Hol, E.M., Verheijen, M.H.G., and van Ham, T.J. (2018). Colony-Stimulating Factor 1 Receptor (CSF1R) Regulates Microglia Density and Distribution, but Not Microglia Differentiation In Vivo. Cell Rep 24, 1203–1217 e1206. 10.1016/j.celrep.2018.06.113.

114. Tamhankar, P.M., Zhu, B., Tamhankar, V.P., Mithbawkar, S., Seabra, L., Livingston, J.H., Ikeuchi, T., and Crow, Y.J. (2020). A Novel Hypomorphic CSF1R Gene Mutation in the Biallelic State Leading to Fatal Childhood Neurodegeneration. Neuropediatrics 51, 302–306. 10.1055/s-0040-1702161.

115. Daghagh, H., Rahbar Kafshboran, H., Daneshmandpour, Y., Nasiri Aghdam, M., Talebian, S., Nouri Nojadeh, J., Hamzeiy, H., Biskup, S., and Sakhinia, E. (2023). Homozygous mutation in CSF1R causes brain abnormalities, neurodegeneration, and dysosteosclerosis (BANDDOS). Bioimpacts 13, 183–190. 10.34172/bi.2022.23528.

116. Dulski, J., Souza, J., Santos, M.L., and Wszolek, Z.K. (2023). Brain abnormalities, neurodegeneration, and dysosteosclerosis (BANDDOS): new cases, systematic literature review, and associations with CSF1R-ALSP. Orphanet J Rare Dis 18, 160. 10.1186/s13023-023-02772-9.

117. Inoue, K., Qin, Y., Xia, Y., Han, J., Yuan, R., Sun, J., Xu, R., Jiang, J.X., Greenblatt, M.B., and Zhao, B. (2023). Bone marrow Adipoq-lineage progenitors are a major cellular source of M-CSF that dominates bone marrow macrophage development, osteoclastogenesis, and bone mass. Elife 12. 10.7554/eLife.82118.

118. Nandi, S., Akhter, M.P., Seifert, M.F., Dai, X.M., and Stanley, E.R. (2006). Developmental and functional significance of the CSF-1 proteoglycan chondroitin sulfate chain. Blood 107, 786–795. 10.1182/blood-2005-05-1822.

119. Batoon, L., Keshvari, S., Irvine, K.M., Ho, E., Caruso, M., Patkar, O.L., Sehgal, A., Millard, S.M., Hume, D.A., and Pettit, A.R. (2024). Relative contributions of osteal macrophages and osteoclasts to postnatal bone development in CSF1R-deficient rats and phenotype rescue following wild-type bone marrow cell transfer. J Leukoc Biol 116, 753–765. 10.1093/jleuko/qiae077.

120. Batoon, L., Millard, S.M., Wullschleger, M.E., Preda, C., Wu, A.C., Kaur, S., Tseng, H.W., Hume, D.A., Levesque, J.P., Raggatt, L.J., and Pettit, A.R. (2019). CD169(+) macrophages are critical for osteoblast maintenance and promote intramembranous and endochondral ossification during bone repair. Biomaterials 196, 51–66. 10.1016/j.biomaterials.2017.10.033.

121. Alexander, K.A., Chang, M.K., Maylin, E.R., Kohler, T., Muller, R., Wu, A.C., Van Rooijen, N., Sweet, M.J., Hume, D.A., Raggatt, L.J., and Pettit, A.R. (2011). Osteal macrophages promote in vivo intramembranous bone healing in a mouse tibial injury model. J Bone Miner Res 26, 1517–1532. 10.1002/jbmr.354.

122. Blumer, M.J., Longato, S., and Fritsch, H. (2008). Localization of tartrate-resistant acid phosphatase (TRAP), membrane type-1 matrix metalloproteinases (MT1-MMP) and macrophages during early endochondral bone formation. J Anat 213, 431–441. 10.1111/j.1469-7580.2008.00958.x.

123. Chang, M.K., Raggatt, L.J., Alexander, K.A., Kuliwaba, J.S., Fazzalari, N.L., Schroder, K., Maylin, E.R., Ripoll, V.M., Hume, D.A., and Pettit, A.R. (2008). Osteal tissue macrophages are intercalated throughout human and mouse bone lining tissues and regulate osteoblast function in vitro and in vivo. J Immunol 181, 1232–1244. 10.4049/jimmunol.181.2.1232.

124. Batoon, L., Koh, A.J., Millard, S.M., Grewal, J., Choo, F.M., Kannan, R., Kinnaird, A., Avey, M., Teslya, T., Pettit, A.R., et al. (2024). Induction of osteoblast apoptosis stimulates macrophage efferocytosis and paradoxical bone formation. Bone Res 12, 43. 10.1038/s41413-024-00341-9.

125. Mun, S.H., Jastrzebski, S., Kalinowski, J., Zeng, S., Oh, B., Bae, S., Eugenia, G., Khan, N.M., Drissi, H., Zhou, P., et al. (2021). Sexual Dimorphism in Differentiating Osteoclast Precursors Demonstrates Enhanced Inflammatory Pathway Activation in Female Cells. J Bone Miner Res 36, 1104–1116. 10.1002/jbmr.4270.

126. Herbert, B.A., Valerio, M.S., Gaestel, M., and Kirkwood, K.L. (2015). Sexual Dimorphism in MAPK-Activated Protein Kinase-2 (MK2) Regulation of RANKL-Induced Osteoclastogenesis in Osteoclast Progenitor Subpopulations. PLoS One 10, e0125387. 10.1371/journal.pone.0125387.

127. Paglia, D.N., Yang, X., Kalinowski, J., Jastrzebski, S., Drissi, H., and Lorenzo, J. (2016). Runx1 Regulates Myeloid Precursor Differentiation Into Osteoclasts Without Affecting Differentiation Into Antigen Presenting or Phagocytic Cells in Both Males and Females. Endocrinology 157, 3058–3069. 10.1210/en.2015-2037.

128. Cenci, S., Weitzmann, M.N., Roggia, C., Namba, N., Novack, D., Woodring, J., and Pacifici, R. (2000). Estrogen deficiency induces bone loss by enhancing T-cell production of TNF-alpha. J Clin Invest 106, 1229–1237. 10.1172/JCI11066.

129. Kimble, R.B., Srivastava, S., Ross, F.P., Matayoshi, A., and Pacifici, R. (1996). Estrogen deficiency increases the ability of stromal cells to support murine osteoclastogenesis via an interleukin-1and tumor necrosis factor-mediated stimulation of macrophage colony-stimulating factor production. J Biol Chem 271, 28890–28897. 10.1074/jbc.271.46.28890.

130. Srivastava, S., Weitzmann, M.N., Kimble, R.B., Rizzo, M., Zahner, M., Milbrandt, J., Ross, F.P., and Pacifici, R. (1998). Estrogen blocks M-CSF gene expression and osteoclast formation by regulating phosphorylation of Egr-1 and its interaction with Sp-1. J Clin Invest 102, 1850–1859. 10.1172/JCI4561.

131. Przystanska, A., Rewekant, A., Sroka, A., Gedrange, T., Ekkert, M., Jonczyk-Potoczna, K., and Czajka-Jakubowska, A. (2020). Sexual dimorphism of maxillary sinuses in children and adolescents – A retrospective CT study. Ann Anat 229, 151437. 10.1016/j.aanat.2019.151437.

132. Hall, B.K. (1988). The Embryonic Development of Bone. American Scientist 76, 174–181.

133. Marvaso, V., and Bernard, G.W. (1977). Initial intramembraneous osteogenesis in vitro. Am J Anat 149, 453–468. 10.1002/aja.1001490403.

134. Sandberg, M.M. (1991). Matrix in cartilage and bone development: current views on the function and regulation of major organic components. Ann Med 23, 207–217. 10.3109/07853899109148050.

135. Takarada, T., Nakazato, R., Tsuchikane, A., Fujikawa, K., Iezaki, T., Yoneda, Y., and Hinoi, E. (2016). Genetic analysis of Runx2 function during intramembranous ossification. Development 143, 211–218. 10.1242/dev.128793.

136. Zimmermann, B. (1992). Degeneration of osteoblasts involved in intramembranous ossification of fetal rat calvaria. Cell Tissue Res 267, 75–84. 10.1007/BF00318693.

137. Hall, B.K., and Miyake, T. (2000). All for one and one for all: condensations and the initiation of skeletal development. BioEssays 22, 138–147. 10.1002/(sici)1521-1878(200002)22:2<138::Aid-bies5>3.0.Co;2-4.

138. Mallo, M. (1998). Embryological and genetic aspects of middle ear development. Int. J. Dev. Biol. 42, 11–22.

139. Miyake, T., Cameron, A.M., and Hall, B.K. (1996). Stage-specific onset of condensation and matrix deposition for Meckel’s and other first arch cartilages in inbred C57BL/6 mice. J Craniofac Genet Dev Biol 16, 32–47.

140. Wood, J.L., Hughes, A.J., Mercer, K.J., and Chapman, S.C. (2010). Analysis of chick (Gallus gallus) middle ear columella formation. BMC Dev Biol 10, 16. 10.1186/1471-213X-10-16.

141. Chai, Y., Jiang, X., Ito, Y., Bringas, P., Jr., Han, J., Rowitch, D.H., Soriano, P., McMahon, A.P., and Sucov, H.M. (2000). Fate of the mammalian cranial neural crest during tooth and mandibular morphogenesis. Development 127, 1671–1679. 10.1242/dev.127.8.1671.

142. Jiang, X., Iseki, S., Maxson, R.E., Sucov, H.M., and Morriss-Kay, G.M. (2002). Tissue origins and interactions in the mammalian skull vault. Dev Biol 241, 106–116. 10.1006/dbio.2001.0487.

143. Jaenisch, R. (1985). Mammalian neural crest cells participate in normal embryonic development on microinjection into post-implantation mouse embryos. Nature 318, 181–183. 10.1038/318181a0.

144. Tan, S.S., and Morriss-Kay, G.M. (1986). Analysis of cranial neural crest cell migration and early fates in postimplantation rat chimaeras. Development 98, 21–58. 10.1242/dev.98.1.21.

145. Yoshida, T., Vivatbutsiri, P., Morriss-Kay, G., Saga, Y., and Iseki, S. (2008). Cell lineage in mammalian craniofacial mesenchyme. Mech Dev 125, 797–808. 10.1016/j.mod.2008.06.007.

146. Bildsoe, H., Loebel, D.A., Jones, V.J., Hor, A.C., Braithwaite, A.W., Chen, Y.T., Behringer, R.R., and Tam, P.P. (2013). The mesenchymal architecture of the cranial mesoderm of mouse embryos is disrupted by the loss of Twist1 function. Dev Biol 374, 295–307. 10.1016/j.ydbio.2012.12.004.

147. Tamura, Y., Matsumura, K., Sano, M., Tabata, H., Kimura, K., Ieda, M., Arai, T., Ohno, Y., Kanazawa, H., Yuasa, S., et al. (2011). Neural crest-derived stem cells migrate and differentiate into cardiomyocytes after myocardial infarction. Arterioscler Thromb Vasc Biol 31, 582–589. 10.1161/ATVBAHA.110.214726.

148. Kamalakar, A., McKinney, J.M., Salinas Duron, D., Amanso, A.M., Ballestas, S.A., Drissi, H., Willett, N.J., Bhattaram, P., Garcia, A.J., Wood, L.B., and Goudy, S.L. (2021). JAGGED1 stimulates cranial neural crest cell osteoblast commitment pathways and bone regeneration independent of canonical NOTCH signaling. Bone 143, 115657. 10.1016/j.bone.2020.115657.

149. Hu, Y., Wang, L., Zhao, Z., Lu, W., Fan, J., Gao, B., Luo, Z., Jie, Q., Shi, X., and Yang, L. (2020). Cytokines CCL2 and CXCL1 may be potential novel predictors of early bone loss. Mol Med Rep 22, 4716–4724. 10.3892/mmr.2020.11543.

150. Miyamoto, K., Ninomiya, K., Sonoda, K.H., Miyauchi, Y., Hoshi, H., Iwasaki, R., Miyamoto, H., Yoshida, S., Sato, Y., Morioka, H., et al. (2009). MCP-1 expressed by osteoclasts stimulates osteoclastogenesis in an autocrine/paracrine manner. Biochem Biophys Res Commun 383, 373–377. 10.1016/j.bbrc.2009.04.020.

151. Sul, O.J., Ke, K., Kim, W.K., Kim, S.H., Lee, S.C., Kim, H.J., Kim, S.Y., Suh, J.H., and Choi, H.S. (2012). Absence of MCP-1 leads to elevated bone mass via impaired actin ring formation. J Cell Physiol 227, 1619–1627. 10.1002/jcp.22879.

152. Tamasi, J.A., Vasilov, A., Shimizu, E., Benton, N., Johnson, J., Bitel, C.L., Morrison, N., and Partridge, N.C. (2013). Monocyte chemoattractant protein-1 is a mediator of the anabolic action of parathyroid hormone on bone. J Bone Miner Res 28, 1975–1986. 10.1002/jbmr.1933.

153. Hardaway, A.L., Herroon, M.K., Rajagurubandara, E., and Podgorski, I. (2015). Marrow adipocyte-derived CXCL1 and CXCL2 contribute to osteolysis in metastatic prostate cancer. Clin Exp Metastasis 32, 353–368. 10.1007/s10585-015-9714-5.

154. Onan, D., Allan, E.H., Quinn, J.M., Gooi, J.H., Pompolo, S., Sims, N.A., Gillespie, M.T., and Martin, T.J. (2009). The chemokine Cxcl1 is a novel target gene of parathyroid hormone (PTH)/PTH-related protein in committed osteoblasts. Endocrinology 150, 2244–2253. 10.1210/en.2008-1597.

155. Hoshino, A., Iimura, T., Ueha, S., Hanada, S., Maruoka, Y., Mayahara, M., Suzuki, K., Imai, T., Ito, M., Manome, Y., et al. (2010). Deficiency of chemokine receptor CCR1 causes osteopenia due to impaired functions of osteoclasts and osteoblasts. J Biol Chem 285, 28826–28837. 10.1074/jbc.M109.099424.

156. Lee, D., Shin, K.J., Kim, D.W., Yoon, K.A., Choi, Y.J., Lee, B.N.R., and Cho, J.Y. (2018). CCL4 enhances preosteoclast migration and its receptor CCR5 downregulation by RANKL promotes osteoclastogenesis. Cell Death Dis 9, 495. 10.1038/s41419-018-0562-5.

157. Taddei, S.R., Queiroz-Junior, C.M., Moura, A.P., Andrade, I., Jr., Garlet, G.P., Proudfoot, A.E., Teixeira, M.M., and da Silva, T.A. (2013). The effect of CCL3 and CCR1 in bone remodeling induced by mechanical loading during orthodontic tooth movement in mice. Bone 52, 259–267. 10.1016/j.bone.2012.09.036.

158. Wintges, K., Beil, F.T., Albers, J., Jeschke, A., Schweizer, M., Claass, B., Tiegs, G., Amling, M., and Schinke, T. (2013). Impaired bone formation and increased osteoclastogenesis in mice lacking chemokine (C-C motif) ligand 5 (Ccl5). J Bone Miner Res 28, 2070–2080. 10.1002/jbmr.1937.

159. Yu, D., Zhang, S., Ma, C., Huang, S., Xu, L., Liang, J., Li, H., Fan, Q., Liu, G., and Zhai, Z. (2023). CCL3 in the bone marrow microenvironment causes bone loss and bone marrow adiposity in aged mice. JCI Insight 8. 10.1172/jci.insight.159107.

160. Araujo-Pires, A.C., Vieira, A.E., Francisconi, C.F., Biguetti, C.C., Glowacki, A., Yoshizawa, S., Campanelli, A.P., Trombone, A.P., Sfeir, C.S., Little, S.R., and Garlet, G.P. (2015). IL-4/CCL22/CCR4 axis controls regulatory T-cell migration that suppresses inflammatory bone loss in murine experimental periodontitis. J Bone Miner Res 30, 412–422. 10.1002/jbmr.2376.

161. Ha, J., Choi, H.S., Lee, Y., Kwon, H.J., Song, Y.W., and Kim, H.H. (2010). CXC chemokine ligand 2 induced by receptor activator of NF-kappa B ligand enhances osteoclastogenesis. J Immunol 184, 4717–4724. 10.4049/jimmunol.0902444.

162. Ha, J., Lee, Y., and Kim, H.H. (2011). CXCL2 mediates lipopolysaccharide-induced osteoclastogenesis in RANKL-primed precursors. Cytokine 55, 48–55. 10.1016/j.cyto.2011.03.026.

163. Ma, C., Gao, J., Liang, J., Wang, F., Xu, L., Bu, J., He, B., Liu, G., Niu, R., and Liu, G. (2023). CCL12 induces trabecular bone loss by stimulating RANKL production in BMSCs during acute lung injury. Exp Mol Med 55, 818–830. 10.1038/s12276-023-00970-w.

164. Yang, Y., Zhou, X., Li, Y., Chen, A., Liang, W., Liang, G., Huang, B., Li, Q., and Jin, D. (2019). CXCL2 attenuates osteoblast differentiation by inhibiting the ERK1/2 signaling pathway. J Cell Sci 132. 10.1242/jcs.230490.

165. Yano, S., Mentaverri, R., Kanuparthi, D., Bandyopadhyay, S., Rivera, A., Brown, E.M., and Chattopadhyay, N. (2005). Functional expression of beta-chemokine receptors in osteoblasts: role of regulated upon activation, normal T cell expressed and secreted (RANTES) in osteoblasts and regulation of its secretion by osteoblasts and osteoclasts. Endocrinology 146, 2324–2335. 10.1210/en.2005-0065.

166. Kwak, H.B., Ha, H., Kim, H.N., Lee, J.H., Kim, H.S., Lee, S., Kim, H.M., Kim, J.Y., Kim, H.H., Song, Y.W., and Lee, Z.H. (2008). Reciprocal cross-talk between RANKL and interferon-gamma-inducible protein 10 is responsible for bone-erosive experimental arthritis. Arthritis Rheum 58, 1332–1342. 10.1002/art.23372.

167. Marahleh, A., Kitaura, H., Ohori, F., Kishikawa, A., Ogawa, S., Shen, W.R., Qi, J., Noguchi, T., Nara, Y., and Mizoguchi, I. (2019). TNF-alpha Directly Enhances Osteocyte RANKL Expression and Promotes Osteoclast Formation. Front Immunol 10, 2925. 10.3389/fimmu.2019.02925.

168. Takahashi, T., Wada, T., Mori, M., Kokai, Y., and Ishii, S. (1996). Overexpression of the granulocyte colony-stimulating factor gene leads to osteoporosis in mice. Lab Invest 74, 827–834.

169. Takayanagi, H., Kim, S., Matsuo, K., Suzuki, H., Suzuki, T., Sato, K., Yokochi, T., Oda, H., Nakamura, K., Ida, N., et al. (2002). RANKL maintains bone homeostasis through c-Fos-dependent induction of interferon-beta. Nature 416, 744–749. 10.1038/416744a.

170. Yu, H., Zhang, T., Lu, H., Ma, Q., Zhao, D., Sun, J., and Wang, Z. (2021). Granulocyte colony-stimulating factor (G-CSF) mediates bone resorption in periodontitis. BMC Oral Health 21, 299. 10.1186/s12903-021-01658-1.

171. Zhang, Y.H., Heulsmann, A., Tondravi, M.M., Mukherjee, A., and Abu-Amer, Y. (2001). Tumor necrosis factor-alpha (TNF) stimulates RANKL-induced osteoclastogenesis via coupling of TNF type 1 receptor and RANK signaling pathways. J Biol Chem 276, 563–568. 10.1074/jbc.M008198200.

172. Zhao, R., Chen, N.N., Zhou, X.W., Miao, P., Hu, C.Y., Qian, L., Yu, Q.W., Zhang, J.Y., Nie, H., Chen, X.H., et al. (2014). Exogenous IFN-beta regulates the RANKL-c-Fos-IFN-beta signaling pathway in the collagen antibody-induced arthritis model. J Transl Med 12, 330. 10.1186/s12967-014-0330-y.

173. Kaneshiro, S., Ebina, K., Shi, K., Higuchi, C., Hirao, M., Okamoto, M., Koizumi, K., Morimoto, T., Yoshikawa, H., and Hashimoto, J. (2014). IL-6 negatively regulates osteoblast differentiation through the SHP2/MEK2 and SHP2/Akt2 pathways in vitro. J Bone Miner Metab 32, 378–392. 10.1007/s00774-013-0514-1.

174. Kitamura, H., Kawata, H., Takahashi, F., Higuchi, Y., Furuichi, T., and Ohkawa, H. (1995). Bone marrow neutrophilia and suppressed bone turnover in human interleukin-6 transgenic mice. A cellular relationship among hematopoietic cells, osteoblasts, and osteoclasts mediated by stromal cells in bone marrow. Am J Pathol 147, 1682–1692.

175. Liu, H., Feng, W., Yimin Cui, J., Lv, S., Hasegawa, T., Sun, B., Li, J., Oda, K., Amizuka, N., and Li, M. (2014). Histological Evidence of Increased Osteoclast Cell Number and Asymmetric Bone Resorption Activity in the Tibiae of Interleukin-6-Deficient Mice. J Histochem Cytochem 62, 556–564. 10.1369/0022155414537830.

176. Palmqvist, P., Persson, E., Conaway, H.H., and Lerner, U.H. (2002). IL-6, leukemia inhibitory factor, and oncostatin M stimulate bone resorption and regulate the expression of receptor activator of NF-kappa B ligand, osteoprotegerin, and receptor activator of NF-kappa B in mouse calvariae. J Immunol 169, 3353–3362. 10.4049/jimmunol.169.6.3353.

177. Poli, V., Balena, R., Fattori, E., Markatos, A., Yamamoto, M., Tanaka, H., Ciliberto, G., Rodan, G.A., and Costantini, F. (1994). Interleukin-6 deficient mice are protected from bone loss caused by estrogen depletion. EMBO J 13, 1189–1196. 10.1002/j.1460-2075.1994.tb06368.x.

178. Hayden, J.M., Mohan, S., and Baylink, D.J. (1995). The insulin-like growth factor system and the coupling of formation to resorption. Bone 17, 93S–98S. 10.1016/8756-3282(95)00186-h.

179. Howard, G.A., Bottemiller, B.L., Turner, R.T., Rader, J.I., and Baylink, D.J. (1981). Parathyroid hormone stimulates bone formation and resorption in organ culture: evidence for a coupling mechanism. Proc Natl Acad Sci U S A 78, 3204–3208. 10.1073/pnas.78.5.3204.

180. Nakamura, M., Udagawa, N., Matsuura, S., Mogi, M., Nakamura, H., Horiuchi, H., Saito, N., Hiraoka, B.Y., Kobayashi, Y., Takaoka, K., et al. (2003). Osteoprotegerin regulates bone formation through a coupling mechanism with bone resorption. Endocrinology 144, 5441–5449. 10.1210/en.2003-0717.

181. Tang, Y., Wu, X., Lei, W., Pang, L., Wan, C., Shi, Z., Zhao, L., Nagy, T.R., Peng, X., Hu, J., et al. (2009). TGF-beta1-induced migration of bone mesenchymal stem cells couples bone resorption with formation. Nat Med 15, 757–765. 10.1038/nm.1979.

182. Zhao, C., Irie, N., Takada, Y., Shimoda, K., Miyamoto, T., Nishiwaki, T., Suda, T., and Matsuo, K. (2006). Bidirectional ephrinB2-EphB4 signaling controls bone homeostasis. Cell Metab 4, 111–121. 10.1016/j.cmet.2006.05.012.

183. Hu, Z.L., Shi, M., Huang, Y., Zheng, M.H., Pei, Z., Chen, J.Y., Han, H., and Ding, Y.Q. (2011). The role of the transcription factor Rbpj in the development of dorsal root ganglia. Neural Dev 6, 14. 10.1186/1749-8104-6-14.

184. Ngan, E.S., Garcia-Barcelo, M.M., Yip, B.H., Poon, H.C., Lau, S.T., Kwok, C.K., Sat, E., Sham, M.H., Wong, K.K., Wainwright, B.J., et al. (2011). Hedgehog/Notch-induced premature gliogenesis represents a new disease mechanism for Hirschsprung disease in mice and humans. J Clin Invest 121, 3467–3478. 10.1172/JCI43737.

185. Rabbani, P., Takeo, M., Chou, W., Myung, P., Bosenberg, M., Chin, L., Taketo, M.M., and Ito, M. (2011). Coordinated activation of Wnt in epithelial and melanocyte stem cells initiates pigmented hair regeneration. Cell 145, 941–955. 10.1016/j.cell.2011.05.004.

186. Profaci, C.P., Harvey, S.S., Bajc, K., Zhang, T.Z., Jeffrey, D.A., Zhang, A.Z., Nemec, K.M., Davtyan, H., O’Brien, C.A., McKinsey, G.L., et al. (2024). Microglia are not necessary for maintenance of blood-brain barrier properties in health, but PLX5622 alters brain endothelial cholesterol metabolism. Neuron 112, 2910–2921 e2917. 10.1016/j.neuron.2024.07.015.

187. Faure, L., Wang, Y., Kastriti, M.E., Fontanet, P., Cheung, K.K.Y., Petitpre, C., Wu, H., Sun, L.L., Runge, K., Croci, L., et al. (2020). Single cell RNA sequencing identifies early diversity of sensory neurons forming via bi-potential intermediates. Nat Commun 11, 4175. 10.1038/s41467-020-17929-4.

188. Hudacova, E., Abaffy, P., Kaplan, M.M., Krausova, M., Kubista, M., and Machon, O. (2025). Single-cell transcriptomic resolution of osteogenesis during craniofacial morphogenesis. Bone 190, 117297. 10.1016/j.bone.2024.117297.

189. Luo, R., Gao, J., Wehrle-Haller, B., and Henion, P.D. (2003). Molecular identification of distinct neurogenic and melanogenic neural crest sublineages. Development 130, 321–330. 10.1242/dev.00213.

190. Motohashi, T., Kitagawa, D., Watanabe, N., Wakaoka, T., and Kunisada, T. (2014). Neural crest-derived cells sustain their multipotency even after entry into their target tissues. Dev Dyn 243, 368–380. 10.1002/dvdy.24072.

191. Motohashi, T., Yamanaka, K., Chiba, K., Miyajima, K., Aoki, H., Hirobe, T., and Kunisada, T. (2011). Neural crest cells retain their capability for multipotential differentiation even after lineage-restricted stages. Dev Dyn 240, 1681–1693. 10.1002/dvdy.22658.

192. Wilson, Y.M., Richards, K.L., Ford-Perriss, M.L., Panthier, J.J., and Murphy, M. (2004). Neural crest cell lineage segregation in the mouse neural tube. Development 131, 6153–6162. 10.1242/dev.01533.

193. Yamane, T., Hayashi, S.-I., Mizoguchi, M., Yamazaki, H., and Kunisada, T. (1999). Derivation of melanocytes from embryonic stem cells in culture. Developmental Dynamics 216, 450–458. 10.1002/(sici)1097-0177(199912)216:4/5<450::Aid-dvdy13>3.0.Co;2-0.

194. McFarlane, L., Truong, V., Palmer, J.S., and Wilhelm, D. (2013). Novel PCR assay for determining the genetic sex of mice. Sex Dev 7, 207–211. 10.1159/000348677.

195. Tiscornia, G., Singer, O., Ikawa, M., and Verma, I.M. (2003). A general method for gene knockdown in mice by using lentiviral vectors expressing small interfering RNA. Proc Natl Acad Sci U S A 100, 1844–1848. 10.1073/pnas.0437912100.

196. The Jackson Laboratory. Protocol 25394: Standard PCR Assay – Tg(Wnt1-cre). https://www.jax.org/Protocol?stockNumber=022501&protocolID=25394.

197. The Jackson Laboratory. Protocol 29436: Standard PCR Assay – Gt(ROSA)26Sor(tdTomato-WPRE). https://www.jax.org/Protocol?stockNumber=007914&protocolID=29436.

198. Beckman Coulter Life Sciences (2025). CytExpert Software.

199. ZEISS Microscopy (2025). ZEISS ZEN Microscopy Software.

200. Adobe Inc. (2025). Adobe Photoshop.

201. Rosin, J.M., Kurrasch, D.M., and Cobb, J. (2015). Shox2 is required for the proper development of the facial motor nucleus and the establishment of the facial nerves. BMC Neurosci 16, 39. 10.1186/s12868-015-0176-0.

202. Tosun, B., Wolff, L.I., Houben, A., Nutt, S., and Hartmann, C. (2022). Osteoclasts and Macrophages-Their Role in Bone Marrow Cavity Formation During Mouse Embryonic Development. J Bone Miner Res 37, 1761–1774. 10.1002/jbmr.4629.

203. Ralphs, J.R. (1992). Chondrogenesis and myogenesis in micromass cultures of mesenchyme from mouse facial primordia. In Vitro Cell Dev Biol 28A, 369–372. 10.1007/BF02877061.

204. Ueharu, H., Yang, J., Komatsu, Y., and Mishina, Y. (2022). Isolation and Culture of Cranial Neural Crest Cells from the First Branchial Arch of Mice. Bio Protoc 12, e4371. 10.21769/BioProtoc.4371.

205. Tophkhane, S.S., Fu, K., Verheyen, E.M., and Richman, J.M. (2024). Craniofacial studies in chicken embryos confirm the pathogenicity of human FZD2 variants associated with Robinow syndrome. Dis Model Mech 17. 10.1242/dmm.050584.

206. Rosin, J.M., Sinha, S., Biernaskie, J., and Kurrasch, D.M. (2021). A subpopulation of embryonic microglia respond to maternal stress and influence nearby neural progenitors. Dev Cell 56, 1326–1345 e1326. 10.1016/j.devcel.2021.03.018.

207. Hao, Y., Stuart, T., Kowalski, M.H., Choudhary, S., Hoffman, P., Hartman, A., Srivastava, A., Molla, G., Madad, S., Fernandez-Granda, C., and Satija, R. (2024). Dictionary learning for integrative, multimodal and scalable single-cell analysis. Nat Biotechnol 42, 293–304. 10.1038/s41587-023-01767-y.

208. Hao, Y., Hao, S., Andersen-Nissen, E., Mauck, W.M., 3rd, Zheng, S., Butler, A., Lee, M.J., Wilk, A.J., Darby, C., Zager, M., et al. (2021). Integrated analysis of multimodal single-cell data. Cell 184, 3573–3587 e3529. 10.1016/j.cell.2021.04.048.

209. Stuart, T., Butler, A., Hoffman, P., Hafemeister, C., Papalexi, E., Mauck, W.M., 3rd, Hao, Y., Stoeckius, M., Smibert, P., and Satija, R. (2019). Comprehensive Integration of Single-Cell Data. Cell 177, 1888–1902 e1821. 10.1016/j.cell.2019.05.031.

210. Butler, A., Hoffman, P., Smibert, P., Papalexi, E., and Satija, R. (2018). Integrating single-cell transcriptomic data across different conditions, technologies, and species. Nat Biotechnol 36, 411–420. 10.1038/nbt.4096.

211. Satija, R., Farrell, J.A., Gennert, D., Schier, A.F., and Regev, A. (2015). Spatial reconstruction of single-cell gene expression data. Nat Biotechnol 33, 495–502. 10.1038/nbt.3192.

212. Posit team (2025). RStudio: Integrated Development Environment for R (Posit Software, PBC).

213. R Core Team (2024). R: A Language and Environment for Statistical Computing (R Foundation for Statistical Computing).

214. Hagiwara, K., Obayashi, T., Sakayori, N., Yamanishi, E., Hayashi, R., Osumi, N., Nakazawa, T., and Nishida, K. (2014). Molecular and cellular features of murine craniofacial and trunk neural crest cells as stem cell-like cells. PLoS One 9, e84072. 10.1371/journal.pone.0084072.

215. Schindelin, J., Arganda-Carreras, I., Frise, E., Kaynig, V., Longair, M., Pietzsch, T., Preibisch, S., Rueden, C., Saalfeld, S., Schmid, B., et al. (2012). Fiji: an open-source platform for biological-image analysis. Nat Methods 9, 676–682. 10.1038/nmeth.2019.

216. Schneider, C.A., Rasband, W.S., and Eliceiri, K.W. (2012). NIH Image to ImageJ: 25 years of image analysis. Nat Methods 9, 671–675. 10.1038/nmeth.2089.

217. Gutiérrez, M.L., Guevara, J., and Barrera, L.A. (2012). Semi-automatic grading system in histologic and immunohistochemistry analysis to evaluate in vitro chondrogenesis. Universitas Scientiarum 17, 167–178.

218. GraphPad Software (2025). GraphPad Prism.

219. Kay, M., Elkin, L., Higgins, J.J., and Wobbrock, J.O. (2025). mjskay/ARTool: ARTool 0.11.2 (Zenodo).

220. Wobbrock, J.O., Findlater, L., Gergle, D., and Higgins, J.J. (2011). The aligned rank transform for nonparametric factorial analyses using only anova procedures. Proceedings of the SIGCHI Conference on Human Factors in Computing Systems. Association for Computing Machinery.

