## Supplemental Figures & Tables for "CSF1R+ macrophage and osteoclast depletion impairs neural crest proliferation and craniofacial morphogenesis"

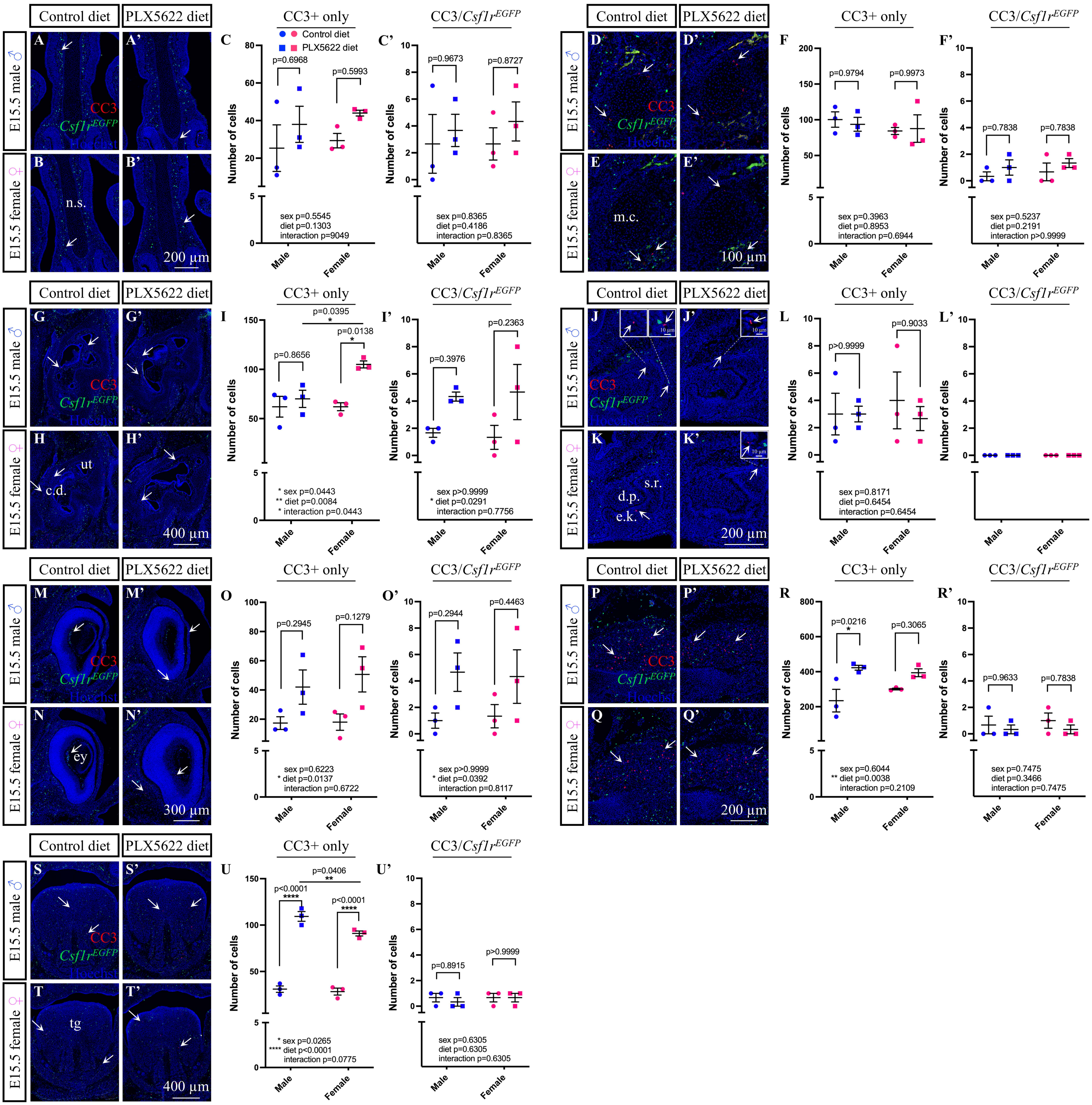
Control diet
PLX5622 diet
E15.5 male ♂
E15.5 female ♀
A
A'
B
B'
CC3
Csf1rEGFP
Hoechst
n.s.
200 µm
C
CC3+ only
Control diet
PLX5622 diet
p=0.6968
p=0.5993
sex p=0.5545
diet p=0.1303
interaction p=9049
C'
CC3/Csf1rEGFP
p=0.9673
p=0.8727
sex p=0.8365
diet p=0.4186
interaction p=0.8365
Number of cells
Male
Female
D
D'
E
E'
m.c.
100 µm
F
p=0.9794
p=0.9973
sex p=0.3963
diet p=0.8953
interaction p=0.6944
F'
p=0.7838
p=0.7838
sex p=0.5237
diet p=0.2191
interaction p>0.9999
G
G'
H
H'
ut
c.d.
400 µm
I
p=0.0395
p=0.0138
p=0.8656
* sex p=0.0443
** diet p=0.0084
* interaction p=0.0443
I'
p=0.2363
p=0.3976
sex p>0.9999
* diet p=0.0291
interaction p=0.7756
J
J'
K
K'
10 µm
s.r.
d.p.
e.k.
200 µm
L
p>0.9999
p=0.9033
sex p=0.8171
diet p=0.6454
interaction p=0.6454
L'
M
M'
N
N'
ey
300 µm
O
p=0.2945
p=0.1279
sex p=0.6223
* diet p=0.0137
interaction p=0.6722
O'
p=0.2944
p=0.4463
sex p>0.9999
* diet p=0.0392
interaction p=0.8117
P
P'
Q
Q'
200 µm
R
p=0.0216
p=0.3065
sex p=0.6044
** diet p=0.0038
interaction p=0.2109
R'
p=0.9633
p=0.7838
sex p=0.7475
diet p=0.3466
interaction p=0.7475
S
S'
T
T'
tg
400 µm
U
p=0.0406
p<0.0001
p<0.0001
* sex p=0.0265
**** diet p<0.0001
interaction p=0.0775
U'
p=0.8915
p>0.9999
sex p=0.6305
diet p=0.6305
interaction p=0.6305

**Supplementary Figure 1. Exposure to PLX5622 During Gestation Leads to Tissue-Specific Increases in Apoptosis.** (A-U') Immunofluorescent staining and quantification of active cleaved Caspase 3 (CC3) single-positive (arrows) and CC3+/*Csf1r*$^{EGFP+}$ double-positive cells in and around the E15.5 nasal septum (n.s.; A-C'), Meckel's cartilage (m.c.; D-F'), ear (G-I'), maxillary incisor (J-L'), eye (ey; M-O'), trigeminal (P-R'), and tongue (tg; S-U'). N=3 embryos per sex/treatment from 2-3 dams. Abbreviations: c.d., cochlear duct; d.p., dental papilla; e.k., enamel knot; s.r., stellate reticulum; ut, utricle. Counts represent mean ± SEM and were analyzed by a two-way ANOVA with Tukey's post-hoc test.

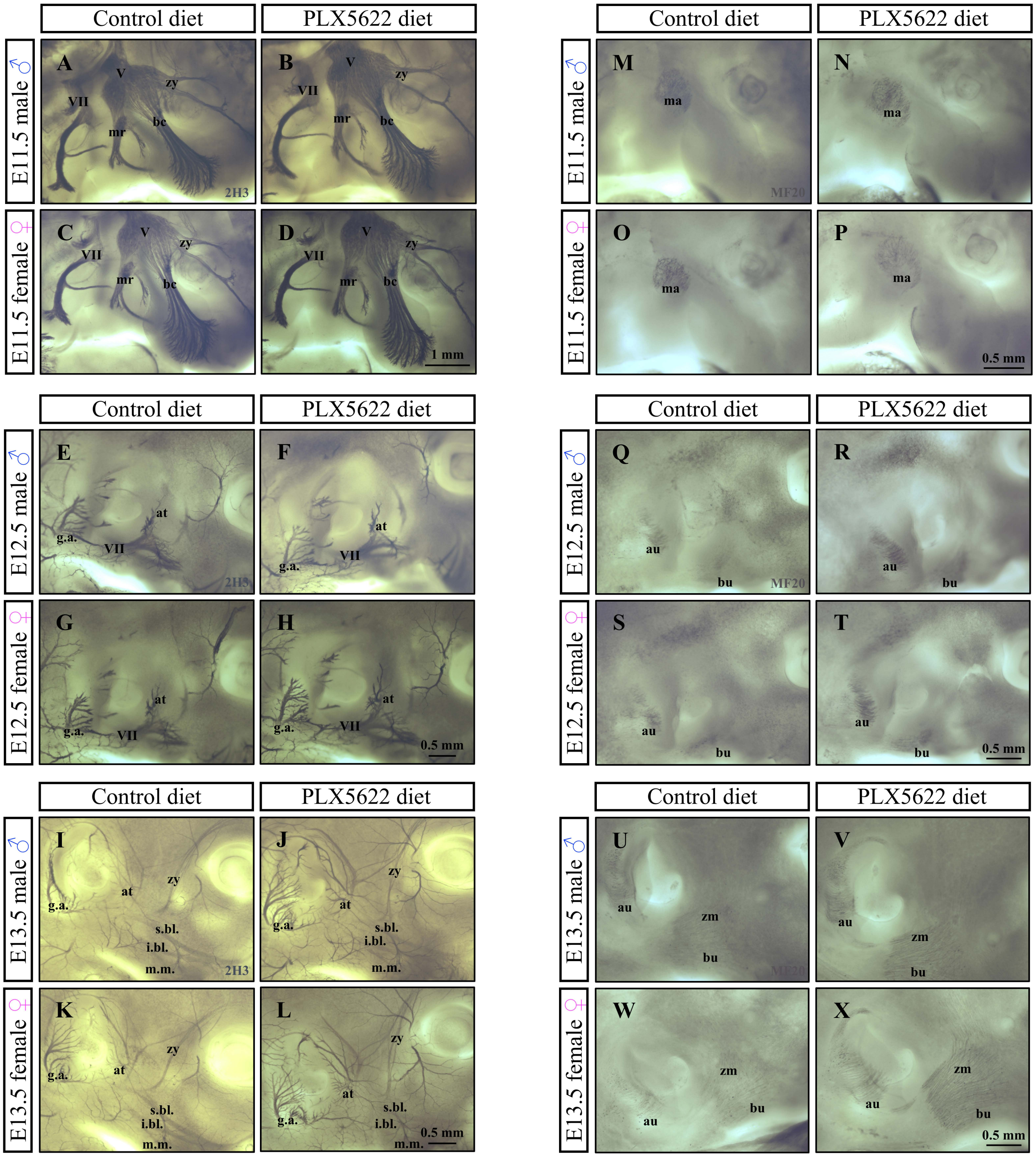
Control diet
PLX5622 diet
E11.5 male ♂
E11.5 female ♀
E12.5 male ♂
E12.5 female ♀
E13.5 male ♂
E13.5 female ♀
A B C D E F G H I J K L M N O P Q R S T U V W X
V
VII
zy
mr
bc
2H3
1 mm
ma
MF20
0.5 mm
at
g.a.
au
bu
s.bl.
i.bl.
m.m.
zm

**Supplementary Figure 2. Prenatal Exposure to PLX5622 Does Not Impact Nerve or Muscle Development.** (A-L) Whole-mount 2H3 immunostaining of E11.5 (A-D), E12.5 (E-H), and E13.5 (I-L) CD1 embryos. N=3-7 embryos per sex/treatment/time-point from 2-3 dams. (M-X) Whole-mount MF20 immunostaining of E11.5 (M-P), E12.5 (Q-T), and E13.5 (U-X) CD1 embryos. N=3-4 embryos per sex/treatment/time-point from 2-3 dams. Abbreviations: V, trigeminal nerve; VII, facial nerve; at, auriculotemporal nerve; au, auricularis; bc, buccal branch of the trigeminal; bu, buccinator; g.a., great auricular nerve; i.bl., inferior buccolabial nerve; ma, masseter; mr, marginal branch of the trigeminal; m.m., marginal mandibular nerve; s.bl., superior buccolabial nerve; zm, zygomaticomandibularis; zy, zygomatic branch of the trigeminal.

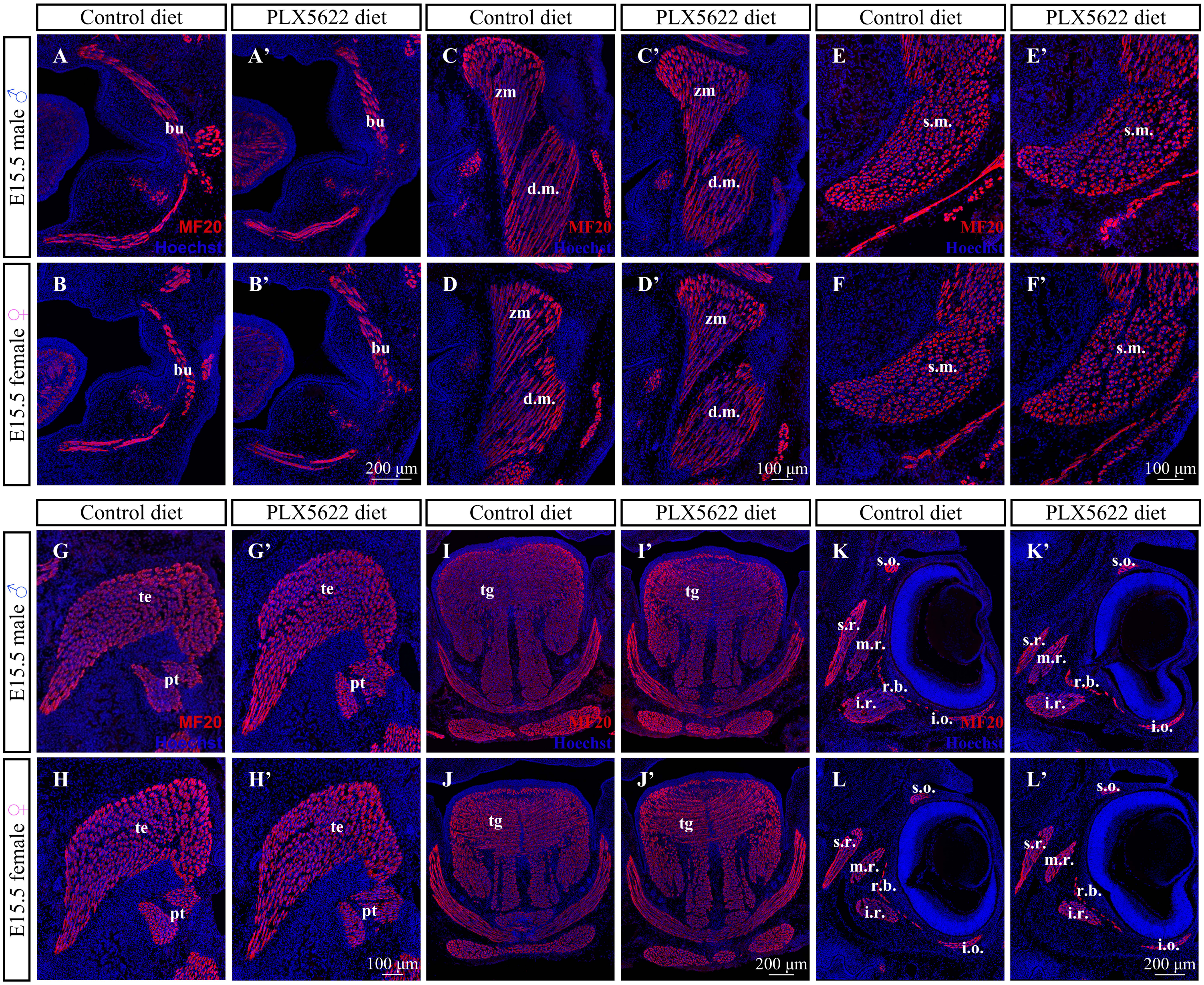
Control diet
PLX5622 diet
Control diet
PLX5622 diet
Control diet
PLX5622 diet
E15.5 male ♂
E15.5 female ♀
A
A'
B
B'
bu
MF20
Hoechst
200 µm
C
C'
D
D'
zm
d.m.
100 µm
E
E'
F
F'
s.m.
100 µm
G
G'
H
H'
te
pt
100 µm
I
I'
J
J'
tg
200 µm
K
K'
L
L'
s.o.
s.r.
m.r.
r.b.
i.r.
i.o.
200 µm

**Supplementary Figure 3. Gestational Exposure to PLX5622 Does Not Disrupt Muscle Fiber Structure**. (A-L') Immunofluorescent staining with MF20 in E15.5 C57BL/6 craniofacial tissue cryosections suggest that muscle fiber diameter and cellularity are comparable between control and PLX5622 embryos in the buccinator (bu; A-B'), zygomaticomandibularis (zm; C-D'), masseter (C-F'), temporalis (te) and pterygoid (pt; G-H'), tongue (tg; I-J'), and extraocular muscles (K-L'). N=3 embryos per sex/treatment from 2-3 dams. Abbreviations: d.m., deep masseter; i.o., inferior oblique; i.r., inferior rectus; m.r., medial rectus; r.b., retractor bulbi; s.m., superficial masseter; s.o., superior oblique; s.r., superior rectus.

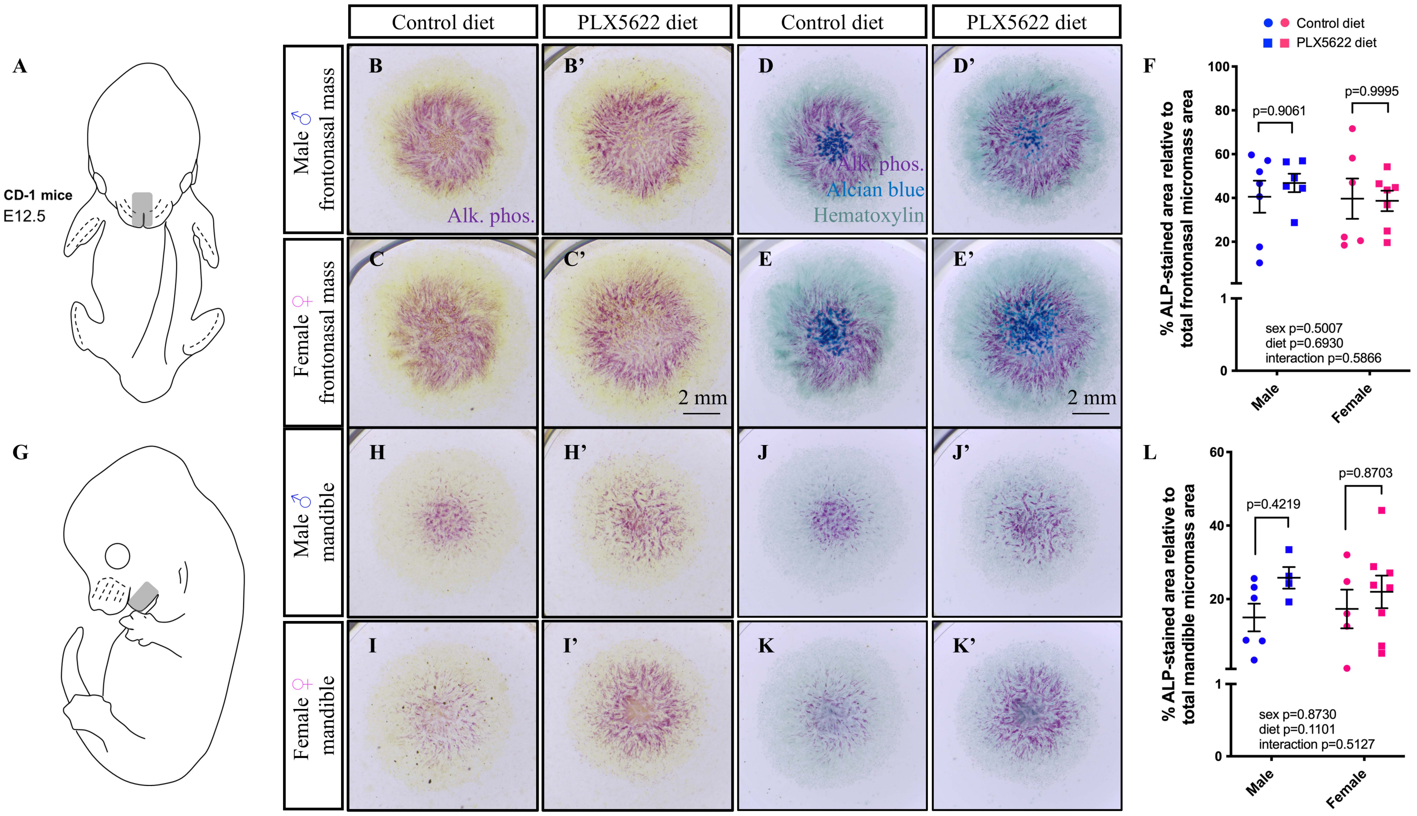

Control diet
PLX5622 diet
Control diet
PLX5622 diet
A
CD-1 mice
E12.5
B
B'
D
D'
Male ♂ frontonasal mass
Alk. phos.
Alk. phos.
Alcian blue
Hematoxylin
C
C'
E
E'
Female ♀ frontonasal mass
2 mm
2 mm
F
Control diet
PLX5622 diet
% ALP-stained area relative to total frontonasal micromass area
p=0.9061
p=0.9995
sex p=0.5007
diet p=0.6930
interaction p=0.5866
Male
Female
G
H
H'
J
J'
Male ♂ mandible
I
I'
K
K'
Female ♀ mandible
L
% ALP-stained area relative to total mandible micromass area
p=0.4219
p=0.8703
sex p=0.8730
diet p=0.1101
interaction p=0.5127
Male
Female

**Supplementary Figure 4. Assessing Osteogenesis in Facial Micromass Cultures Collected from Embryos Exposed to PLX5622 During Gestation.** (A and G) Schematic illustrating micro-dissected area for collecting mesenchymal cells from the E12.5 CD1 frontonasal mass (A) and mandible (G) for micromass culture. (B-F, H-L) Alkaline phosphatase (Alk. phos. or ALP), Alcian blue, and hematoxylin staining and quantification of E12.5 micromass cultures collected from the frontonasal mass (B-F) and mandible (H-L) of control and PLX5622 embryos. N=4-8 cultures per sex/treatment from 3 dams. Measurements represent mean ± SEM and were analyzed by a two-way ANOVA with Tukey's post-hoc test.

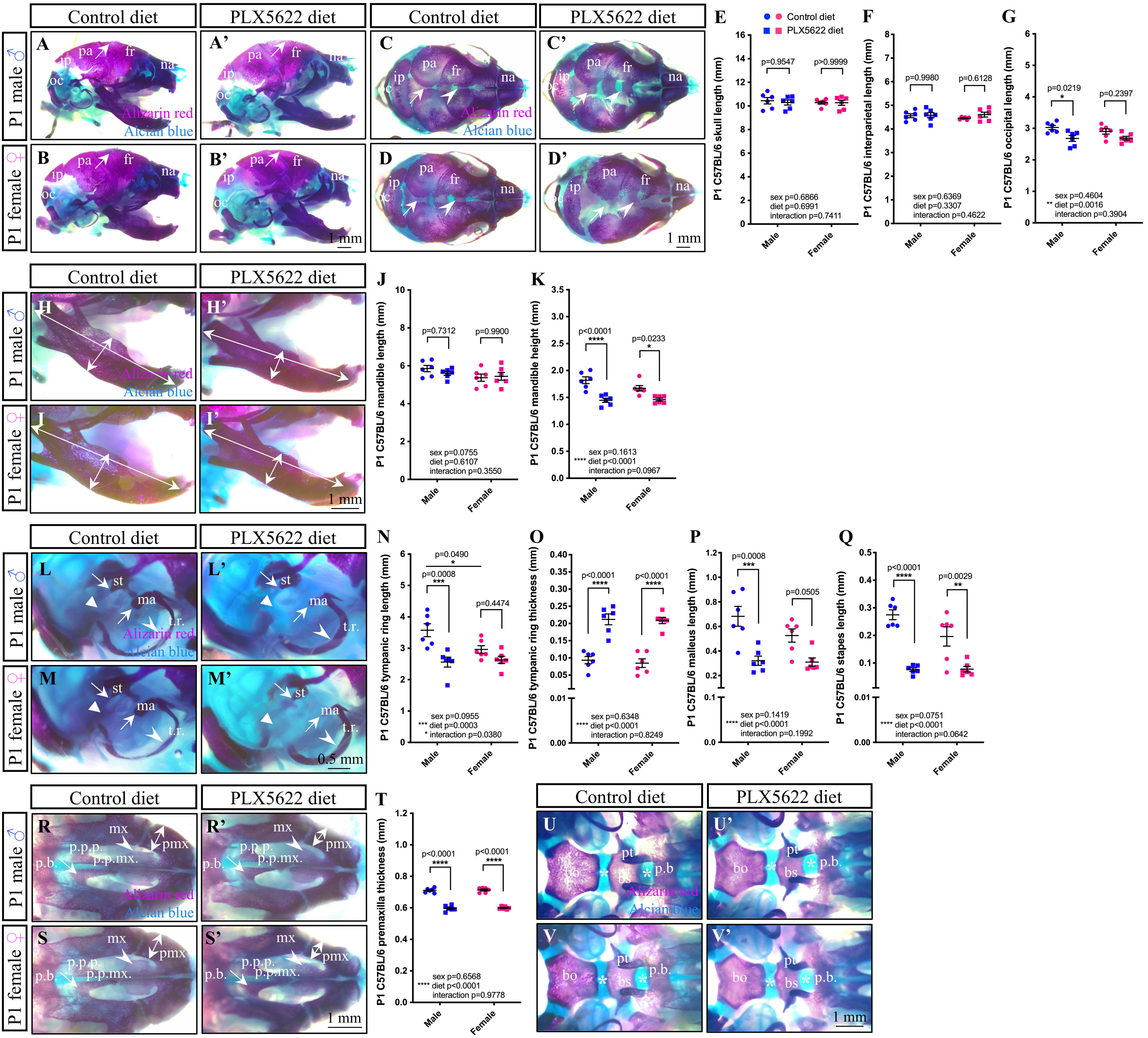

Control diet
PLX5622 diet
P1 male ♂
P1 female ♀
Alizarin red
Alcian blue
A
A'
B
B'
C
C'
D
D'
pa
fr
na
ip
oc
1 mm
E
F
G
Control diet
PLX5622 diet
P1 C57BL/6 skull length (mm)
p=0.9547
p>0.9999
sex p=0.6866
diet p=0.6991
interaction p=0.7411
P1 C57BL/6 interparietal length (mm)
p=0.9980
p=0.6128
sex p=0.6369
diet p=0.3307
interaction p=0.4622
P1 C57BL/6 occipital length (mm)
p=0.0219
p=0.2397
sex p=0.4604
** diet p=0.0016
interaction p=0.3904
Male
Female
H
H'
I
I'
J
K
P1 C57BL/6 mandible length (mm)
p=0.7312
p=0.9900
sex p=0.0755
diet p=0.6107
interaction p=0.3550
P1 C57BL/6 mandible height (mm)
p<0.0001
p=0.0233
sex p=0.1613
**** diet p<0.0001
interaction p=0.0967
L
L'
M
M'
st
ma
t.r.
0.5 mm
N
O
P
Q
P1 C57BL/6 tympanic ring length (mm)
p=0.0490
p=0.0008
p=0.4474
sex p=0.0955
*** diet p=0.0003
* interaction p=0.0380
P1 C57BL/6 tympanic ring thickness (mm)
p<0.0001
p<0.0001
sex p=0.6348
**** diet p<0.0001
interaction p=0.8249
P1 C57BL/6 malleus length (mm)
p=0.0008
p=0.0505
sex p=0.1419
**** diet p<0.0001
interaction p=0.1992
P1 C57BL/6 stapes length (mm)
p<0.0001
p=0.0029
sex p=0.0751
**** diet p<0.0001
interaction p=0.0642
R
R'
S
S'
mx
pmx
p.p.p.
p.p.mx.
p.b.
T
P1 C57BL/6 premaxilla thickness (mm)
p<0.0001
p<0.0001
sex p=0.6568
**** diet p<0.0001
interaction p=0.9778
U
U'
V
V'
bo
bs
pt
p.b.

**Supplementary Figure 5. Strain Differences in PLX5622-Driven Craniofacial Bone Morphogenesis Phenotypes.** (A-B') Lateral view of P1 C57BL/6 skulls highlights doming of the cranial vault in PLX5622 pups compared to controls (arrows). (C-D') Dorsal view of P1 C57BL/6 skulls shows impaired sagittal sutures (arrows) and impaired interfrontal sutures (arrowheads) in PLX5622 pups compared to controls. (E-G) Quantification of skull (E), interparietal (F), and occipital (G) bone lengths in P1 control and PLX5622 C57BL/6 pups. (H-I') Lateral view of P1 C57BL/6 mandibles. (J-K) Quantification of mandible length (J) and height (K). Mandible length (horizontal arrows) is consistent between control and PLX5622 pups, while mandible height (vertical arrows) is disrupted in both sexes. (L-Q) Lateral view of the P1 C57BL/6 ear (L-M') and quantification of tympanic ring (t.r.) length (N) and thickness (O) and malleus (P) and stapes (Q) lengths. Arrows mark disrupted ossicle development (incus absent in C57BL/6 pups regardless of diet) of the malleus (ma), and stapes (st). Triangles mark disruptions to the otic capsule. Arrowheads mark disruptions to the tympanic ring. (R-S) Ventral view of the P1 C57BL/6 palate highlight disrupted hard palate morphogenesis (arrowheads). (T) Quantification of premaxilla (pm) thickness. (U-V') Ventral view of the P1 C57BL/6 cranial base shows widening of the spheno-occipital and intersphenoid synchondroses (asterisks) in PLX5622 pups compared to controls. N=6 pups per sex/treatment from 3-4 dams. Abbreviations: bo, basioccipital; bs, basisphenoid; fr, frontal bone; ip, interparietal bone; mx, maxilla; na, nasal bone; oc, occipital bone; pa, parietal bone; p.b., palatine bone; p.p.mx., palatine process of the maxilla; p.p.p., palatine process of the palatine; pt, pterygoid process. Measurements represent mean ± SEM and were analyzed by a two-way ANOVA with Tukey's post hoc test.

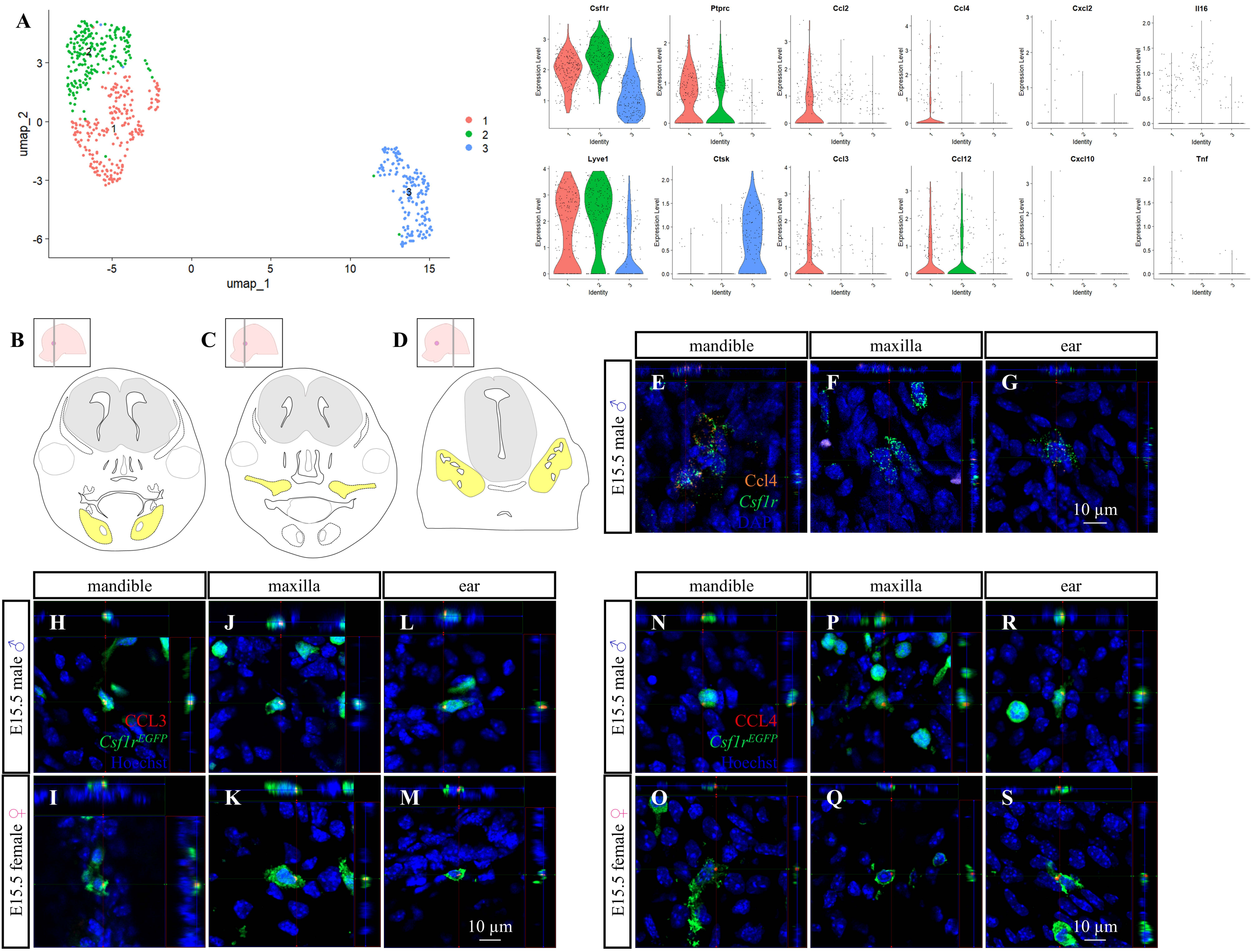
A
umap_1
umap_2
1
2
3
Csf1r
Ptprc
Ccl2
Ccl4
Cxcl2
Il16
Lyve1
Ctsk
Ccl3
Ccl12
Cxcl10
Tnf
Expression Level
Identity
B
C
D
mandible
maxilla
ear
E15.5 male ♂
E15.5 female ♀
E
F
G
Ccl4
Csf1r
DAPI
10 μm
H
I
J
K
L
M
CCL3
Csf1r^EGFP
Hoechst
N
O
P
Q
R
S
CCL4

**Supplementary Figure 6. Cytokines are expressed by *Csf1r*/CSF1R+ cells in Craniofacial Tissues**. (A) UMAP clustering of *Csf1r*-expressing cells from published E12.5 and E13.5 FVB/NJ and C57BL/6J craniofacial mesenchyme single-cell RNA sequencing datasets[1,2]. Violin plots show expression of a number of macrophage and/or osteoclast markers across the three clusters. Cluster 1 highly expresses *Ccl2, Ccl3, Ccl4,* and *Ccl12*. *Ccl12* is also highly expressed in cluster 2. Cluster 3 expresses osteoclast marker *Ctsk*. (B-D) Schematics illustrating which areas of coronal sections of the mandible (B), maxilla (C), and ear (D) were utilized for imaging in panels E-S. (E-G) Orthographic projections of cells expressing both *Csf1r* and *Ccl4* in the E15.5 CD1 mandible (E), maxilla (F), and ear (G). (H-S) Orthographic projections of $Csf1r^{EGFP+}$ cells that co-express CCL3 (H-M) or CCL4 (N-S) in the E15.5 mandible (H, I, N, O), maxilla (J, K, P, Q), and ear (L, M, R, S).

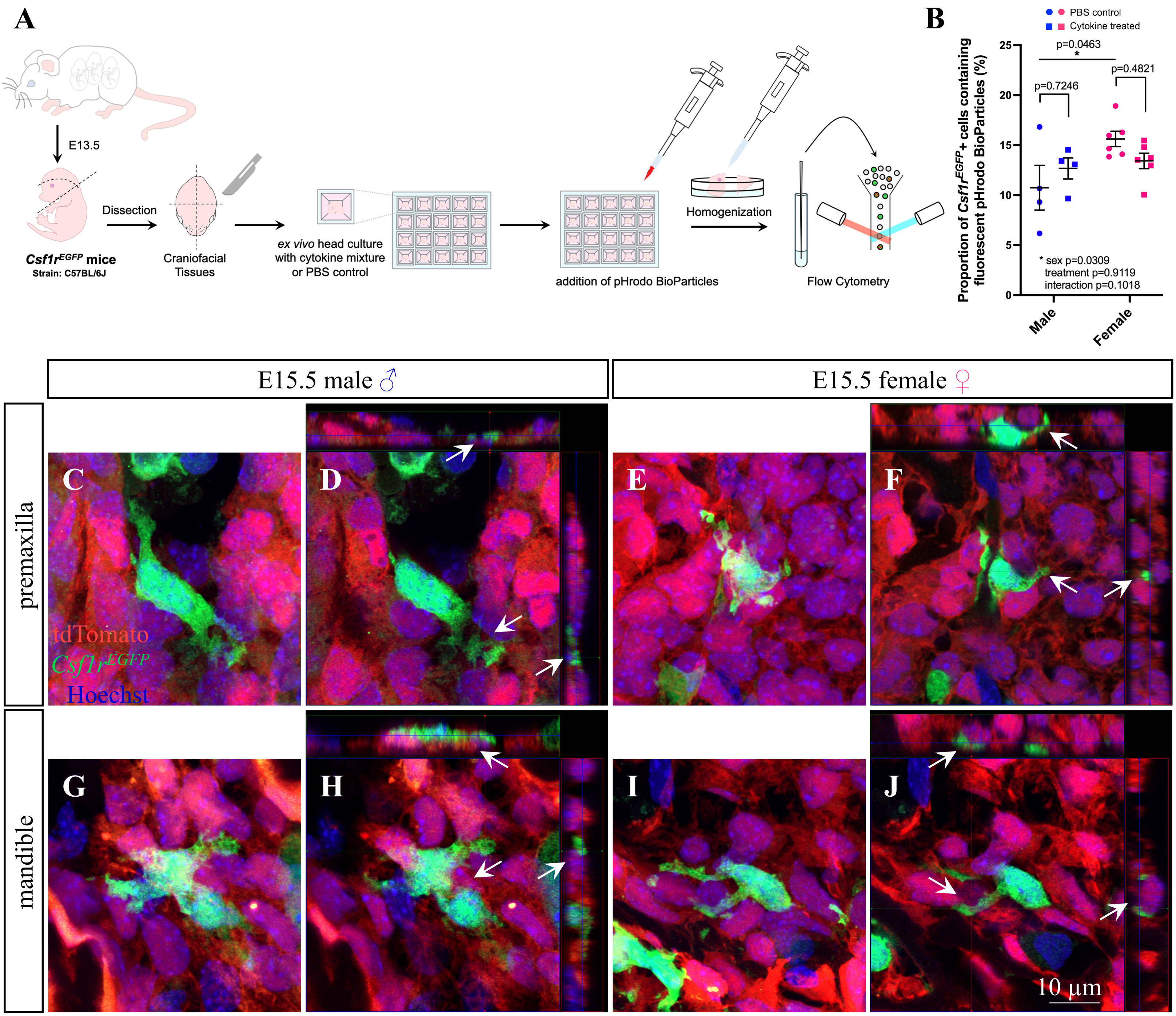
A
E13.5
Dissection
Csf1r^EGFP mice
Strain: C57BL/6J
Craniofacial Tissues
ex vivo head culture with cytokine mixture or PBS control
addition of pHrodo BioParticles
Homogenization
Flow Cytometry
B
PBS control
Cytokine treated
Proportion of Csf1r^EGFP+ cells containing fluorescent pHrodo BioParticles (%)
p=0.0463
p=0.7246
p=0.4821
* sex p=0.0309
treatment p=0.9119
interaction p=0.1018
Male
Female
E15.5 male ♂
E15.5 female ♀
premaxilla
mandible
C
D
E
F
G
H
I
J
tdTomato
Csf1r^EGFP
Hoechst
10 μm

**Supplementary Figure 7. Phagocytic Activity of CSF1R+ Cells is not Altered in Response to Cytokine Exposure but CSF1R+ Cells Interact with Neural Crest Cells in Craniofacial Tissues.** (A) Schematic illustrating E13.5 *Csf1r*$^{EGFP}$ craniofacial tissue collection, PBS or cytokine treatment, pHrodo *E. coli* BioParticle incubation, and flow cytometry for EGFP+ cell phagocytosis assay. (B) Quantification of the proportion of *Csf1r*$^{EGFP+}$ cells containing fluorescent pHrodo BioParticles using flow cytometry (N=4-6 embryos per sex/treatment from 3 dams). Counts represent mean ± SEM and were analyzed by a two-way ANOVA with Tukey's post-hoc test. (C-J) Immunofluorescent images (C, E, G, I) and orthographic projections (D, F, H, J) of *Csf1r*$^{EGFP+}$ cells interacting with *Wnt1*$^{Cre}$-driven tdTomato+ cells (arrows) in the E15.5 male (C, D, G, H) and female (E, F, I, J) premaxilla (C-F) and mandible (G-J).

**Supplementary Table 1.** Exposure to PLX5622 During Gestation Does Not Affect Litter Size, Number of Embryo Resorptions or Embryo Sex Ratio.

| | | Number of embryos per litter | Number of embryo resorptions per litter (Number of litters with resorptions) | Sex ratio of male to female embryos per litter | Number of litters analyzed |
|---|---|---|---|---|---|
| **CD1** | **Control diet** | 13.43 ± 0.69 | 0.43 ± 0.20 (3/7) | 1.14 ± 0.25 | 7 |
| | **PLX5622 diet** | 11.71 ± 0.52 | 0.29 ± 0.18 (2/7) | 1.47 ± 0.51 | 7 |
| | **t-test** | p=0.0698 | p=0.6110 | p=0.5676 | – |
| **C57BL/6** | **Control diet** | 6.85 ± 0.56 | 0.62 ± 0.22 (7/26) | 1.28 ± 0.20 | 26 |
| | **PLX5622 diet** | 7.29 ± 0.49 | 0.91 ± 0.29 (10/23) | 1.53 ± 0.28 | 23 |
| | **t-test** | p=0.5683 | p=0.4176 | p=0.4612 | – |

**Supplementary Table 2.** Luminex Multiplex Analysis of Cytokines and Chemokines with Little or No Response to Gestational PLX5622 Exposure.

| Cytokine/ chemokine | Sex | Control diet concentration mean ± SEM (pg/mL) | PLX5622 diet concentration mean ± SEM (pg/mL) | Control diet mean ± SEM $log_2$[conc (pg/mL)] | PLX5622 diet mean ± SEM $log_2$[conc (pg/mL)] | Two-way ANOVA results (p values) ($log_2$ data used) (int = interaction) | Tukey's post-hoc test p-value (CON vs. PLX) |
|---|---|---|---|---|---|---|---|
| **CCL17** | Male | 0.69 ± 0.04 | 0.54 ± 0.01 | -0.560 ± 0.088 | -0.884 ± 0.036 | **int: 0.4171** **diet: ****<0.0001** | ***0.0146** |
| | Female | 0.78 ± 0.06 | 0.56 ± 0.02 | -0.390 ± 0.096 | -0.830 ± 0.043 | **sex: 0.1258** | *****0.0006** |
| **CCL19** | Male | 18.55 ± 1.37 | 21.74 ± 3.89 | 4.183 ± 0.103 | 4.308 ± 0.204 | **int: 0.0654** **diet: 0.3468** | **0.9062** |
| | Female | 21.34 ± 1.68 | 16.27 ± 0.75 | 4.384 ± 0.104 | 4.011 ± 0.070 | **sex: 0.7155** | **0.2003** |
| **CCL20** | Male | 0.81 ± 0.03 | 0.70 ± 0.02 | -0.318 ± 0.047 | -0.523 ± 0.046 | **int: 0.6653** **diet: ****<0.0001** | ***0.0270** |
| | Female | 0.82 ± 0.04 | 0.69 ± 0.02 | -0.295 ± 0.060 | -0.542 ± 0.039 | **sex: 0.9652** | ****0.0057** |
| **CCL21** | Male | 5542.67 ± 713.41 | 3811.79 ± 374.76 | 12.333 ± 0.197 | 11.831 ± 0.162 | **int: *0.0146** **diet: 0.7361** | **0.1859** |
| | Female | 3868.52 ± 431.87 | 5058.44 ± 573.44 | 11.846 ± 0.163 | 12.231 ± 0.164 | **sex: 0.8022** | **0.4008** |
| **CXCL9** | Male | 2.29 ± 0.77 | 1.11 ± 0.09 | 0.742 ± 0.377 | 0.107 ± 0.120 | **int: 0.3338** **diet: **0.0015** | **0.3070** |
| | Female | 2.11 ± 0.44 | 0.82 ± 0.06 | 0.819 ± 0.303 | -0.315 ± 0.100 | **sex: 0.5021** | ***0.0174** |
| **Eotaxin** | Male | 33.34 ± 3.83 | 32.13 ± 3.11 | 4.981 ± 0.169 | 4.948 ± 0.148 | **int: 0.6778** **diet: 0.5333** | **>0.9999** |
| | Female | 34.89 ± 6.01 | 28.58 ± 1.74 | 4.985 ± 0.213 | 4.817 ± 0.084 | **sex: 0.6949** | **0.9762** |
| **IGF-1** | Male | 1182.74 ± 123.16 | 1769.62 ± 91.52 | 10.151 ± 0.139 | 10.774 ± 0.075 | **int: 0.0633** **diet: 0.0588** | ***0.0460** |
| | Female | 1365.03 ± 243.37 | 1239.96 ± 104.38 | 10.233 ± 0.254 | 10.239 ± 0.115 | **sex: 0.1668** | **>0.9999** |
| **IL-2** | Male | 0.85 ± 0.04 | 0.70 ± 0.07 | -0.247 ± 0.073 | -0.579 ± 0.149 | **int: *0.0300** **diet: 0.4389** | **0.1660** |
| | Female | 0.74 ± 0.04 | 0.83 ± 0.06 | -0.458 ± 0.075 | -0.296 ± 0.111 | **sex: 0.7438** | **0.7101** |
| **IL-9** | Male | 17.39 ± 4.63 | 10.31 ± 2.58 | 3.781 ± 0.344 | 3.084 ± 0.300 | **int: *0.0473** **diet: 0.7448** | **0.3448** |
| | Female | 7.28 ± 1.63 | 10.26 ± 1.83 | 2.663 ± 0.248 | 3.169 ± 0.266 | **sex: 0.0858** | **0.6149** |
| **IL-16** | Male | 89.24 ± 21.64 | 66.98 ± 10.52 | 6.216 ± 0.287 | 5.946 ± 0.202 | **int: 0.3450** **diet: *0.0417** | **0.8433** |
| | Female | 92.99 ± 22.11 | 48.46 ± 3.61 | 6.282 ± 0.284 | 5.568 ± 0.105 | **sex: 0.5060** | **0.1511** |
| **MMP-9** | Male | 571.39 ± 76.38 | 463.28 ± 39.49 | 9.070 ± 0.172 | 8.810 ± 0.131 | **int: 0.7126** **diet: 0.1056** | **0.4843** |
| | Female | 506.99 ± 33.99 | 448.62 ± 25.86 | 8.956 ± 0.108 | 8.791 ± 0.081 | **sex: 0.6038** | **0.7972** |
| **OPN** | Male | 3169.85 ± 320.26 | 3159.47 ± 191.52 | 11.579 ± 0.131 | 11.604 ± 0.088 | **int: 0.1392** **diet: 0.2078** | **0.9984** |
| | Female | 3218.71 ± 318.29 | 2538.11 ± 102.01 | 11.601 ± 0.133 | 11.300 ± 0.059 | **sex: 0.1967** | **0.2161** |
| **TIMP-1** | Male | 1202.48 ± 161.46 | 1300.59 ± 96.37 | 10.141 ± 0.175 | 10.311 ± 0.114 | **int: 0.0560** **diet: 0.4242** | **0.8398** |
| | Female | 1393.82 ± 153.51 | 1035.84 ± 94.67 | 10.376 ± 0.157 | 9.973 ± 0.123 | **sex: 0.7222** | **0.2186** |
| **VEGF** | Male | 4.86 ± 0.92 | 2.67 ± 0.82 | 2.031 ± 0.324 | 1.004 ± 0.361 | **int: *0.0196** **diet: 0.3502** | **0.0957** |
| | Female | 3.58 ± 0.61 | 4.32 ± 0.47 | 1.596 ± 0.329 | 2.051 ± 0.141 | **sex: 0.3170** | **0.7112** |

**CSF-1, CXCL5, EPO, GM-CSF, IFN-γ, IL-1α, IL-1β, IL-3, IL-4, IL-5, IL-7, IL-10, IL-11, IL-12p40, IL-12p70, IL-13, IL-15, IL-17, IL-20, and LIF content were at or below the level of detection.**

**Supplementary Table 3.** Comparison of Phenotypes Between PLX5622 Mice and *Csf1/Csf1r* Mutant Rodent Models.

| **PLX5622 mice (gestational exposure)** | ***Csf1***$^{op/op}$ **mice** | ***Csf1r* KO mice** | ***Csf1***$^{tl/tl}$ **rat** | ***Csf1r* KO rat** | ***Csf1r***$^{\Delta FIRE/\Delta FIRE}$ **mouse** |
|---|---|---|---|---|---|
| **CD1** (this study)[3,4]<br><br>**C57BL/6J:C57BL/6N** (this study) | **C3FeB6F1/J** *A*/*A*$^{w-J}$ [5-14]<br><br>**C3FeB6F1/J** *A*/*A*$^{w-J}$ **x CD1**[15,16]<br><br>**C57BL/6J**[12]<br><br>**C57BL/6N**[17]<br><br>**FVB/NJ**[16,18-20] | **C3FeB6F1/J** *A*/*A*$^{w-J}$ **x CD1**[15]<br><br>**C57BL/6**[21,22]<br><br>**C57BL/6N**[17]<br><br>**FVB/NJ**[19,20,23,24]<br><br>**FVB.129X1**[25] | **Fischer**[26-35]<br><br>**Osborne-Mendel**[36]<br><br>**Lewis**[37] | **Dark Agouti**[38-42]<br><br>**Sprague Dawley**[39]<br><br>**DA:SD**[43-45] | **C57BL/6J**[46]<br><br>**C57BL/6J:CBA**[46-51] |
| Significant depletion of macrophages throughout craniofacial tissues (this study)[3]:<br><br>21-63% depletion in E11.5-17.5 whole head (this study)<br><br>39-73% depletion around E15.5 nasal septum (this study)<br><br>61-78% depletion around E15.5 Meckel's cartilage (this study)<br><br>65-80% depletion in E15.5 ear (this study)<br><br>57-79% depletion in E15.5 eye (this study)<br><br>41-59% depletion in | Significant reduction in most tissue macrophages (26-100%)[5,6,15,16]<br><br>50-65% reduction in ear[7]<br><br>reduction in tooth pulp, gingiva, alveolar bone, tongue[52] | Significant reduction in tissue macrophages (32-93%)[15]<br><br>~50-100% reduction in cochlea[21,22] | Significant reduction in macrophages in peritoneal and pleural lavages (>98%)[31] | Significant reduction in tissue macrophages (~30-100%)[38-40,42,45] | Normal numbers of macrophages in most tissues[47-49]<br><br>~50-100% depletion of Langerhans cells and cardiac, kidney, and large peritoneal macrophages[48]<br><br>absence of embryonic macrophages up to E13[48] |

| | | | | | |
|---|---|---|---|---|---|
| E15.5 trigeminal (this study)<br><br>48-73% depletion in E15.5 tongue (this study) | | | | | |
| ~99% microglial depletion in hypothalamus by E15.5[4] | 0-64% reduction in microglia[8-10,20]<br><br>24-30% microglial reduction in hypothalamus[53]<br><br>No microglial reduction in adulthood in C57BL/6N[17] | ~99% reduction in microglia in hypothalamus, ~94% reduction in overall brain[17,20,25]<br><br>~85% reduction in microglia by E12.5[20] | Not shown | Absence of microglia[39,41,45] | Absence of microglia[47-51] |
| Absence of embryonic osteoclasts and TRAP activity at E15.5 (this study) | Reduction of osteoclasts and TRAP activity[11,12,15,16,18,29,52]<br><br>Remaining osteoclasts are small and abnormal[12,29] | Further reduction of osteoclasts and TRAP activity compared to *Csf1*$^{op/op}$ mice[15]<br><br>Absence of normal osteoclasts[23] | Absence (≥99%) of osteoclasts and TRAP activity[26-32,35] | Absence of osteoclasts and TRAP activity[38,45] | Normal numbers of osteoclasts and TRAP activity[48] |
| Decreased cranio-skeletal size[3]; decreased skull length (this study) | Decreased skull size and length[6,12,15,16,18] | Decreased skull size and length[15,25] | Decreased head size and length (not reported in study but observed in figures)[33,36] | Decreased skull size and length (not reported in study but observed in figures)[38,40,44,45] | No postnatal growth retardation[48] |
| Domed skull (this study)[3,4] | Domed skull[5-7,12,15,16,18] | Domed skull[15,25] | Not shown | Domed skull[41,45] | Not shown |
| Decreased calvarial bone size and density (interparietal and occipital significant only in CD1; parietal, frontal, and nasal observed) (this study) | Deformities in flat bony plates[15,16] | Deformities in flat bony plates[15] | Not shown | Decreased calvarial bone size and density[38,40,41,43] | Not shown |
| Impaired cranial sutures (coronal, sagittal, and interfrontal) (this study) | Not shown | Not shown | Not shown | Impaired cranial sutures[38,40,43] (coronal, sagittal, lambdoid, and interfrontal) | Not shown |

| | | | | | |
|---|---|---|---|---|---|
| Decreased mandible size[3]; increased mandible bone density at E15.5, decreased mandible height at P1, decreased mandible length in P1 CD1 females only (this study) | Increased mandible bone density[15]<br><br>Decreased mandible size and length, possibly decreased mandible height (not reported in study but observed in figures)[6,15,16,18] | Increased mandible bone density[15]<br><br>Decreased mandible size and length, possibly decreased mandible height (not reported in study but observed in figures)[15] | Not shown | Decreased mandible size and length (not reported in study but observed in figures)[44,45] | Not shown |
| Significantly smaller malleus and stapes, absence of incus (malleus not significant in C57BL/6 females; incus absent in all C57BL/6) (this study)<br><br>Shorter and thicker tympanic ring (not shorter in C57BL/6 females) (this study)<br><br>Decreased otic capsule bone density (this study) | Hypertrophy of malleus, incus, and stapes[7,13],<br><br>Increased otic capsule bone density[7] (not reported in study but observed in figures)[16]<br><br>Impaired hearing[7,14] | Impaired hearing[24] | Smaller and thicker stapes, immature bone in all auditory ossicles[34]<br><br>Majority hearing impaired[34] | Shorter tympanic ring (not reported in study but observed in figures)[44]<br><br>Increased otic capsule bone density (not reported in study but observed in figures)[44]<br><br>No evidence of sensory defects[44,45] | Not shown |
| Increased premaxilla bone density at E15.5 (this study)<br><br>Decreased premaxilla thickness at P1 (this study) | Increased premaxilla bone density (not reported in study but observed in figures)[15] | Increased premaxilla bone density (not reported in study but observed in figures)[15] | Not shown | Not shown | Not shown |
| Decreased palatine process size causing impaired palatine sutures (variable), no overt clefting (this study) | Not shown | Not shown | Not shown | Not shown | Not shown |

| | | | | | |
|---|---|---|---|---|---|
| Decreased cranial base bone size (only presphenoid significant) (this study)<br><br>Widened synchondroses (only intersphenoid significant) (this study) | Increased cranial base bone density (not reported in study but observed in figures)[15] | Increased cranial base bone density (not reported in study but observed in figures)[15] | Not shown | Increased cranial base bone density[44] | Not shown |
| Ectopic incisor enamel, extra molar cusp[4] | Toothless[5-7,12,15,16,18,54]<br><br>Unerupted molars visible with normal crowns and malformed roots[11,52] | Toothless[15,25] | Toothless[26,31,36,37]<br><br>Unerupted molars visible[36] | Toothless[38,40,41]<br><br>Unerupted rudimentary teeth visible[45] | Teeth develop and erupt normally[48] |
| Enlarged lateral ventricles (this study)<br><br>Lesions at cortico-striato-amygdalar boundary (this study) | Normal lateral ventricles[19] | Enlarged lateral ventricles[17,19] | Not shown | Enlarged lateral ventricles[40,41,44,45] | Normal lateral ventricles[48]<br><br>Lesions at cortico-striato-amygdalar and cortico-septal boundaries embryonically[46] |

**Supplementary Table 4.** Sequences of Primers Used for Genotyping for Sex and Transgenes.

| REAGENT or RESOURCE | SOURCE | IDENTIFIER |
|---|---|---|
| Oligonucleotides | | |
| SX (*Sly* & *Xlr*) forward primer: GATGATTTGAGTGGAAATGTGAGGTA | McFarlane et al.[55] | N/A |
| SX (*Sly* & *Xlr*) reverse primer: CTTATGTTTATAGGCATGCACCATGTA | McFarlane et al.[55] | N/A |
| *GFP* forward primer: AAGTTCATCTGCACCACCG | Tiscornia et al.[56] | N/A |
| *GFP* reverse primer: TCCTTGAAGAAGATGGTGCG | Tiscornia et al.[56] | N/A |
| Transgene forward primer for $Wnt1^{Cre}$ genotyping: CAGCGCCGCAACTATAAGAG | The Jackson Laboratory[57] | N/A |
| Transgene reverse primer for $Wnt1^{Cre}$ genotyping: CATCGACCGGTAATGCAG | The Jackson Laboratory[57] | N/A |
| Internal positive control forward primer for $Wnt1^{Cre}$ genotyping: CAAATGTTGCTTGTCTGGTG | The Jackson Laboratory[57] | N/A |
| Internal positive control reverse primer for $Wnt1^{Cre}$ genotyping: GTCAGTCGAGTGCACAGTTT | The Jackson Laboratory[57] | N/A |
| WT forward primer for $Rosa26^{tdTomato}$ genotyping: AAGGGAGCTGCAGTGGAGTA | The Jackson Laboratory[58] | N/A |
| WT reverse primer for $Rosa26^{tdTomato}$ genotyping: CCGAAAATCTGTGGGAAGTC | The Jackson Laboratory[58] | N/A |
| tdTomato forward primer for $Rosa26^{tdTomato}$ genotyping: CTGTTCCTGTACGGCATGG | The Jackson Laboratory[58] | N/A |
| WPRE mutant reverse primer for $Rosa26^{tdTomato}$ genotyping: GGCATTAAAGCAGCGTATCC | The Jackson Laboratory[58] | N/A |

### SUPPLEMENTAL REFERENCES


1. Angelozzi, M., Pellegrino da Silva, R., Gonzalez, M.V., and Lefebvre, V. (2022). Single-cell atlas of craniogenesis uncovers SOXC-dependent, highly proliferative, and myofibroblast-like osteodermal progenitors. Cell Rep *40*, 111045. 10.1016/j.celrep.2022.111045.
2. Rajderkar, S.S., Paraiso, K., Amaral, M.L., Kosicki, M., Cook, L.E., Darbellay, F., Spurrell, C.H., Osterwalder, M., Zhu, Y., Wu, H., et al. (2024). Dynamic enhancer landscapes in human craniofacial development. Nat Commun *15*, 2030. 10.1038/s41467-024-46396-4.
3. Nagra, A., Katsube, M., Gao, W., Rosin, J.M., and Vora, S.R. (2023). Embryonic inhibition of colony-stimulating factor 1 receptor impacts craniofacial morphogenesis. Orthod Craniofac Res *26 Suppl 1*, 20-28. 10.1111/ocr.12671.
4. Rosin, J.M., Vora, S.R., and Kurrasch, D.M. (2018). Depletion of embryonic microglia using the CSF1R inhibitor PLX5622 has adverse sex-specific effects on mice, including accelerated weight gain, hyperactivity and anxiolytic-like behaviour. Brain Behav Immun *73*, 682-697. 10.1016/j.bbi.2018.07.023.
5. Cecchini, M.G., Dominguez, M.G., Mocci, S., Wetterwald, A., Felix, R., Fleisch, H., Chisholm, O., Hofstetter, W., Pollard, J.W., and Stanley, E.R. (1994). Role of colony stimulating factor-1 in the establishment and regulation of tissue macrophages during postnatal development of the mouse. Development *120*, 1357-1372. 10.1242/dev.120.6.1357.
6. Ryan, G.R., Dai, X.M., Dominguez, M.G., Tong, W., Chuan, F., Chisholm, O., Russell, R.G., Pollard, J.W., and Stanley, E.R. (2001). Rescue of the colony-stimulating factor 1 (CSF-1)-nullizygous mouse (Csf1(op)/Csf1(op)) phenotype with a CSF-1 transgene and identification of sites of local CSF-1 synthesis. Blood *98*, 74-84. 10.1182/blood.v98.1.74.
7. Okano, T., and Kishimoto, I. (2019). Csf1 Signaling Regulates Maintenance of Resident Macrophages and Bone Formation in the Mouse Cochlea. Frontiers in Neurology *10*. 10.3389/fneur.2019.01244.
8. Kondo, Y., and Duncan, I.D. (2009). Selective reduction in microglia density and function in the white matter of colony-stimulating factor-1-deficient mice. J Neurosci Res *87*, 2686-2695. 10.1002/jnr.22096.
9. Sasaki, A., Yokoo, H., Naito, M., Kaizu, C., Shultz, L.D., and Nakazato, Y. (2000). Effects of macrophage-colony-stimulating factor deficiency on the maturation of microglia and brain macrophages and on their expression of scavenger receptor. Neuropathology *20*, 134-142. 10.1046/j.1440-1789.2000.00286.x.
10. Wegiel, J., Wisniewski, H.M., Dziewiatkowski, J., Tarnawski, M., Kozielski, R., Trenkner, E., and Wiktor-Jedrzejczak, W. (1998). Reduced number and altered morphology of microglial cells in colony stimulating factor-1-deficient osteopetrotic op/op mice. Brain Res *804*, 135-139. 10.1016/s0006-8993(98)00618-0.
11. Kodama, H., Yamasaki, A., Nose, M., Niida, S., Ohgame, Y., Abe, M., Kumegawa, M., and Suda, T. (1991). Congenital osteoclast deficiency in osteopetrotic (op/op) mice is cured by injections of macrophage colony-stimulating factor. J Exp Med *173*, 269-272. 10.1084/jem.173.1.269.
12. Marks, S.C., Jr., and Lane, P.W. (1976). Osteopetrosis, a new recessive skeletal mutation on chromosome 12 of the mouse. J Hered *67*, 11-18. 10.1093/oxfordjournals.jhered.a108657.

13. Kanzaki, S., Takada, Y., Niida, S., Takeda, Y., Udagawa, N., Ogawa, K., Nango, N., Momose, A., and Matsuo, K. (2011). Impaired vibration of auditory ossicles in osteopetrotic mice. Am J Pathol *178*, 1270-1278. 10.1016/j.ajpath.2010.11.063.
14. Michaelson, M.D., Bieri, P.L., Mehler, M.F., Xu, H., Arezzo, J.C., Pollard, J.W., and Kessler, J.A. (1996). CSF-1 deficiency in mice results in abnormal brain development. Development *122*, 2661-2672. 10.1242/dev.122.9.2661.
15. Dai, X.M., Ryan, G.R., Hapel, A.J., Dominguez, M.G., Russell, R.G., Kapp, S., Sylvestre, V., and Stanley, E.R. (2002). Targeted disruption of the mouse colony-stimulating factor 1 receptor gene results in osteopetrosis, mononuclear phagocyte deficiency, increased primitive progenitor cell frequencies, and reproductive defects. Blood *99*, 111-120. 10.1182/blood.v99.1.111.
16. Dai, X.M., Zong, X.H., Sylvestre, V., and Stanley, E.R. (2004). Incomplete restoration of colony-stimulating factor 1 (CSF-1) function in CSF-1-deficient Csf1op/Csf1op mice by transgenic expression of cell surface CSF-1. Blood *103*, 1114-1123. 10.1182/blood-2003-08-2739.
17. Erblich, B., Zhu, L., Etgen, A.M., Dobrenis, K., and Pollard, J.W. (2011). Absence of colony stimulation factor-1 receptor results in loss of microglia, disrupted brain development and olfactory deficits. PLoS One *6*, e26317. 10.1371/journal.pone.0026317.
18. Wei, S., Nandi, S., Chitu, V., Yeung, Y.G., Yu, W., Huang, M., Williams, L.T., Lin, H., and Stanley, E.R. (2010). Functional overlap but differential expression of CSF-1 and IL-34 in their CSF-1 receptor-mediated regulation of myeloid cells. J Leukoc Biol *88*, 495-505. 10.1189/jlb.1209822.
19. Nandi, S., Gokhan, S., Dai, X.M., Wei, S., Enikolopov, G., Lin, H., Mehler, M.F., and Stanley, E.R. (2012). The CSF-1 receptor ligands IL-34 and CSF-1 exhibit distinct developmental brain expression patterns and regulate neural progenitor cell maintenance and maturation. Dev Biol *367*, 100-113. 10.1016/j.ydbio.2012.03.026.
20. Ginhoux, F., Greter, M., Leboeuf, M., Nandi, S., See, P., Gokhan, S., Mehler, M.F., Conway, S.J., Ng, L.G., Stanley, E.R., et al. (2010). Fate mapping analysis reveals that adult microglia derive from primitive macrophages. Science *330*, 841-845. 10.1126/science.1194637.
21. Miwa, T., Rengasamy, G., Liu, Z., Ginhoux, F., and Okano, T. (2024). Contribution of circulating monocytes in maintaining homeostasis of resident macrophages in postnatal and young adult mouse cochlea. Sci Rep *14*, 62. 10.1038/s41598-023-50634-y.
22. Kishimoto, I., Okano, T., Nishimura, K., Motohashi, T., and Omori, K. (2019). Early Development of Resident Macrophages in the Mouse Cochlea Depends on Yolk Sac Hematopoiesis. Front Neurol *10*, 1115. 10.3389/fneur.2019.01115.
23. Dai, X.-M., Zong, X.-H., Akhter, M.P., and Stanley, E.R. (2004). Osteoclast Deficiency Results in Disorganized Matrix, Reduced Mineralization, and Abnormal Osteoblast Behavior in Developing Bone. Journal of Bone and Mineral Research *19*, 1441-1451. https://doi.org/10.1359/JBMR.040514.
24. Chitu, V., Gokhan, Ş., Nandi, S., Mehler, M.F., and Stanley, E.R. (2016). Emerging Roles for CSF-1 Receptor and its Ligands in the Nervous System. Trends in Neurosciences *39*, 378-393. https://doi.org/10.1016/j.tins.2016.03.005.
25. Bennett, F.C., Bennett, M.L., Yaqoob, F., Mulinyawe, S.B., Grant, G.A., Hayden Gephart, M., Plowey, E.D., and Barres, B.A. (2018). A Combination of Ontogeny and CNS Environment Establishes Microglial Identity. Neuron *98*, 1170-1183 e1178. 10.1016/j.neuron.2018.05.014.

26. Van Wesenbeeck, L., Odgren, P.R., MacKay, C.A., D'Angelo, M., Safadi, F.F., Popoff, S.N., Van Hul, W., and Marks, S.C., Jr. (2002). The osteopetrotic mutation toothless (tl) is a loss-of-function frameshift mutation in the rat Csf1 gene: Evidence of a crucial role for CSF-1 in osteoclastogenesis and endochondral ossification. Proc Natl Acad Sci U S A *99*, 14303-14308. 10.1073/pnas.202332999.
27. Odgren, P.R., Kim, N., van Wesenbeeck, L., MacKay, C., Mason-Savas, A., Safadi, F.F., Popoff, S.N., Lengner, C., van-Hul, W., Choi, Y., and Marks, S.C., Jr. (2001). Evidence that the rat osteopetrotic mutation toothless (tl) is not in the TNFSF11 (TRANCE, RANKL, ODF, OPGL) gene. Int J Dev Biol *45*, 853-859.
28. Marks, S.C., Jr., Mackay, C.A., Jackson, M.E., Larson, E.K., Cielinski, M.J., Stanley, E.R., and Aukerman, S.L. (1993). The skeletal effects of colony-stimulating factor-1 in toothless (osteopetrotic) rats: persistent metaphyseal sclerosis and the failure to restore subepiphyseal osteoclasts. Bone *14*, 675-680. 10.1016/8756-3282(93)90091-n.
29. Marks, S.C., Jr., Iizuka, T., MacKay, C.A., Mason-Savas, A., and Cielinski, M.J. (1997). The effects of colony-stimulating factor-1 on the number and ultrastructure of osteoclasts in toothless (tl) rats and osteopetrotic (op) mice. Tissue Cell *29*, 589-595. 10.1016/s0040-8166(97)80059-6.
30. Joseph, B.K., Marks, S.C., Jr., Hume, D.A., Waters, M.J., and Symons, A.L. (1999). Insulin-like growth factor-I (IGF-I) and IGF-I receptor (IGF-IR) immunoreactivity in normal and osteopetrotic (toothless, tl/tl) rat tibia. Growth Factors *16*, 279-291. 10.3109/08977199909069146.
31. Marks, S.C., Jr., Wojtowicz, A., Szperl, M., Urbanowska, E., MacKay, C.A., Wiktor-Jedrzejczak, W., Stanley, E.R., and Aukerman, S.L. (1992). Administration of colony stimulating factor-1 corrects some macrophage, dental, and skeletal defects in an osteopetrotic mutation (toothless, tl) in the rat. Bone *13*, 89-93. 10.1016/8756-3282(92)90365-4.
32. Yang, M., Mailhot, G., MacKay, C.A., Mason-Savas, A., Aubin, J., and Odgren, P.R. (2006). Chemokine and chemokine receptor expression during colony stimulating factor-1-induced osteoclast differentiation in the toothless osteopetrotic rat: a key role for CCL9 (MIP-1gamma) in osteoclastogenesis in vivo and in vitro. Blood *107*, 2262-2270. 10.1182/blood-2005-08-3365.
33. Marks, S.C., Lundmark, C., Wurtz, T., Odgren, P.R., MacKay, C.A., Mason-Savas, A., and Popoff, S.N. (1999). Facial development and type III collagen RNA expression: Concurrent repression in the osteopetrotic(Toothless, tl) rat and rescue after treatment with colony-stimulating factor-1. Developmental Dynamics *215*, 117-125. 10.1002/(sici)1097-0177(199906)215:2<117::Aid-dvdy4>3.0.Co;2-d.
34. Aharinejad, S., Grossschmidt, K., Franz, P., Streicher, J., Nourani, F., MacKay, C.A., Firbas, W., Plenk, H., Jr., and Marks, S.C., Jr. (1999). Auditory ossicle abnormalities and hearing loss in the toothless (osteopetrotic) mutation in the rat and their improvement after treatment with colony-stimulating factor-1. J Bone Miner Res *14*, 415-423. 10.1359/jbmr.1999.14.3.415.
35. Wisner-Lynch, L.A., Shalhoub, V., and Marks, S.C., Jr. (1995). Administration of colony stimulating factor-1 to toothless osteopetrotic rats normalizes osteoblast, but not osteoclast, gene expression. Bone *16*, 611-618. 10.1016/8756-3282(95)00114-s.
36. Cotton, W.R., and Gaines, J.F. (1974). Unerupted dentition secondary to congenital osteopetrosis in the Osborne-Mendel rat. Proc Soc Exp Biol Med *146*, 554-561. 10.3181/00379727-146-38146.

37. Dobbins, D.E., Sood, R., Hashiramoto, A., Hansen, C.T., Wilder, R.L., and Remmers, E.F. (2002). Mutation of macrophage colony stimulating factor (Csf1) causes osteopetrosis in the tl rat. Biochem Biophys Res Commun *294*, 1114-1120. 10.1016/S0006-291X(02)00598-3.
38. Batoon, L., Keshvari, S., Irvine, K.M., Ho, E., Caruso, M., Patkar, O.L., Sehgal, A., Millard, S.M., Hume, D.A., and Pettit, A.R. (2024). Relative contributions of osteal macrophages and osteoclasts to postnatal bone development in CSF1R-deficient rats and phenotype rescue following wild-type bone marrow cell transfer. J Leukoc Biol *116*, 753-765. 10.1093/jleuko/qiae077.
39. Hume, D.A., Teakle, N., Keshvari, S., and Irvine, K.M. (2023). Macrophage deficiency in CSF1R-knockout rat embryos does not compromise placental or embryo development. J Leukoc Biol *114*, 421-433. 10.1093/jleuko/qiad052.
40. Keshvari, S., Caruso, M., Teakle, N., Batoon, L., Sehgal, A., Patkar, O.L., Ferrari-Cestari, M., Snell, C.E., Chen, C., Stevenson, A., et al. (2021). CSF1R-dependent macrophages control postnatal somatic growth and organ maturation. PLoS Genet *17*, e1009605. 10.1371/journal.pgen.1009605.
41. Patkar, O.L., Caruso, M., Teakle, N., Keshvari, S., Bush, S.J., Pridans, C., Belmer, A., Summers, K.M., Irvine, K.M., and Hume, D.A. (2021). Analysis of homozygous and heterozygous Csf1r knockout in the rat as a model for understanding microglial function in brain development and the impacts of human CSF1R mutations. Neurobiol Dis *151*, 105268. 10.1016/j.nbd.2021.105268.
42. Carter-Cusack, D., Huang, S., Keshvari, S., Patkar, O., Sehgal, A., Allavena, R., Byrne, R.A.J., Morgan, B.P., Bush, S.J., Summers, K.M., et al. (2025). Wild-type bone marrow cells repopulate tissue resident macrophages and reverse the impacts of homozygous CSF1R mutation. PLoS Genet *21*, e1011525. 10.1371/journal.pgen.1011525.
43. Hume, D.A., Batoon, L., Sehgal, A., Keshvari, S., and Irvine, K.M. (2022). CSF1R as a Therapeutic Target in Bone Diseases: Obvious but Not so Simple. Curr Osteoporos Rep *20*, 516-531. 10.1007/s11914-022-00757-4.
44. Hume, D.A., Caruso, M., Ferrari-Cestari, M., Summers, K.M., Pridans, C., and Irvine, K.M. (2020). Phenotypic impacts of CSF1R deficiencies in humans and model organisms. Journal of Leukocyte Biology *107*, 205-219. https://doi.org/10.1002/JLB.MR0519-143R.
45. Pridans, C., Raper, A., Davis, G.M., Alves, J., Sauter, K.A., Lefevre, L., Regan, T., Meek, S., Sutherland, L., Thomson, A.J., et al. (2018). Pleiotropic Impacts of Macrophage and Microglial Deficiency on Development in Rats with Targeted Mutation of the Csf1r Locus. J Immunol *201*, 2683-2699. 10.4049/jimmunol.1701783.
46. Lawrence, A.R., Canzi, A., Bridlance, C., Olivie, N., Lansonneur, C., Catale, C., Pizzamiglio, L., Kloeckner, B., Silvin, A., Munro, D.A.D., et al. (2024). Microglia maintain structural integrity during fetal brain morphogenesis. Cell *187*, 962-980 e919. 10.1016/j.cell.2024.01.012.
47. McNamara, N.B., Munro, D.A.D., Bestard-Cuche, N., Uyeda, A., Bogie, J.F.J., Hoffmann, A., Holloway, R.K., Molina-Gonzalez, I., Askew, K.E., Mitchell, S., et al. (2023). Microglia regulate central nervous system myelin growth and integrity. Nature *613*, 120-129. 10.1038/s41586-022-05534-y.
48. Rojo, R., Raper, A., Ozdemir, D.D., Lefevre, L., Grabert, K., Wollscheid-Lengeling, E., Bradford, B., Caruso, M., Gazova, I., Sanchez, A., et al. (2019). Deletion of a Csf1r enhancer selectively impacts CSF1R expression and development of tissue macrophage populations. Nat Commun *10*, 3215. 10.1038/s41467-019-11053-8.

49. Munro, D.A.D., Bradford, B.M., Mariani, S.A., Hampton, D.W., Vink, C.S., Chandran, S., Hume, D.A., Pridans, C., and Priller, J. (2020). CNS macrophages differentially rely on an intronic Csf1r enhancer for their development. Development *147*. 10.1242/dev.194449.
50. Kiani Shabestari, S., Morabito, S., Danhash, E.P., McQuade, A., Sanchez, J.R., Miyoshi, E., Chadarevian, J.P., Claes, C., Coburn, M.A., Hasselmann, J., et al. (2022). Absence of microglia promotes diverse pathologies and early lethality in Alzheimer's disease mice. Cell Rep *39*, 110961. 10.1016/j.celrep.2022.110961.
51. Munro, D.A.D., Bestard-Cuche, N., McQuaid, C., Chagnot, A., Shabestari, S.K., Chadarevian, J.P., Maheshwari, U., Szymkowiak, S., Morris, K., Mohammad, M., et al. (2024). Microglia protect against age-associated brain pathologies. Neuron *112*, 2732-2748.e2738. 10.1016/j.neuron.2024.05.018.
52. Nagahama, S.I., Cunningham, M.L., Lee, M.Y., and Byers, M.R. (1998). Normal development of dental innervation and nerve/tissue interactions in the colony-stimulating factor-1 deficient osteopetrotic mouse. Developmental Dynamics *211*, 52-59. 10.1002/(sici)1097-0177(199801)211:1<52::Aid-aja5>3.0.Co;2-6.
53. Cohen, P.E., Zhu, L., Nishimura, K., and Pollard, J.W. (2002). Colony-stimulating factor 1 regulation of neuroendocrine pathways that control gonadal function in mice. Endocrinology *143*, 1413-1422. 10.1210/endo.143.4.8754.
54. Pollard, J.W., and Hennighausen, L. (1994). Colony stimulating factor 1 is required for mammary gland development during pregnancy. Proc Natl Acad Sci U S A *91*, 9312-9316. 10.1073/pnas.91.20.9312.
55. McFarlane, L., Truong, V., Palmer, J.S., and Wilhelm, D. (2013). Novel PCR assay for determining the genetic sex of mice. Sex Dev *7*, 207-211. 10.1159/000348677.
56. Tiscornia, G., Singer, O., Ikawa, M., and Verma, I.M. (2003). A general method for gene knockdown in mice by using lentiviral vectors expressing small interfering RNA. Proc Natl Acad Sci U S A *100*, 1844-1848. 10.1073/pnas.0437912100.
57. The Jackson Laboratory. Protocol 25394: Standard PCR Assay - Tg(Wnt1-cre). https://www.jax.org/Protocol?stockNumber=022501&protocolID=25394.
58. The Jackson Laboratory. Protocol 29436: Standard PCR Assay - Gt(ROSA)26Sor(tdTomato-WPRE). https://www.jax.org/Protocol?stockNumber=007914&protocolID=29436.
